# Revealing the evolutionary history and contemporary population structure of Pacific salmon in the Fraser River through genome resequencing

**DOI:** 10.1101/2024.04.29.591761

**Authors:** Kris A. Christensen, Anne-Marie Flores, Dionne Sakhrani, Carlo A. Biagi, Robert H. Devlin, Ben J. G. Sutherland, Ruth E. Withler, Eric B. Rondeau, Ben F. Koop

## Abstract

The Fraser River once supported massive salmon returns, but now years with half of the recorded historical maximum are considered good. There is substantial interest from surrounding communities, governments, and other groups to increase salmon returns for both human use and for functional ecosystems. To help generate resources for this endeavour, we resequenced hundreds of genomes at moderate coverage (∼16x) of Chinook (*Oncorhynchus tshawytscha*), coho (*O. kisutch*), and sockeye salmon (*O. nerka*) from the Fraser River. The resequenced genomes are an important resource that can give us new insights. In this study, we found evidence that Chinook salmon have 1.5-2x more polymorphic loci than coho or sockeye salmon. Using principal component analysis (PCA) and admixture analysis, we also identified genetic groups similar to those previously identified with only a few microsatellite markers. As the higher density data supports these previous genetic groups, it suggests that the identity of these groups is not overly sensitive to the number of genetic markers or when the groups were sampled. With the increased resolution from resequenced genomes, we were able to further identify factors influencing these genetic groups, including isolation-by-distance, migration barriers, recolonization from different glacial refugia, and environmental factors like precipitation. We were also able to identify 20 potentially adaptive loci among the genetic groups by analyzing runs of homozygosity. All of the resequenced genomes have been submitted to a public database where they can be used as a reference for the contemporary genomics of Fraser River salmon.

**Article Summary:** Concerns over Fraser River salmon declines led us to generate resources to better understand the genetics of these salmon. Our findings were similar to previous studies that examined a very small fraction of the genetic markers examined here. We expanded upon previous studies by identifying possible influences on the genetics of Fraser River salmon. These included environmental factors, historical differences, and a lack of connectivity in the river caused by the Fraser Canyon.

## Introduction

The Fraser River in Canada may have existed in some form for over 66 million years (Tribe 2005). The modern flow of the river is much more recent as there is evidence of a major change in flow around 760,000 years ago (Andrews *et al*. 2012). During the last glaciation period (the last in the Pleistocene), the entire region where the modern river is located may have been covered by the Cordilleran Ice Sheet, which reached its estimated maximum around 19,000 years ago (Clark *et al*. 2009). The retreat of the Cordilleran Ice Sheet began as early as 18,000 years ago and there were ice-free regions in British Columbia (where the Fraser River is located) as early as 17,700 years ago (Warner *et al*. 1982; Darvill *et al*. 2018). Some regions of British Columbia may have even been ice-free during the glacial maximum (e.g., (Byun *et al*. 1999; McPhail 2007; Stewart *et al*. 2009)). The Cordilleran Ice Sheet fully retreated by around 11,000 years ago, and its only remnants are the glaciers that still exist in modern times (Clague 2017). These massive changes in the Fraser River environment likely caused local extinctions of plants and animals from the river.

Based on fossil evidence, there were salmonids in the Fraser River before it was covered by glaciers (Harington 1996). From paleogeographic evidence mentioned above and this fossil evidence, we can infer that modern salmon in the river must have re-colonized after glaciers receded. There is similar evidence of colonization of other species to this region, such as different plants, deer, and other ray-finned fishes (Soltis *et al*. 1997; Bernatchez and Wilson 1998; Hewitt 2000; McPhail 2007; Latch *et al*. 2009; Beatty and Provan 2010). Colonization or re-colonization of the 65-66 species of ray-finned fishes native to British Columbia was thought to have started around 13,000 years ago (McPhail 2007). However, fossil evidence indicates that kokanee (*O. nerka*) could have re-colonized the interior of British Columbia, possibly via the Columbia River, as early as 18,000 years ago (Harington 1996). This is consistent with a large isolated and unique kokanee population in the upper Columbia River and Fraser River system that remains today (Beacham and Withler 2017; Christensen *et al*. 2020).

While there is a large body of paleogeographic research and a fossil record describing glaciation and species colonization of the Fraser River after glaciers retreated, we often do not have information on specific locations or populations. We can turn to genetic studies to complement what we understand from these other two fields of study. We expect to observe genetic legacies of re-colonization events in modern populations (Hewitt 1996, 2000). These legacies may influence modern populations, and understanding them will be important for evidence-based conservation and management. Several researchers have suggested that Fraser River salmon have experienced recent genetic bottlenecks that could be the outcome of re-colonization events (Wehrhahn and Powell 1987; Wood *et al*. 1994; Rondeau *et al*. 2023).

Generally, there were two to three Fraser River (e.g., lower, middle, and upper) Chinook (*O. tshawytscha*), sockeye (*O. nerka*), or coho salmon (*O. kisutch*) genetic groups identified from previous studies (Wood *et al*. 1994; Small *et al*. 1998; Teel *et al*. 2000; Beacham *et al*. 2003; Beacham and Withler 2017). This excludes additional groups identified from the Thompson River tributary that were also commonly identified (Small *et al*. 1998; Withler *et al*. 2000; Beacham *et al*. 2003; Beacham and Withler 2017). While these different genetic groups could have originated from different re-colonization events or from different combinations of re-colonization events, they could also have originated as a result of other factors. These factors include: limited gene flow between the groups (e.g., due to barriers like the Fraser Canyon, a phenomenon sometimes referred to as isolation-by-resistance (McRae 2006)), because of stochastic mechanisms like isolation-by-distance, adaptation to environmental conditions (sometimes referred to as isolation-by-environment (Weber *et al*. 2017)), ecotype differentiation in sockeye salmon (Beacham and Withler 2017), or through a combination of these mechanisms. If these genetic groups were a result of different re-colonization events, it is also possible that the re-colonizing populations came from different glacial refugia (Wood *et al*. 1994; Small *et al*. 1998; Teel *et al*. 2000; Withler *et al*. 2000), which would increase their overall distinctiveness.

Wehrhahn and Powell (1987) note that the Fraser Canyon might limit gene flow between lower and upper Fraser River salmon. There are several identified velocity barriers in the Fraser Canyon that could influence salmon migration and particularly limit migration of smaller salmon (Wright 2022).

This was dramatically emphasized between 1913-1914 when construction left a massive rockslide in the Fraser Canyon that partially blocked some salmon species, and completely blocked pink salmon (*O. gorbuscha*) (Kew 1992; Grant and Pestal 2009; Pess *et al*. 2012). This single event reduced salmon returns to around a quarter of historic returns to the Fraser River during peak years (Pacific Salmon Commission – psc.org). If the Fraser Canyon acts as a barrier, we might expect unique genetic characteristics above and below the Fraser Canyon for all the species that must traverse it.

Upper and lower Fraser River salmon are also separated by large distances and because they have natal homing with variable straying (reviewed in (Quinn 1993; Keefer and Caudill 2013; Bett *et al*. 2017)), we expect population structure to be influenced by isolation-by-distance (e.g., (Wright 1943; Weber *et al*. 2017; Aguillon *et al*. 2017)). In general, we would expect isolation-by-distance to increase the longer temporally that salmon are isolated from each other and with increased distance. We also expect to observe nearby groups to have higher genetic similarity than those further away for all species. An exception to this expectation would be salmon with different life history types (e.g., odd and even-year pink salmon (Christensen *et al*. 2021)).

The expectation of increased genetic distance with increased geographic distance will not be the main driver of differentiation if genetic groups were largely influenced by separate re-colonization events. If re-colonization occurred by different genetic groups (e.g., populations from different glacial refugia), we might expect genetic distance to be unrelated to geographic distance in the absence of strong gene-flow or long periods of time. Patterns of re-colonization from different genetic groups would also be expected to be variable among species.

Adaptation to the environment is of particular interest for the conservation and management of a species because salmon from an area could be uniquely suited to that region. The headwaters of the Fraser River flow north down the Rocky Mountain Trench, which leads to the Interior Plateau, followed by the Fraser Canyon, and finally the river outlets through a major delta in the metropolis of Vancouver, British Columbia (Reynoldson *et al*. 2005). These different regions have variable climates, elevations, and river velocities (Reynoldson *et al*. 2005). They are also separated by great distances.

Genetic signatures correlated with environmental factors could be from neutral factors such as isolation-by-distance or from non-neutral factors such as local adaptation or other forms of selection. Researchers have not yet found a reliable method for distinguishing among the different mechanisms with genetic data alone (reviewed in (Bierne *et al*. 2013; Lotterhos and Whitlock 2015; Ahrens *et al*. 2018; Saravanan *et al*. 2020)), and other experiments need to be used to firmly establish particular genetic adaptations.

In this study, we analyze 954 resequenced genomes from three species of salmon (317 sockeye, 360 Chinook, and 277 coho salmon), mainly from the Fraser River, to understand differences among the species, identify genetic groups among the collections of samples within each species, and to characterize genetic adaptation to different environments. Data related to genetic structure and local adaptation are particularly important to conservation and management. By using data from multiple species, we compare among species to identify common patterns. This work provides a foundational dataset that will be openly available and of significant use to the salmonid research community for years to come.

## Materials and Methods

### Sampling and whole genome resequencing

Chinook, coho, and sockeye salmon were sampled from Fraser River tributaries and lakes or were from other bodies of water from previous studies (Figure 1, Figure S1, File S1, (Christensen *et al*. 2020; Rondeau *et al*. 2023)). Some of these sampling locations include hatchery sources (File S1).

**Figure 1.**
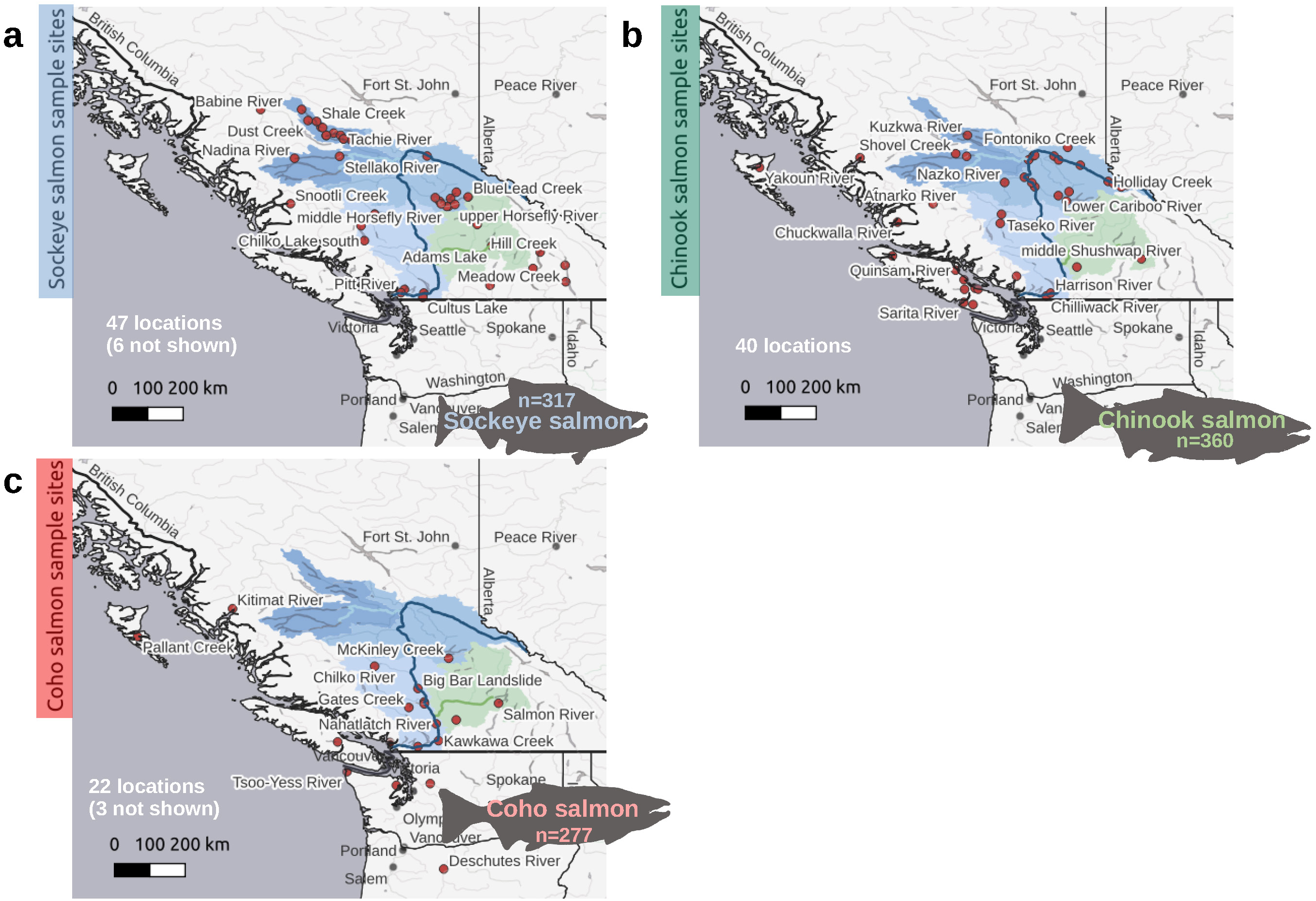
Fraser River salmon sampling locations. a) Each point on the map represents a sockeye salmon sampling site (average of 6.5 samples per location). Each location is shown as the closest point of the specified body of water to the Fraser River. The Fraser River and associated watersheds are highlighted on the map. The Nechako River is highlighted as the top left watershed (darker blue, with the river as a lighter blue), and the Thompson River is highlighted as the bottom right watershed (green). Watershed and geographical data are from ced.org and naturalearthdata.com, respectively. The map was generated using QGIS software. Information on sampling locations outside of the boundaries of this map can be found in File S1. b) Same as a) for Chinook salmon samples (average of 7.8 samples per site). c) Same as a) for coho salmon samples (average of 6.9 samples per site – excluding the Big Bar Landslide sampling site).

Most modern hatchery stocks originate from local sources (e.g. (Heard 2012)), and they should reflect the current Fraser River population if not the historical population. The samples were collected during various years (File S1). For coho salmon, there were 125 salmon collected at the Big Bar Landslide between 2019-2021. Spawning locations were unknown for these samples. These were included to identify potentially unknown genetic structure since it is difficult to reach upper Fraser River sites during coho salmon spawning.

Samples were collected by Fisheries and Oceans Canada personnel in compliance with the Canadian Council on Animal Care Guidelines, and under the authority of the Fisheries and Oceans Canada Pacific Region Animal Care Committee (Ex.7.1). Samples were taken either as operculum-clips or as scales. Both were desiccated and stored on Whatman paper.

Genomic DNA was extracted from tissue, after an overnight incubation in 95% ethanol, using a Quick-DNA Kit (Zymo Research). Other genomic DNA samples were previously extracted by Fisheries and Oceans Canada using automated BioSprint extractions, as per manufacturer’s instructions (Qiagen). DNA samples were then sent to the Michael Smith Genome Science Centre (Vancouver, BC) for library preparation and whole genome resequencing.

Whole genome sequencing libraries were prepared by shearing the DNA samples individually using a Covaris LE220 (duty cycle: 20%, PIP: 450, cycles per burst: 200, time per run: 90 s; with pulse spin after 45 s). Individual libraries were then constructed using the MGIEasy PCR-Free DNA Library Prep Set (MGI Tech Co.). The indexed libraries were then pooled and sequenced on a MGISeq-G400 sequencer (paired-end 200 bp). Data used in this study were also taken from previous studies (Christensen *et al*. 2020; Rondeau *et al*. 2023) that used Illumina sequencing technology.

### SNP calling and filtering

Reads from each individual were aligned to their respective species reference genome assembly (Chinook salmon: GCF_018296145.1 (Christensen *et al*. 2018), coho salmon: GCF_002021735.2 (Rondeau *et al*. 2023), sockeye salmon: unreleased version 2 (Christensen *et al*. 2020)) with BWA (version 0.7.17, parameter -M) (Li and Durbin 2009, 2010; Li 2013). Reads were sorted with SAMtools (version 1.12, default parameters) (Danecek *et al*. 2021). Picard (version 2.26.3, default parameters) (Broad Institute 2019) was used to add read group information and mark reads that were suspected PCR duplicates. GATK (version 3.8) (McKenna *et al*. 2010; Van der Auwera *et al*. 2013) was then used to call nucleotide variants for each individual (parameters: -T HaplotypeCaller, --genotyping_mode DISCOVERY, --emitRefConfidence GVCF) and then combined (parameters: -T GenotypeGVCFs, --max_alternate_alleles 3). Truth and training SNP datasets (File S2) were then used to recalibrate nucleotide variant scores and filter variants using GATK (parameters -T ApplyRecalibration, --mode SNP –ts_filter_level 99.5). The truth SNPs came from multiple studies (Brieuc *et al*. 2014; Meek *et al*. 2016; Nichols *et al*. 2016; Larson *et al*. 2017; Veale and Russello 2017; Rondeau *et al*. 2023), and the training SNPs are described in File S2.

Additional filters were used to remove indels, SNPs with more or fewer than two alleles, or that were missing genotypes in more than 10% of the individuals. SNPs were removed if their mean depths were outside a range of 8-100x, and if they had less than 0.01 minor allele frequency (MAF). All filtering was performed using VCFtools (version 0.1.15) (Danecek *et al*. 2011). The MAF would eliminate alleles found in fewer than 6-8 heterozygous individuals (or 3-4 homozygous individuals), depending on the species and its respective sample size. This threshold was chosen to reduce sequencing errors, but to still keep all but the rarest variants. The MAF filter was not used for the SMC++ analysis. Linkage disequilibrium was evaluated and used to filter SNPs in some analyses, as noted below. The prune add-on for BCFtools (version 1.9) (Danecek *et al*. 2021) was used to filter based on linkage disequilibrium (parameters: +prune, -w 20kb, -l 0.4, -n 2). The filters used for all analyses are shown in Table 1.

**Table 1.** SNP filtering parameters used for each analysis.

| Analyses | Indels, >10% missing, depth | MAF 0.01 | LD Filter | 15x coverage | Relatedness |
| --- | --- | --- | --- | --- | --- |
| SMC++ | all |  |  |  |  |
| Rehh | all | all |  |  |  |
| Coverage | all | all |  |  |  |
| ROH | all | all |  |  |  |
| Relatedness | all | all | all |  |  |
| Admixture | all | all | all |  |  |
| % Heterozygous | all | all |  | all |  |
| Nucleotide diversity | all | all |  | all |  |
| Polymorphic loci | all | all |  | all |  |
| PCA | all | all | all | some* | some* |

\*See Figure S4 for all PCAs, including some with the 15x SNP coverage and relatedness filters.

### Mapping variants among species

To compare analyses among species, variants were mapped from the coho and sockeye salmon genomes to the Chinook salmon genome using a pipeline, *MapVCF2NewGenome* (see *Data Availability*). This pipeline was also used to map sockeye salmon variants, which were on scaffolds, to a chromosome level assembly (this assembly was submitted to the NCBI as version 2 of the reference genome assembly – now available as GCA_034236695.1). The sockeye salmon chromosome mapped version was needed for several analyses.

### Coverage, relatedness metrics, and runs of homozygosity (ROH)

Coverage was assessed using a python script, *VCFStats* (see *Data Availability*). This script finds the average depth of all SNPs per individual. It also counts the number and type of genotypes per individual. This information was used to remove individuals, using VCFtools, that had average depth/coverage less than 8x, as a preliminary PCA analysis showed clustering of individuals based on coverage rather than geography if they were below this threshold. These individuals are not discussed and were not analyzed further. Individuals were filtered with an average depth less than 15x depending on the analysis (noted for each analysis). This was because the frequency of heterozygous genotypes dropped when coverage was below 15x and some analyses were sensitive to this issue (Table 1). Genotype information and coverage was visualized in R (R Core Team 2022) using ggplot2 (Wickham 2016).

The related package (version 0.8, parameters of coancestry: ritland = 1) (Pew *et al*. 2015) was used to calculate relatedness (Ritland 1996) in R. Values below zero (e.g., when markers are not shared) and above one (e.g., when rare markers are shared) are expected with this type of estimation (Ritland 1996). There were many more private alleles in coastal populations of Chinook and coho salmon, which may have made them appear much more related. The linkage disequilibrium filtered SNP datasets were converted to the related format using a python script, *VCF2Relate* (see *Data Availability*). The reshape2 (Wickham 2007) and pheatmap (Kolde 2019) packages were used to visualize relatedness. Related individuals were included in most analyses, but we tested what would happen if related individuals were removed from PCA analyses. A relatedness score of 0.15 (an arbitrary value greater than what might be expected for first cousins, but less than for half-sibs – depending on dataset) was used to filter all but one individual from pairs that might be related. Most samples were taken as adults and related individuals were not expected to be common within or among sampling collections.

Runs of homozygosity were identified using PLINK (version 1.9, parameters: --homozyg – double-id, --allow-extra-chr, --homozyg-snp 25, --homozyg-kb, --homozyg-density, --homozyg-gap, --homozyg-window-het 1, --homozyg-window-snp 50, --homozyg-window-threshold 0.05, --homozyg-window-missing 5) (Chang *et al*. 2015; “PLINK 1.9”). This was to compare species levels of runs of homozygosity. All individuals were included in these analyses. No consistent relationship was observed between coverage and runs of homozygosity (e.g., runs of homozygosity might be expected to increase as coverage decreases since there are fewer heterozygous genotypes as noted above). We plotted each individual when comparing runs of homozygosity among species to determine if coverage or relatedness might influence species comparisons.

### Population structure and clustering analyses

PCA and admixture analyses were used to identify distinct genetic groups for each species. The admixture software requires independent variants, and PCAs can be sensitive to LD blocks, which can cause clustering that is unrelated to population structure. For this reason, the LD filtered SNPs were used in these analyses. To perform the admixture analyses, the LD filtered data was converted from VCF format to a suitable format using PLINK (version 1.9, parameters: --double-id, --allow-extra-chr). We converted the chromosome names to numbers using Unix commands. The admixture analysis was performed using ADMIXTURE (version 1.3.0, parameters: --cv) (Alexander *et al*. 2009). Cluster values 1-20 were tested and we accepted the value with the lowest cross-validation score. Admixture ancestry values were visualized in R for individuals and in QGIS (QGIS Development Team 2022) as an average score per sampling site.

PCA were used to assess the groupings produced from the admixture analyses. PLINK (parameters: --pca, --double-id, --allow-extra-chr) was used to perform the PCA, and they were visualized in R using ggplot2, reshape2, and ggrepel (Slowikowski 2021). PCA were tried with all the individuals, individuals with ≥15x coverage, and with individuals that were highly related removed.

The outputs of these three approaches were compared to determine the influence of coverage and relatedness on clustering and the groups produced from the admixture analyses. We also verified that samples among the new and previous datasets clustered together if they were from the same sampling site.

### Environmental variable PCA

To determine whether population structure was associated with environmental factors, we clustered sampling sites by environmental factors using PCA in R (parameters: prcomp, scale=T). Each sampling site was assigned to an admixture group if the average ancestry value was ≥ 0.7 for a particular site. Environmental factors were taken from the WorldClim version 2.1 dataset (Fick and Hijmans 2017). Elevation for each site was either estimated from Google maps, mapcarta, or from data downloaded from the Federal Geospatial Platform (maps.canada.ca) and viewed in QGIS. Distance to the ocean was estimated as the river distance with QGIS or Google maps.

### Historical estimates of effective population size

To estimate effective population size through time, we used the program SMC++ (parameters: - c 1000000) (Terhorst *et al*. 2017). The only mutation rate estimate for these species available at the time of writing was for coho salmon (8e-9 from (Rougemont *et al*. 2020)). A correction was applied to this mutation rate for Chinook and sockeye salmon based on the ratio of total SNPs from these species to the total SNPs from coho salmon. For Chinook salmon, this ratio was 1.7274 for the individuals subset from a similar geographic range and for the same number of individuals (mutation rate of 1.382e-8). For sockeye salmon, this was 0.79785342 for the individuals subset, with a mutation rate of 6.38283e-9. Figures were plotted in R using ggplot2, and ocean surface temperature data was taken from previous studies (Zachos *et al*. 2008; Hansen *et al*. 2013).

### Genetic diversity

The percent of heterozygous genotypes per individual was calculated by dividing the number of heterozygous genotypes (see *Coverage, relatedness metrics, and runs of homozygosity* section) by all genotypes (including missing data to standardize against all variants) and multiplying by 100. If missing genotypes were excluded from the calculation, values changed by 0.09% on average, and the difference ranged from 0.03%-1.4%. Nucleotide diversity (Pi) and the number of polymorphic loci was calculated using the Stacks populations module (version 2.54, default parameters) (Catchen *et al*. 2011, 2013).

The percent of heterozygous genotypes per individual and nucleotide diversity per sampling site were both calculated from the subset of individuals with ≥15x coverage because these analyses were sensitive to coverage (i.e., as coverage increased, so did the percent of heterozygous genotypes until ≥15x coverage). The number of sampling sites were included in the reported percent of heterozygous genotypes per individual to show if the reported value could be sensitive to relatedness (e.g., fewer locations could be more sensitive to highly related individuals from a single location). Since the nucleotide diversity was plotted per sampling site using the inverse distance weighting interpolation analysis in QGIS, other nearby sites can also be used to determine if relatedness influenced nucleotide diversity regionally.

### Admixture group private alleles

A python script was used to identify private alleles among admixture groups of each species (*PrivateAllele* see *Data Availability*). Alleles were identified that were unique to each group if they were present for a specified number of individuals in that group (parameter: -min 3). Individuals were assigned to an admixture group if they had ancestry values ≥ 0.7 for a particular group. Salmon with ancestry values below 0.7 were excluded from these analyses. The private allele counts were visualized in R using ggplot and the gridExtra library (Auguie 2017).

The number of private alleles per individual among admixture groups was identified using another python script, *PrivateAllelePerInd* (see *Data Availability*). This metric identifies if there are individuals or sampling sites that were more responsible for the number of private alleles within a genetic group. The minimum number of individuals with the private allele was set to one. While decreasing this parameter to one could increase the chance that SNP calling errors are included, it also allows us to get an idea for the number of private alleles from a group that are within the genome of each individual. We expect sequencing errors to be evenly distributed among sampling locations and similar among individuals.

### Potentially adaptive variants

The rehh library (Gautier and Vitalis 2012; Gautier *et al*. 2017) in R was used to identify extended haplotype homozygosity within and among admixture groups (assignments were made for individuals with ≥ 0.7 ancestry values). To perform the rehh analyses, we used SHAPEIT5 (Hofmeister *et al*. 2023) (version 5.1.1, parameter: phase_common_static) to first phase the genotypes of the MAF 0.01 filtered variants. The output was then converted to VCF format using BCFtools, and the different admixture groups were separated using VCFtools. The rehh command *data2haplohh* was then used to read in each admixture group VCF file, and the *scan_hh* command (parameters, polarized=FALSE, interpolate=FALSE) was used to calculate extended haplotype homozygosity. The within population metric iHS was calculated using the *ihh2ihs* command (default parameters), and the pairwise population metric Rsb was calculated using the *ines2rsb* command (default parameters). The *calc_candidate_regions* function (parameters: threshold=10, window_size=10000, min_n_extr_mrk=8) was used to identify regions with significant differences in extended haplotype homozygosity (the Bonferroni correction p-value threshold was between 8.8-9.13 after -log10 transformation depending on species). A higher threshold was used to identify only the strongest candidates as this is only a preliminary study. We also examined overlapping 10 kbp windows among all three species to identify candidates of potential convergent evolution.

## Results

### Coverage, relatedness metrics, and runs of homozygosity (ROH)

We identified ∼6-13 million SNPs per species (Figure 2a). Chinook salmon had 1.58-2.07x more SNPs than either coho or sockeye salmon, respectively. Between 1-1.75% of all SNP loci were common among species, and only up to 0.1% were found to be polymorphic in all species (Figure 2b). Chinook salmon had a reduction in the total length of runs of homozygosity (ROH), consistent with a higher number of polymorphic loci (Figure 2c). Sockeye and coho salmon had on average ∼3.5-3.6x longer total lengths of ROH than Chinook salmon. The length of ROH decreased as the fraction of heterozygous genotypes increased, but generally sockeye and coho salmon had longer ROH when fractions of heterozygous genotypes were below 0.2 (Figure 2d). No consistent relationship among species was observed between SNP coverage and the total length of ROH (Figure S2).

**Figure 2.**
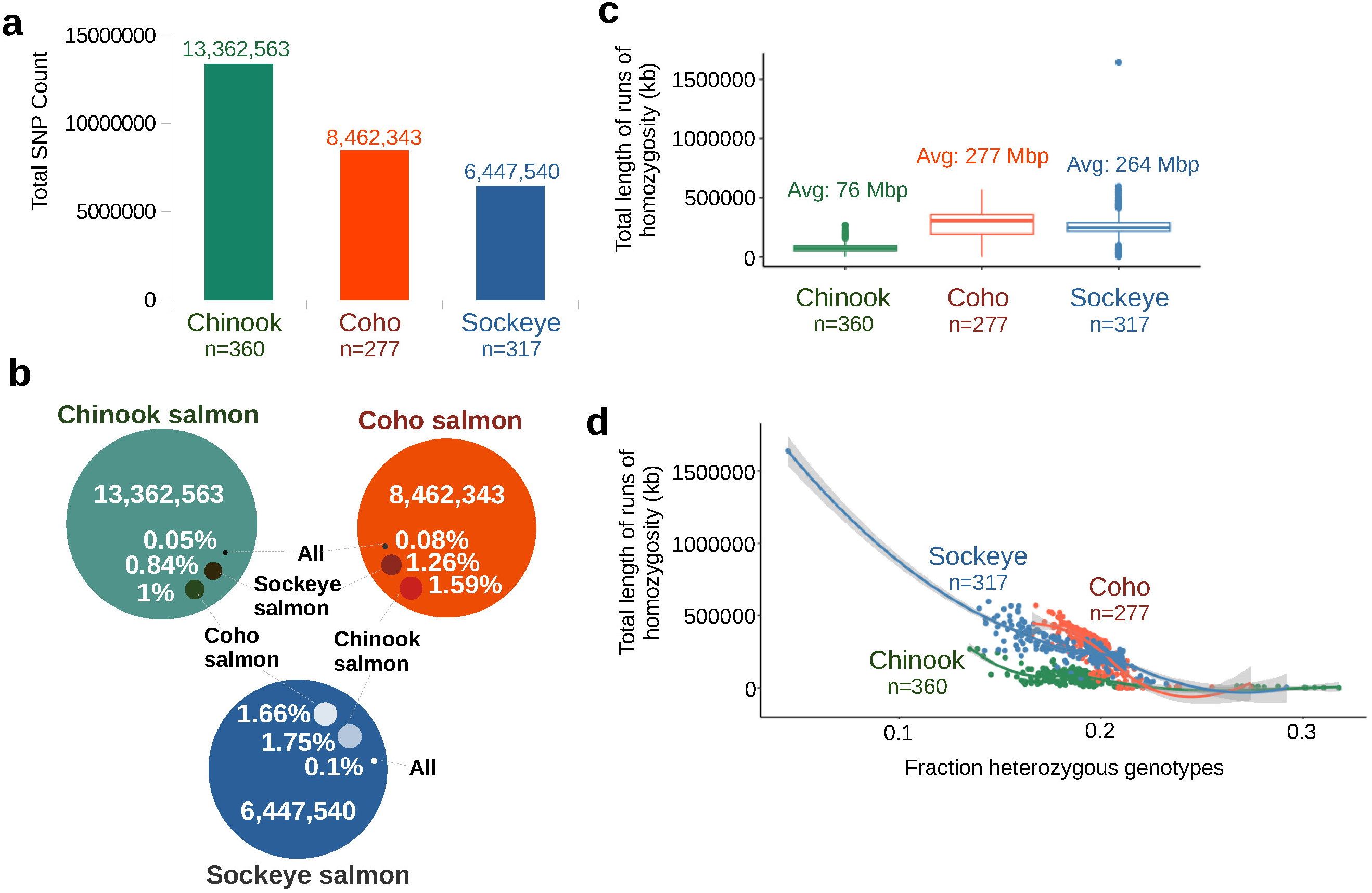
Polymorphic loci per species and runs of homozygosity. a) Counts of all SNPs identified for each species with the same pipeline – differing only in score recalibration and reference genome assemblies. The number of samples per species is shown below each column. b) Overlapping SNP loci among species. SNP loci from sockeye and coho salmon were mapped to the Chinook salmon genome assembly. The percent of loci that overlap are shown for each species relative to the number of SNPs identified in that species. c) Boxplots of the total lengths of runs of homozygosity for each species. d) A scatterplot of runs of homozygosity and the fraction of heterozygous genotypes per individual. The sockeye sample on the far left is a doubled-haploid. Lines were fitted using the loess method.

The average SNP coverage of the combined 954 salmon was 16x (Figure 3a). The average coverage of Chinook salmon was 16.5x (n=360), coho salmon 18.1x (n=277), and sockeye salmon 13.7x (n=317). The percent of heterozygous genotypes for each individual appeared to depend on the SNP coverage until a depth of 15x (Figure 3a). This relationship was consistent in all species (Figure 3b). This issue did not appear to influence the scale of the number of SNPs among species. The average number of heterozygous genotypes per salmon for those with 15x coverage or greater reflected the trend observed for the number of SNPs per species. These values were 1,667,558 in coho salmon, 1,305,776 in sockeye salmon, and 2,636,167 in Chinook salmon (1.58-2.02x greater in Chinook salmon).

**Figure 3.**
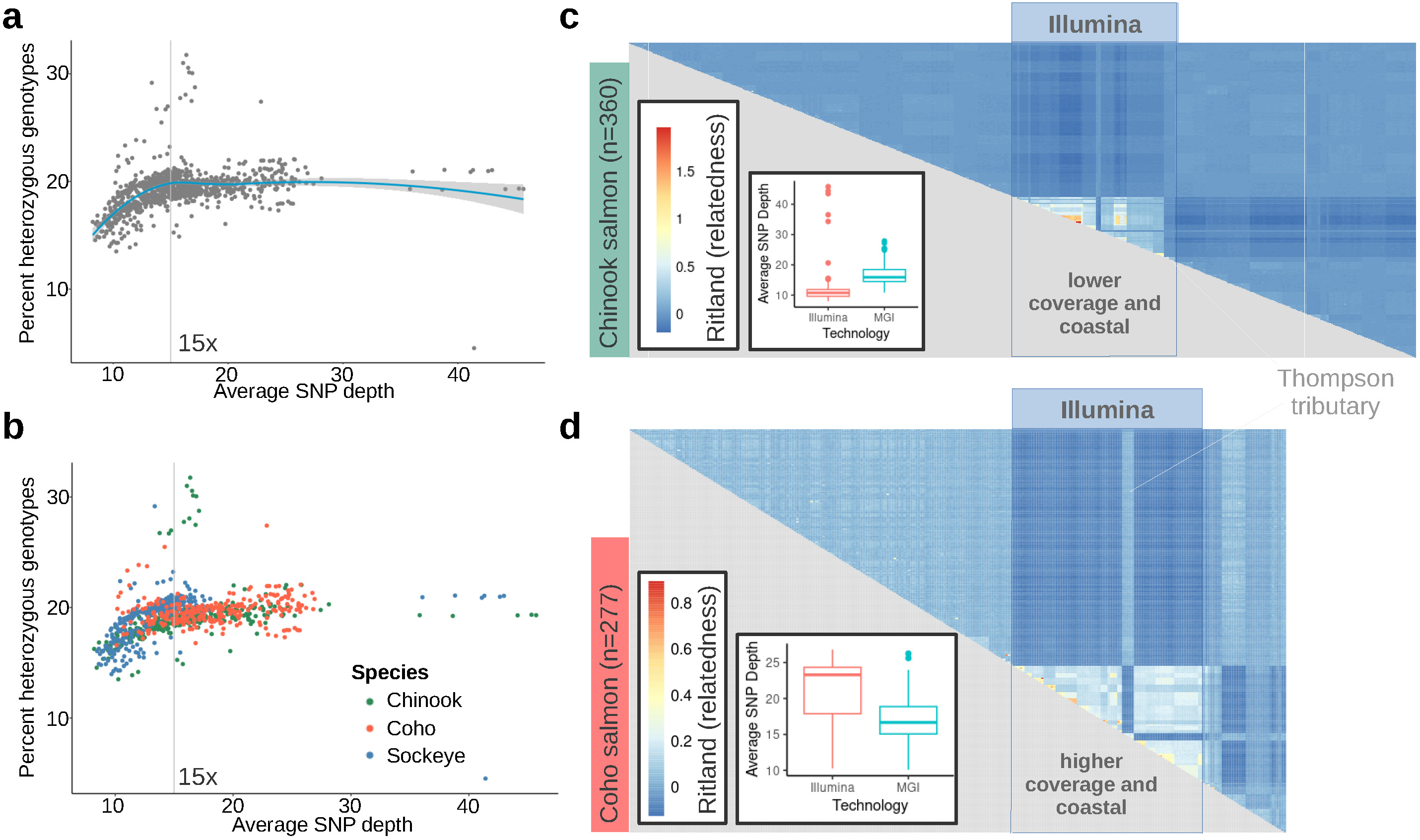
SNP coverage and relatedness metrics. a) A scatterplot of the relationship between the average SNP coverage and the percent of the genotypes that were heterozygous for each salmon (all species). The relationship between the two variables was modelled using the loess method. The vertical line shows 15x coverage, which was used as a threshold in other analyses. b) Same as a), but each salmon is identified by species. c) Heatmap of pairwise relatedness scores between each Chinook salmon. The salmon from a previous study are highlighted under the Illumina banner. A boxplot of SNP depth is displayed in an insert for the previous (Illumina) and current datasets (MGI). Samples from a Thompson River tributary are highlighted from the previous dataset. d) Same as c) for the coho salmon datasets.

Coastal samples (all sites downstream of the Thompson River or outside the Fraser River – see Figure 1) appeared to be highly related in Chinook and coho salmon (Figure 3c and 3d). These high relatedness scores were independent of the coverage or the technology used. In Chinook and coho salmon, coastal samples had much higher counts of private alleles (see below), which is known to increase relatedness metrics (Ritland 1996). Relatedness values were greatly reduced by sub-setting individuals by coastal or non-coastal (data not shown).

### Population structure and clustering analyses

We focused on large-scale resolution of genetic groups in this study, but there was still substantial genetic variation within each of the genetic groups (Figure S3). Each species had three supported Fraser River admixture groups (Figures 4 and S3). All groups were based on geography and were supported by PCA (Figure S4). These included a lower/coastal, mid, and upper Fraser River admixture group.

**Figure 4.**
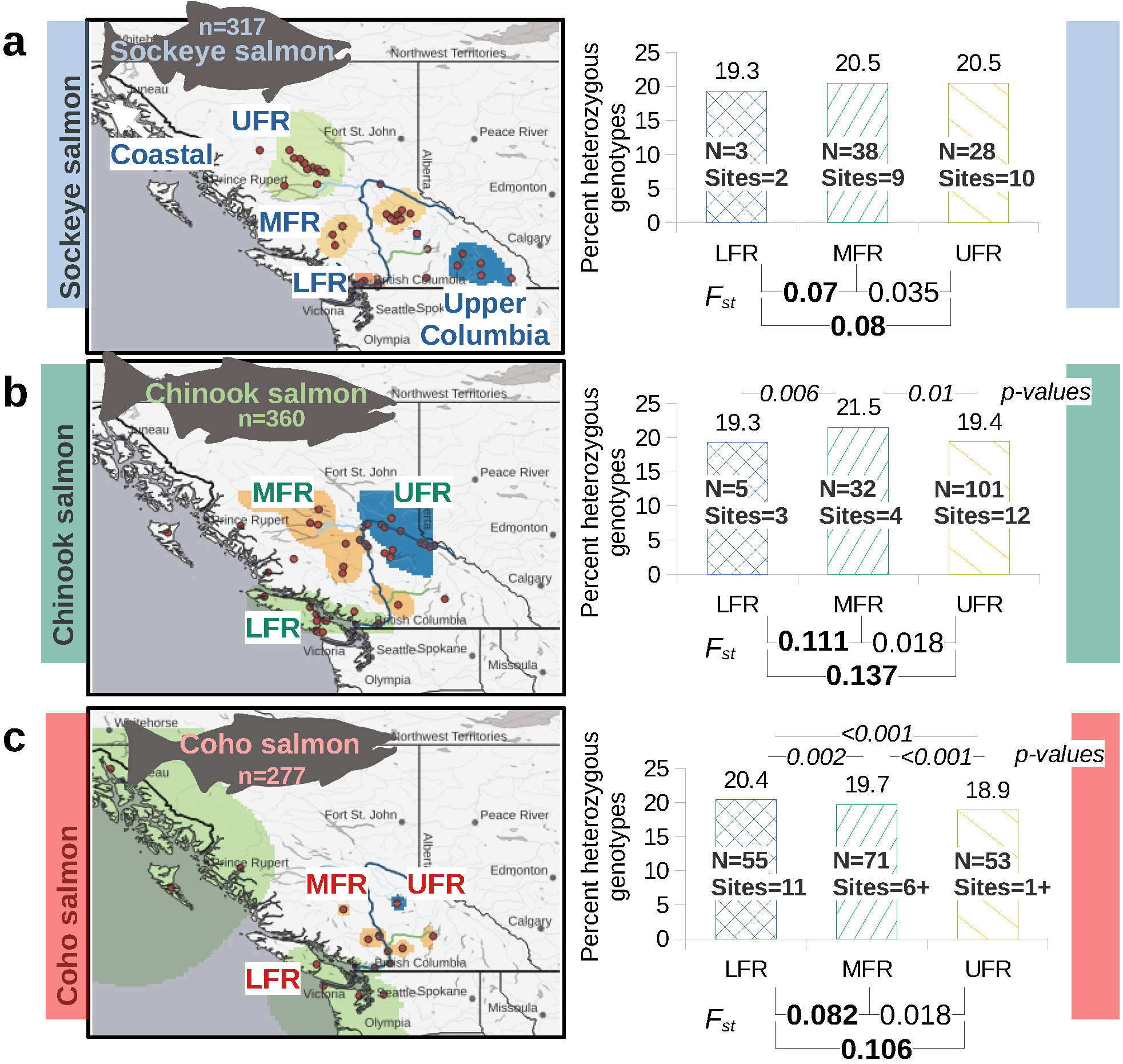
Comparison of Fraser River salmon admixture groups. a) (left) Map of sockeye salmon admixture groups plotted in QGIS by overlaying the admixture ancestry raster plots from Figure S3. Each group is labelled (the colour of the group was randomly assigned). The northern coastal admixture group is outside the frame of this map. (right) Average percent of heterozygous genotypes for each of the Fraser River admixture groups. The different Fraser River admixture group names are lower Fraser River – LFR, mid Fraser River – MFR, and upper Fraser River – UFR. The number of samples (≥15x SNP coverage) and sampling sites per admixture group is shown. Weir and Cockerham’s (Weir and Cockerham 1984) *F_st_*are shown for comparisons between admixture groups. *F_st_* comparisons between coastal (LFR) and interior clusters (MFR and UFR) are in bold text. b) Same as a) for Chinook salmon. The LFR group includes coastal sites outside the Fraser River. Significant differences in the percent of heterozygous genotypes among groups are shown with the *p-value* (two-tailed Welch’s t-test). c) Same as b) for coho salmon.

All of the lower Fraser River (LFR) admixture groups included sample sites in the Fraser Valley south of the Fraser Canyon (approximately where the river turns from a southern flow to a western flow before its outlet to the Pacific Ocean) (Figure 4). In Chinook and coho salmon, the LFR admixture groups also included nearby coastal sites. For coho salmon, this included locations from Alaska to California.

We sampled fewer coho salmon north of the Fraser Valley than for the other species due to accessibility issues during their spawning season, but those north of the valley clustered in a pattern similar to Chinook salmon (Figure 4). All samples, excluding those from McKinley Creek (a tributary of the Horsefly River), and some samples without known natal streams (sampled at the Big Bar Landslide), form one admixture group. In Chinook salmon, a similar admixture group is formed from the collection of sites west of the Horsefly River and from the Nechako River south. This geographic region is known as the Interior Plateau. In sockeye salmon, the mid Fraser River (MFR) group is composed of bodies of water from the Chilcotin and Quesnel River watersheds (Figure 4).

The upper Fraser River (UFR) coho and Chinook salmon admixture groups include most of the Quesnel watershed locations, with the exception of the lower Cariboo River near the confluence with the Fraser River (Figure 4). For coho salmon, this was from a single known sampling location and potentially other locations from salmon that were collected at the Big Bar Landslide. The Chinook salmon UFR group also included sampling sites above the Nechako River (in or near Robson Valley, a part of the Rocky Mountain Trench). The UFR sockeye salmon admixture group is comprised of Nechako River watershed sites.

The LFR admixture groups, in all species, have the greatest genetic differentiation (i.e., *F_st_*) from the other admixture groups (Figure 4). In addition, the UFR admixture groups have higher differentiation from the LFR admixture groups than the MFR groups (Figure 4). *F_st_* values were generally much lower between UFR and MFR admixture groups. While there were some significant differences in the percent of heterozygous genotypes per individual among groups, these do not show a consistent pattern (Figure 4). In Chinook salmon, the significant differences in heterozygous genotypes per individual between the MFR admixture group and the other groups were driven by high values in upper Chilcotin River samples (File S1).

### Environmental factors influencing genetic structure

Sampling locations from different admixture groups clustered by environmental factors (Figure 5, Figure S5). Sites with predominately LFR ancestry values were in regions with higher annual precipitation and higher mean annual temperatures (Figure S5). While there was some overlap between MFR and UFR admixture groups, there was still distinct environmental factors between them overall. This was the case for all three species.

**Figure 5.**
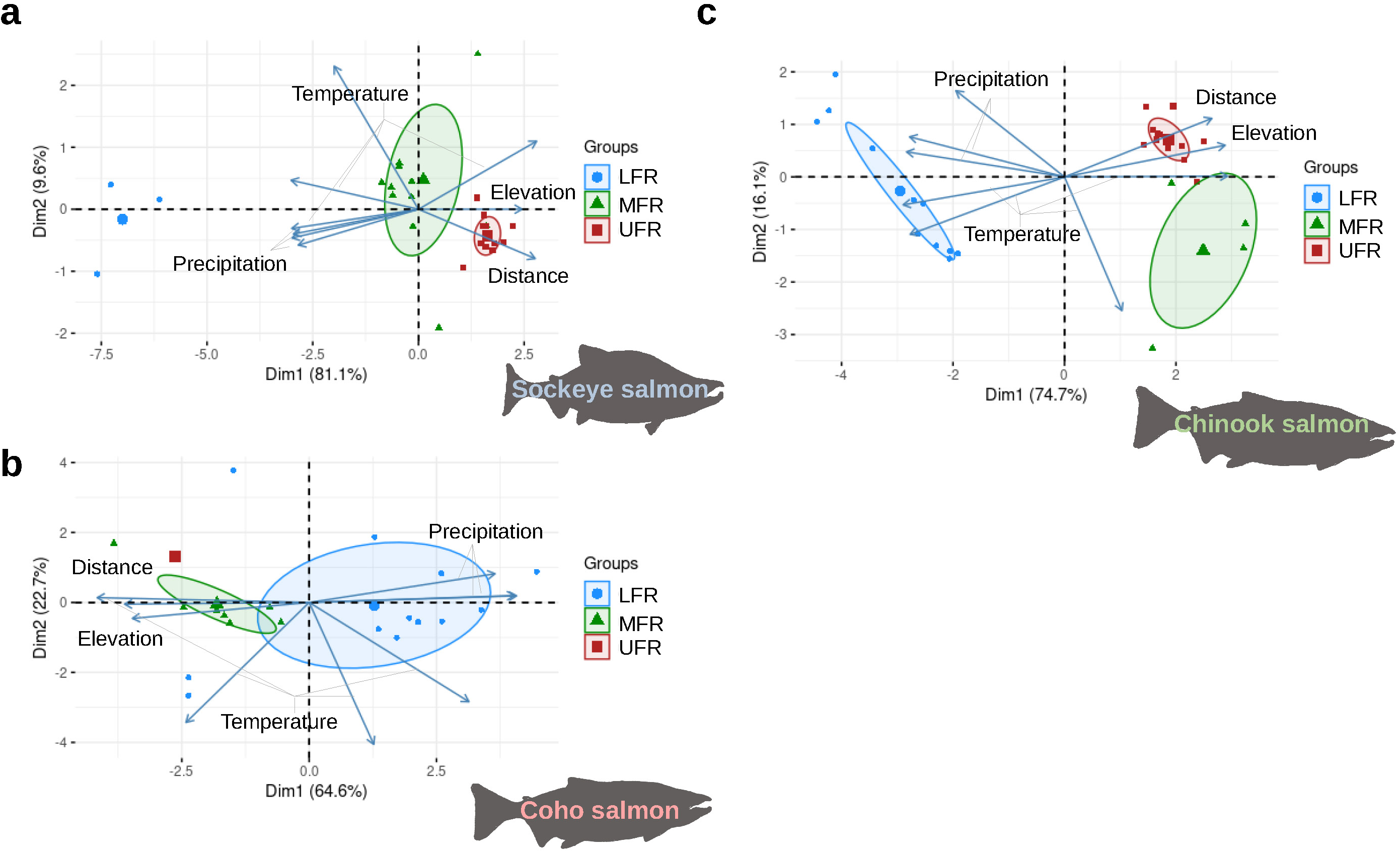
PCA of sampling location environmental factors. a) A biplot of a PCA of environmental factors from each sockeye salmon sampling site. These included environmental factors from WorldClim version 2.1, elevation, and distance to the ocean. Environmental factors were simplified as temperature (including: annual mean temperature, minimum temperature of the coldest month, maximum temperature of the warmest month, and temperature range), precipitation (including: driest month precipitation, wettest month precipitation, and annual precipitation), elevation, and distance to the ocean. Figure S5 does not simplify these variables. Locations were coloured by their assignment to admixture genetic groups, but no genetic data was used to produce these PCA. Only sites that had average ancestry values ≥ 0.7 from the LFR, MFR, and UFR genetic groups are shown for simplicity. See Figure S5 for all locations. Each small symbol represents a sampling site, and the larger symbol represents the centre of the ellipse if drawn (ellipse level 0.4). b) Same as a), except for coho salmon. The LFR group includes coastal sites outside the Fraser River. c) Same as b), except for Chinook salmon.

### Historical estimates of effective population size

Historical estimates of effective population size (N_e_) are strikingly different among species (Figure 6, Figure S6). Prior to 200,000 years before present, almost all Chinook and multiple coho salmon sampled experienced drops in estimates of effective population size (Figure 6, Figure S6). This coincides with a time when there was an increase in ocean surface temperatures (Figure 6d).

**Figure 6.**
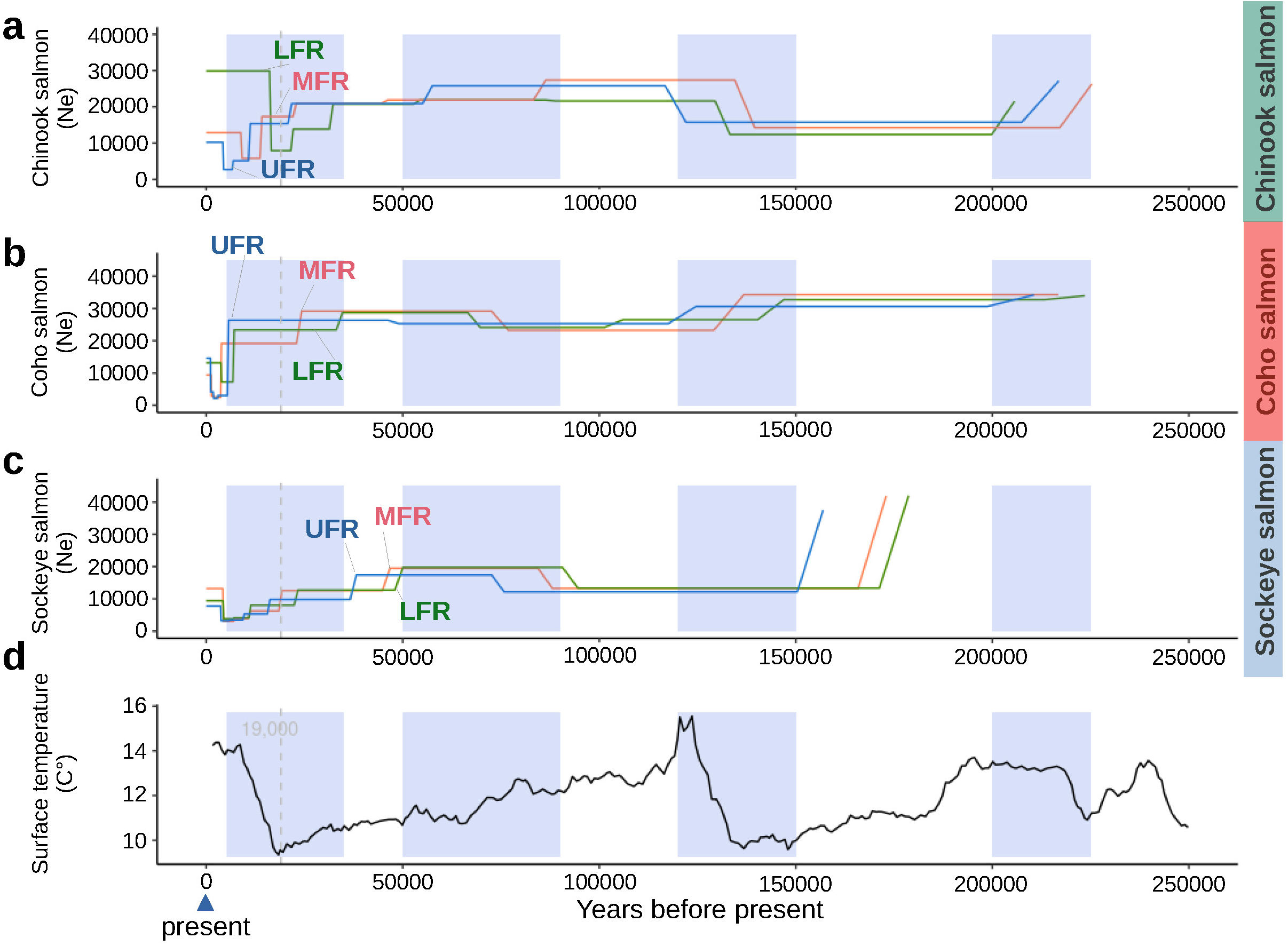
Estimated historical effective population size of Fraser River salmon. a) Effective population size of Chinook salmon estimated using SMC++ and whole genome data. Only one sampling site from each Fraser River admixture group was retained for clarity. Figure S6 shows all sites. The estimated glacial maximum of around 19,000 years ago (Clark *et al*. 2009) is shown with a dashed vertical line. Other regions are highlighted when there were major changes in effective population size for more than one species. b) Same as a) for coho salmon. c) Same as a) for sockeye salmon. d) Estimates of ocean surface temperature (Zachos *et al*. 2008; Hansen *et al*. 2013).

Between 120,000 and 150,000 years before present, coho and sockeye salmon both experienced major declines in estimated effective population size (Figures 6 and 7). In sockeye salmon, there are distinct times when these decreases occurred (Figure 7). For some sockeye salmon locations and for most coho salmon sites, the decrease in effective population size occurred around the penultimate glacial maximum, when ocean temperatures were at their lowest. Neither species recovered to previous estimates of effective population size for over 100,000 years (Figures 6 and S6). This is consistent with the decrease in polymorphic loci and increases of runs of homozygosity compared to Chinook salmon. Interestingly, Chinook salmon effective population size increased during this same time frame. A similar reversal among species was observed between 50,000 and 90,000 years before present, where there were increases in estimates of effective population size for coho and sockeye salmon, but decreases in Chinook salmon (Figure 6, Figure S6).

**Figure 7.**
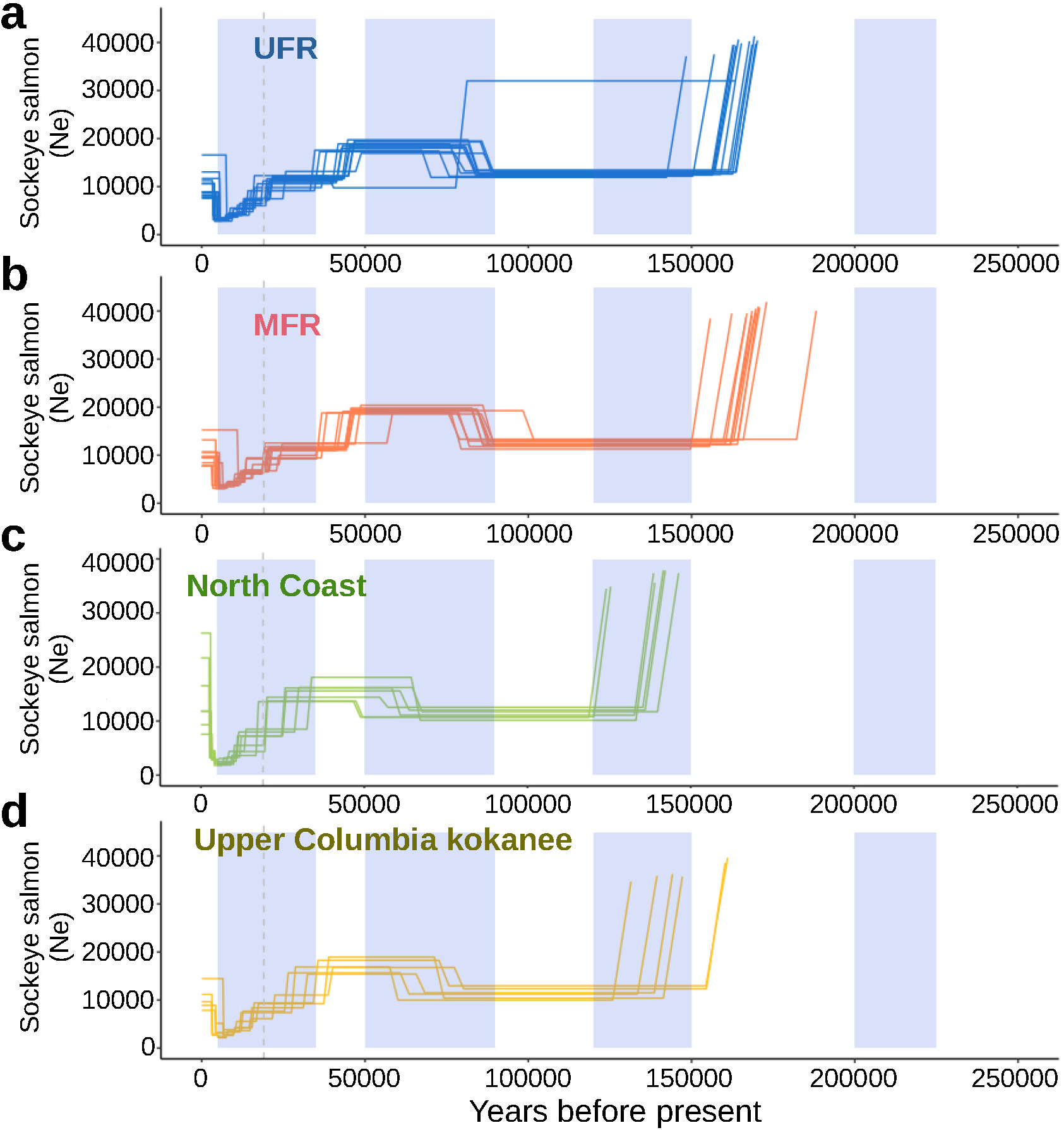
Estimated historical effective population sizes of some admixture groups of sockeye salmon. a) Effective population size of sockeye sampling locations from the UFR admixture group estimated using SMC++ and whole genome data. b) Same as a), but for sites from the MFR admixture group. c) Same as a), but for locations from the northern coastal admixture group. d) Same as a), but for locations from the upper Columbia River kokanee admixture group.

In all species, there were decreases in effective population size around the last glacial maximum, 19,000 years before present (Figures 6 and 7, Figure S6). The timing of these decreases in effective population size varied in time by species, admixture group, and sampling sites within admixture groups (Figures 6 and 7, Figure S6). Most of the demographic histories have a step pattern near or after the last glacial maximum. Some coastal Fraser River groups have earlier decreases in effective population sizes than the other Fraser River groups. The exception being LFR sockeye salmon sites, which have similar or more recent decreases in population size than MFR and UFR locations (Figure S6).

### Understanding the genetic diversity of Fraser River salmon

On average, total runs of homozygosity (ROH) were shorter in LFR locations than MFR and UFR sites of Chinook and coho salmon (Figure 8). Lower ROH is a proxy for greater genetic diversity, suggesting that the Chinook and coho salmon LFR admixture groups have higher diversity than MFR and UFR groups. Sampling sites with the longest average total ROH were the Yakoun River (Haida Gwaii) and Salmon River (tributary of the Thompson River) for Chinook and coho salmon, respectively (Figure S7). Locations with the shortest average total ROH (i.e., highest diversity) for Chinook and coho salmon, respectively, were the upper Chilcotin River (tributary of the Fraser River) and the Deschutes River (tributary of the Columbia River) (Figure S7). ROH levels were most variable in the Morkill and upper Chilcotin River locations for Chinook salmon. The individuals with the shorter ROH in the upper Chilcotin River drove the average to the lowest for Chinook salmon.

**Figure 8.**
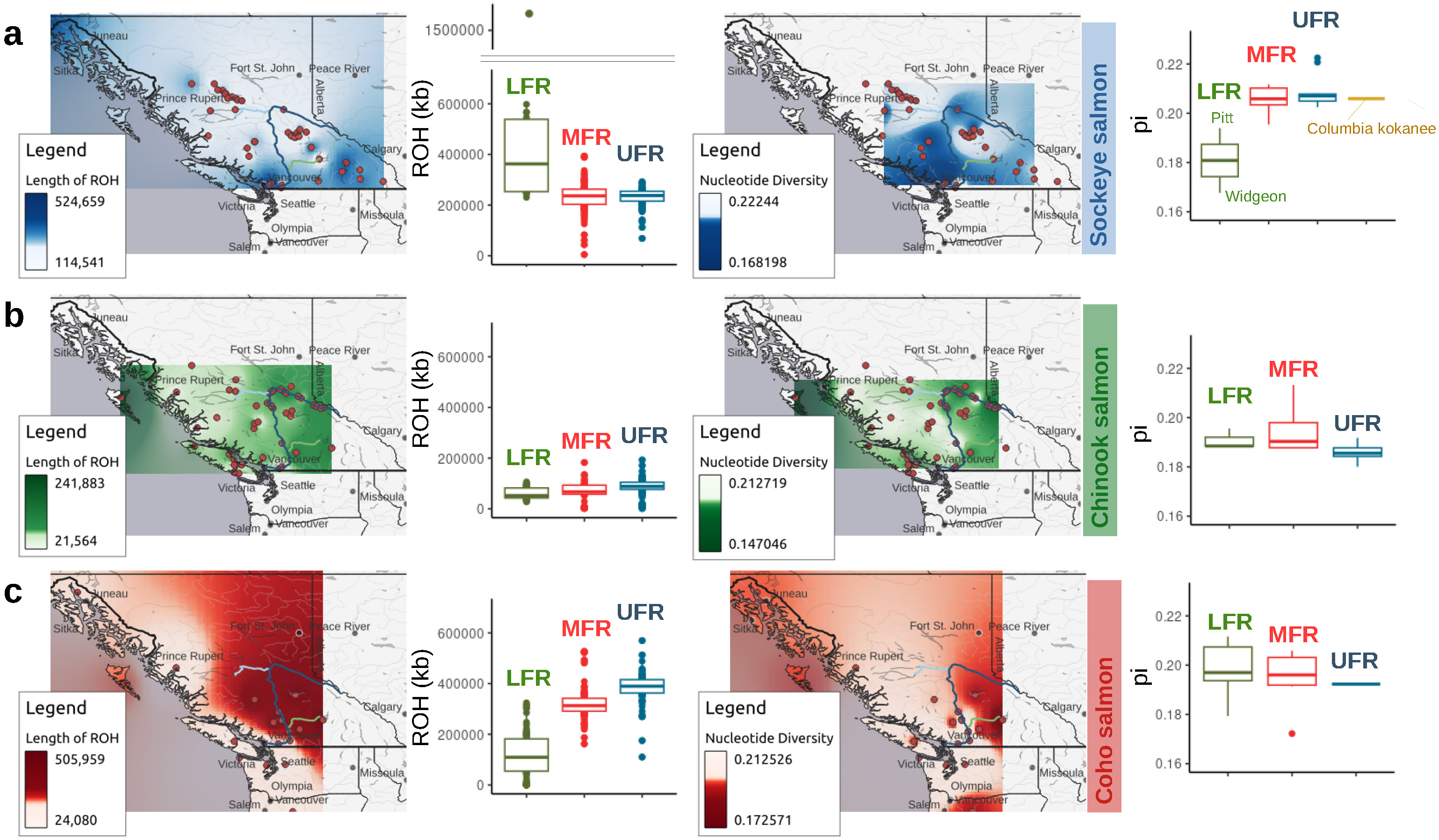
Genetic diversity of Fraser River salmon. a) (left) Map of sockeye salmon total length (kb) of runs of homozygosity (ROH) averaged for each sampling site. This analysis included all individuals. The colour scale is quantile-based. (middle-left) Barplot of total lengths of runs of homozygosity for samples assigned to the different admixture groups with ancestry values ≥ 0.7. (middle-right) Map of sockeye salmon nucleotide diversity (pi) of each sampling site excluding salmon with less than 15x SNP coverage. The colour scale is quantile-based. (right) Nucleotide diversity barplots of sampling sites with average admixture ancestry values ≥ 0.7 from individuals with at least 15x SNP coverage. b) Same as a) for Chinook salmon. The LFR group contains samples outside the Fraser River. c) Same as b) for coho salmon.

There was a reverse trend for ROH in sockeye salmon compared to Chinook and coho salmon (Figure 8). Most of the sockeye salmon sites with the longest ROH (i.e., the lowest diversity) were from the LFR (i.e., Cultus Lake, Pitt River, and Widgeon Slough). Cultus Lake and Widgeon Slough have small population sizes, often under 1000 salmon in recent generations (DFO 2020; Doutaz *et al*. 2023). The Harrison River, although not a part of this admixture group due to being below the 0.7 admixture threshold, had the lowest average ROH of all sockeye salmon collections (Figure S7). The Harrison River, in the early 1980s, had around 300,000 spawning adults (Doutaz *et al*. 2023). Widgeon Slough and possibly the Harrison River were also the only samples of the sea-type ecotype taken from the Fraser River (Beacham and Withler 2017; Doutaz *et al*. 2023), all the other Fraser River samples were likely lake-type. Also, some Harrison River salmon are sea and some are lake-type. Bodies of water in the Yukon, Alaska, Russia, and the upper Columbia River had slightly lower average ROH (i.e, more diverse) compared to the LFR, but higher than sites from the MFR and UFR (Figure 8 and Figure S7).

Nucleotide diversity (π), which is a population-level metric rather than an individual-level metric like ROH, had a more complex pattern. Nucleotide diversity ranged from 0.147 (Chinook salmon – Yakoun River) to 0.22 (sockeye salmon – Pinchi Creek). All sockeye salmon sampling locations had nucleotide diversity metrics above 0.19 except for Widgeon Slough, which was 0.17. Cultus Lake might have had lower nucleotide diversity than Widgeon Slough based on its longer ROH, but this location was excluded from this analysis since it had lower SNP coverage than 15x. Similarly, most northern coastal and upper Columbia River samples were excluded from the nucleotide diversity analysis because they had SNP coverage below the threshold. Most Fraser River salmon locations had comparable nucleotide diversity to upper Columbia River Kokanee that were available for comparison. Similar to the ROH results, the LFR salmon had lower diversity than the MFR and UFR sockeye salmon.

Most Chinook salmon locations had π values between 0.18 and 0.20. However, the Yakoun River (Haida Gwaii) had the lowest π of all sites for Chinook salmon (π = 0.147). Many of the sampling locations with the highest nucleotide diversity were from the MFR (Figure 8). The Fraser River Chinook salmon sampling sites had average nucleotide diversity values comparable or lower than other coastal locations along the BC coast (Figure 8).

Coho salmon ranged in nucleotide diversity from 0.17 to 0.21, with the lowest collection site being the Salmon River in the South Thompson River drainage and the highest being the Kawkawa Creek below the Fraser Canyon (Figure 8). Fraser River coho salmon had comparable or higher levels of nucleotide diversity as those from California to Alaska. The most extreme northern (Kwethluk River) and southern sites (Klamath River) had the second and third lowest nucleotide diversity metrics respectively.

The number of polymorphic loci was highest in Chinook salmon sampling sites, followed by coho salmon and then sockeye salmon, which is consistent with the total number of SNPs identified in each species (Figure S8, Figure 2). While the number of polymorphic loci was dependent on the number of samples per location, it remained higher in Chinook salmon at comparable sample counts per location (Figure S8). Even though Chinook salmon had the greatest number of polymorphic loci per location, nucleotide diversity was often lower than for coho and sockeye salmon locations (Figure S8). This is consistent with higher individual percent heterozygous genotypes in coho and sockeye salmon (Figure 3).

### Admixture group private alleles

There were more private alleles identified in the LFR admixture groups of Chinook and coho salmon than the other admixture groups, with 5-13% of all the identified SNPs being private to the LFR group (Figure 9a). Sockeye salmon had a more even distribution of private alleles among admixture groups than Chinook or coho salmon (Figure 9a). However, the sample size of the LFR sockeye salmon group was the lowest of all comparisons, and this may have impacted the number of private alleles identified. Also, sockeye salmon from the Harrison River (in the coastal region) had admixture ancestry value below 0.7 and were excluded from this analysis. This site had greater genetic diversity metrics than most other sockeye salmon locations (see previous section), and so the removal of this river would be expected to reduce the observed private alleles from this region.

**Figure 9.**
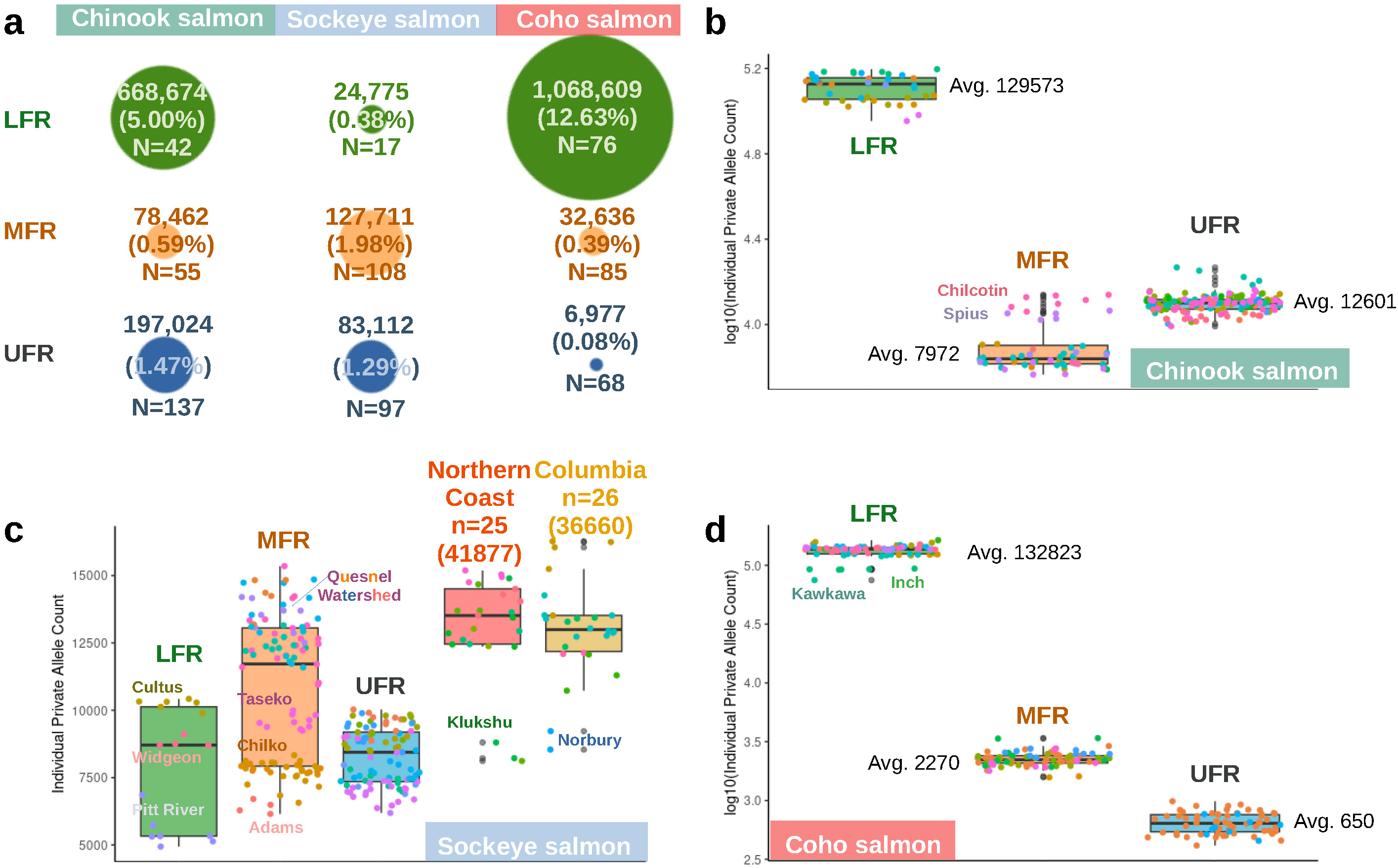
Private allele analyses of Fraser River genetic groups. a) The count of private alleles of Fraser River admixture groups (only individuals with ≥ 0.7 ancestry values were used). Only private alleles identified in at least three individuals from a group were counted. Below the private allele count is the percent that this represents of all the SNPs identified for each species. The circles are scaled by this value. The number of samples per genetic group after these criteria were met are also included. b) Boxplots, with jittered points added, of individual private allele counts of the different Chinook salmon admixture groups. The individual private allele counts are the number of private alleles each individual has from the private alleles common to the admixture group. This is a way to compare how much individuals contribute to the identified private alleles of the group, and to be able to compare these values to understand if sample size might influence group counts. The LFR group has individuals from outside the Fraser River. c) Same as b) for sockeye salmon, except there were five total admixture groups of sockeye salmon and the LFR is strictly Fraser River samples. d) Same as b) for coho salmon.

Analyses of individual private alleles from Chinook and coho salmon provided more evidence that LFR individuals have more unique alleles than individuals from MFR and UFR groups (Figure 9b-d). Each coastal individual had roughly 130,000 private alleles that were not found in the other admixture groups. Other admixture groups, on average, have much fewer individual private alleles compared to LFR salmon (∼ 0.04% -10%). The upper Chilcotin River and Spius River contributed more private alleles to the Chinook salmon MFR admixture group than other rivers in that group (Figure 9b). Likewise, the Kawkawa River contributed fewer private alleles to the coho salmon LFR group than the other rivers (Figure 9d).

Sockeye salmon do not have a comparable admixture group with the large store of private alleles relative to that observed in the LFR coho and Chinook salmon. The northern coast and upper Columbia River kokanee admixture groups have the highest average individual private allele counts (Figure 9c). Most sampling locations in the Fraser River have lower individual private alleles compared to those outside the Fraser River. The main exceptions are samples from the Quesnel River watershed, which have similar values as those outside of the Fraser River (Figure 9c). This is consistent with a higher nucleotide diversity in the Quesnel River watershed than the Chilcotin River watershed (see previous section).

### Potentially adaptive variants

Twenty candidate adaptive loci were identified in pairwise comparisons among admixture groups based on expanded haplotype homozygosity analyses (Figure S9, Table 2, File S3). No significant loci were identified from within admixture group analyses. Most candidate loci were identified among Chinook salmon admixture groups (14 out of 20). Twelve of these loci overlapped with the boundary of a gene (Table 2). Two different homologs of the *trace amine-associated receptor 13C* (TAAR13C) gene overlapped with candidate loci identified in comparisons of admixture groups of sockeye salmon (Figure S9, Table 2). One of these genes was identified from the comparison with the LFR group and the MFR group. The other was identified from the comparison of the LFR group and the UFR group.

**Table 2.**
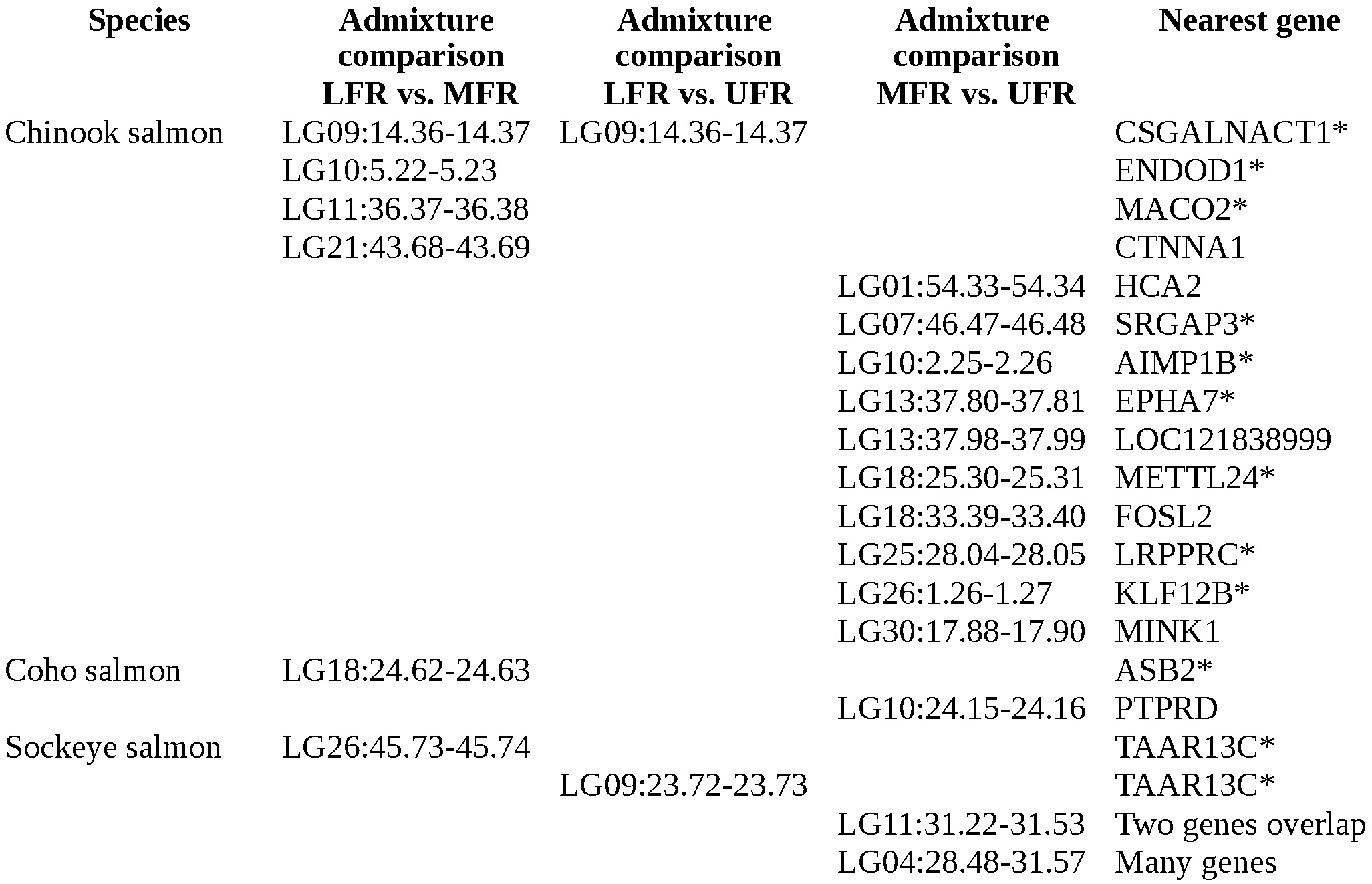
Significant extended haplotype homozygosity among admixture groups.

To identify if there were potential adaptive variants shared among species, we searched for overlapping regions with high -log10 p-values (we were searching for signals of convergent evolution with this analysis). The overlap of 10 kb regions with the highest -log10 p-value was 3.6 among species. The chance of having an overlap with this high of a value was expected by random at least twice by chance given the distribution of p-values. These overlaps were not considered significant and they are not shown or discussed below.

## Discussion

The Fraser River drains a very large and diverse section of British Columbia from the Rocky Mountains to its outlet in the city of Vancouver (Reynoldson *et al*. 2005). This river has at times been the largest producer of salmon in North America (Milne 1964; Northcote and Atagi 1997), and remains important commercially (Henderson and Healey 1993) and culturally for its salmon fisheries. The massive declines in salmon, predominately sockeye and pink salmon from a 1913-1914 landslide caused by the construction of a railway, have never fully recovered (Pacific Salmon Commission – psc.org). There are many pressures potentially for why recovery has not occurred and why there has been continued declines in Fraser River salmon (Arbeider *et al*. 2020; DFO 2021; Doutaz *et al*. 2023). Understanding these pressures and their influence on salmon will be valuable for governance and conservation. Our goal is to provide the genetic tools that will facilitate this understanding and help with recovery.

While there have been many studies to examine population structure in the Fraser River (Small *et al*. 1998; Teel *et al*. 2000; Withler *et al*. 2000; Beacham *et al*. 2001, 2003, 2006b, 2006a, 2017; Nelson *et al*. 2001; Beacham and Withler 2017; Xuereb *et al*. 2022), this is the first to use whole genome resequencing data for a large distribution of multiple salmon species. The improved resolution of resequenced genomes allowed us to compare the genetic diversity among salmon species, calculate nucleotide diversity at the genome level, and evaluate historical influences on population structure.

These types of analyses would suffer using fewer genetic markers. As an example, genetic and nucleotide diversity estimates are dependent on the nucleotide variants used to estimate them (e.g., polymorphic loci in one region might not be polymorphic in another region) and whole genome analyses allow unbiased estimates among locations. These genome sequences also us to capture a better snapshot in time that can be used in comparisons for future studies.

### Species level differences of polymorphic loci

Genetic diversity can be understood at both a species and population level. We first discuss the diversity at a species level and will return to the population level later. From previous research, we expected different levels of polymorphic loci among Pacific salmon, with a generally lower genetic diversity of Pacific salmon compared to other species such as rainbow trout (*O. mykiss*) (Allendorf and Utter 1979). Several studies observed that sockeye salmon, and sometimes coho salmon, have the lowest levels of polymorphic loci or average heterozygosity (Utter *et al*. 1973; Allendorf and Utter 1979; Wood *et al*. 1994).

Our findings indicate that Chinook salmon have ∼1.5-2x the number of polymorphic loci and a corresponding drop in the length of ROH (∼27-29% of the ROH length) when compared to coho or sockeye salmon. We observed the same trend based on the total number of heterozygous genotypes among the species when only looking at individuals with higher than 15x coverage. While sequencing coverage likely impacts the number of polymorphic loci identified, the scale of the differences we observed, the greater geographic sampling distance covered by the species with fewer polymorphic loci, a similar trend in ROH, and a similar trend with heterozygous genotypes of individuals with at least 15x coverage supports that these findings are robust. If we assume that salmon species had similar levels of genetic diversity during speciation and that mutation rates are similar among species, coho and sockeye salmon likely had a greater reduction in standing genetic variation than Chinook salmon.

From modelling of effective population size through time, there is evidence that sockeye and coho salmon experienced a large drop in effective population size around the penultimate glacial maximum that Chinook salmon did not experience (Chinook salmon did experience an earlier drop in effective population size, but the size rebounded after ∼50,000 years). We must consider that we do not have precise estimates of mutation rates for all of these species. Even without precise dates or effective population size estimates, however, we note that the drop in effective population size did not rebound to prior levels in sockeye and coho salmon as it did in most Chinook salmon sites. This could have resulted in a loss of polymorphic loci in both sockeye and coho salmon not observed in Chinook salmon.

While the SNP count, effective population size, and runs of homozygosity data are consistent with this hypothesis, nucleotide diversity and private allele analyses appear to be inconsistent at first glance. For example, why would sockeye and coho salmon have higher nucleotide diversity if Chinook salmon have more polymorphic loci overall (sockeye salmon average π: 0.205, coho salmon: 0.197, Chinook salmon: 0.189)? Also, why would the average private allele count be higher in the LFR coho salmon admixture group than the Chinook salmon LFR group if the Chinook salmon have more polymorphic loci?

First, we note that some coastal samples of sockeye salmon were removed due to low coverage in these analyses, and these samples would have likely lowered the average nucleotide diversity score. Technical artifacts like this and others (e.g., how nucleotide diversity was measured, sequence coverage, or differences among genome assemblies) might help to explain a part of these discrepancies, but they appear to be inadequate for all of the data. For example, at comparable SNP coverages, sockeye salmon had higher proportions of heterozygous genotypes than coho or Chinook salmon even though coho and Chinook salmon have more polymorphic loci overall.

The timing of re-colonization and the distribution of groups with unique genetic characteristics could help to account for some of the discrepancies among genetic diversity metrics. In Chinook and coho salmon, for example, the coastal genetic groups have the largest stores of private alleles and the highest metrics of genetic diversity (comparable between species, even if slightly higher in coho salmon). If there was a difference in the timing or the size of the groups that re-colonized the Fraser River from these coastal genetic groups, we might expect the fraction of private alleles from a group to reflect these differences.

Estimates of effective population size dip in Chinook salmon locations soon after the last glacial maximum when compared to coho salmon. If we assume these dips are founding events and the times are roughly accurate, we infer that Chinook salmon re-colonized the Fraser River earlier. The founding events also appeared to have reduced the effective population size of Chinook salmon sites less than they did for coho salmon, perhaps reflective of larger founding populations in Chinook salmon. With this in mind, even with fewer polymorphic loci overall, the LFR coho salmon might appear to have higher levels of private alleles per individual than Chinook salmon when comparing the LFR group to the MFR and UFR groups because those groups have lower genetic diversity as a result of more recent and smaller founding events. The MFR and UFR coho salmon groups could have retained comparable nucleotide diversity metrics to Chinook salmon groups due to gene flow, or because not as much time has passed for genetic drift to impact this metric as it has for Chinook salmon. Also, heterozygosity excess (e.g., (Wang *et al*. 2016)), where allele frequency varies by chance more often between males and females due to a low breeding population, is another viable explanation for how MFR or UFR coho salmon might have comparable nucleotide diversity to Chinook salmon.

Nucleotide diversity might also be higher in a species with lower overall counts of polymorphic loci and genetic diversity if there was recent admixture between diverse populations. In sockeye salmon, there were at least two distinct demographic patterns identified based on when effective population size declined around the penultimate glacial maximum (perhaps indicative of groups from different glacial refugia as previously suggested (Wood *et al*. 1994; Beacham *et al*. 2006b)). In the Fraser River, both demographic patterns were observed, with the MFR and UFR admixture groups mostly having the earlier decline in effective population size (earlier than 150,000 years before present), while the LFR locations had mostly the later (after 150,000 years before). Gene flow between these groups could explain the higher nucleotide diversity in the MFR and UFR even though sockeye salmon as a whole have fewer polymorphic loci.

Overall, the data point to a model where Chinook salmon have more polymorphic loci, but less diversity at these loci than coho or sockeye salmon. Demographic modelling points to one explanation for how this might have occurred, but other explanations are possible. For example, heterozygosity excess is expected in populations with smaller effective population size (Robertson 1965; Wang *et al*. 2016).

### Influences on the population structure of Fraser River salmon

Turning from the species level to the population level, researchers have known the basic population structure of Fraser River salmon since the mid 1990’s to the mid 2000’s (Wood *et al*. 1994; Small *et al*. 1998; Teel *et al*. 2000; Withler *et al*. 2000; Nelson *et al*. 2001; Beacham *et al*. 2003, 2006a, 2006b, 2017; Beacham and Withler 2017; Xuereb *et al*. 2022). This knowledge was based on relatively few genetic markers but often many samples from a wide distribution. In the current work, we sampled a moderate number of locations and individuals, but resequenced entire genomes. The basic population structure identified in the Fraser River from resequencing genomes was similar to findings from earlier studies using microsatellite and allozyme genetic markers (e.g., lower, mid, and upper groups), except the Thompson River groups were missing since we did not sample the Thompson River drainage enough to reconstruct them. We interpret the consistency among studies to mean that these genetic clusters are stable through time (as sampling took place at various times, even within studies (Beacham *et al*. 2003, 2006a)), the clusters are robust since they can be identified with only a few genetic markers, and the entire genome is impacted by these groupings.

Understanding why we observe these genetic groups is important (e.g., (Nadeau *et al*. 2016; Rougemont *et al*. 2023)). Are they a technical artifact from trying to cluster genomes influenced by isolation-by-distance? Are they an artifact from the transfers between regions of the Fraser River (Withler 1982)? Were they formed from different re-colonization events (or a combinations of events)? Are they a consequence of environmental hurdles to gene-flow? Were they formed from adaptations to specific environmental conditions?

If the groups are technical artifacts from isolation-by-distance, they may not be useful from a conservation or management perspective because the groups were shaped by stochastic processes rather than adaptations to different environments. If instead, groups were shaped by different re-colonization events, they might reflect those colonization histories rather than the environments they occupy. If the groups are a result of limited gene flow, we might expect the salmon to have specific adaptations to environmental factors that influence gene flow (e.g., the ability to pass regions of the Fraser River such as the Fraser Canyon), but not necessarily to other environmental conditions. These are only some examples of why these groups might exist. The genetic groups could also be influenced by a combination of these processes.

Previous researchers have suggested that isolation-by-distance is an important factor influencing salmon genetics (Withler *et al*. 2000; Rougemont *et al*. 2020, 2023). Others have suggested that different re-colonization events (e.g., from different glacial refugia) influence population structure in Fraser River salmon (Wood *et al*. 1994; Small *et al*. 1998; Teel *et al*. 2000; Withler *et al*. 2000). Still others have suggested that barriers to gene-flow like the Fraser Canyon (specifically Hell’s Canyon) are important for population structure (Wehrhahn and Powell 1987). Finally, researchers have also suggested genetic adaptation to environmental conditions are important (Small *et al*. 1998). Analyzing multiple species can help with distinguishing which hypotheses are more supported (Nadeau *et al*. 2016).

While we observed lower *F_st_* between neighbouring admixture groups in all species, which would be consistent with isolation-by-distance, we also observed that private alleles were more common in the UFR group of Chinook salmon than in the MFR group. If populations were delineated strictly by isolation-by-distance, we might expect UFR salmon to have the lowest count of private alleles. Also, in all three species, sampling sites of an admixture group clustered together based on environmental factors in a PCA. If admixture groups were the result of neutral genetic variation or different colonization events alone, we would expect random sorting of groups or for it to be based on distance between locations. This suggests that the environment is a major factor influencing genetic variation in Fraser River salmon, but still does not exclude isolation-by-distance or separate colonization events.

Indeed, from demographic modelling, separate colonization events appear to have been an important source of sockeye salmon in the Fraser River and possibly for Chinook salmon. For sockeye salmon, there are at least two unique demographic histories, with one common to LFR sites and the other to MFR and UFR locations. These groups have unique demographic histories starting as early as the penultimate glacial maximum. In Chinook salmon, LFR sampling locations have an earlier decrease in effective population size that may be indicative of an earlier colonization history. Overall, the variability in demographic histories of the species suggest a complex re-colonization history.

In all three species, there was a delineation between admixture and PCA groups between the LFR group and the MFR group. The LFR groups had higher *F_st_* values among groups, and 5-13% of the SNPs we identified in Chinook and coho salmon were private alleles of the LFR group. If we also consider the velocity barriers between these groups (Wright 2022), we have several pieces of evidence to suggest that there is a substantial gene flow and migration barrier due to the Fraser Canyon. In support of this hypothesis, we observed that the Chinook and coho salmon from the LFR groups had higher nucleotide diversity and that longer runs of homozygosity in coho salmon appeared to be demarcated near Hell’s Canyon in the Fraser Canyon. If the Fraser Canyon is a migration barrier, we would expect to observe these genetic delineations in all species and generally this is what was observed.

In a study of Fraser River sockeye salmon, researchers observed possible adaptations to environmental differences among populations (Eliason *et al*. 2011). They observed coastal populations had significantly different cardiac morphology and performance from other groups, and that cardiac morphology was correlated with migration difficulty. River temperature was also correlated with cardiac performance. This study highlights the importance of the environment on population structure, migration difficulty, and relates the genetic structure among admixture groups to phenotypic variation. We note that in a comparison of extended haplotype homozygosity between the LFR coho salmon with the MFR group, a potential adaptive locus was identified that overlapped with the *Ankyrin Repeat And SOCS Box Containing 2* (ASB2) gene. This gene is thought to be involved in heart development (Yamak *et al*. 2020; Min *et al*. 2021).

### Influence of adaptation on population structure of Fraser River salmon

When comparing genetic groups, we identified twenty candidate adaptive loci. No significant loci overlapped among species, which would have been evidence for convergent evolution. Rather than discuss all twenty regions, we will focus on what these loci reveal in general, and as an example discuss the olfactory receptor gene TAAR13C. Potentially adaptive loci overlapped with two TAAR13C genes and were identified in separate comparisons of the LFR sockeye salmon with MFR and UFR groups.

In general, adaptive loci among the genetic groups reveal that there are different selective pressures along the Fraser River drainage. While the environmental PCA gave us insight into how genetic groups were organized based on environmental components, adaptive loci can reveal how these and other elements shape the genomes of salmon through generations. In the type of analysis we used, we did not need to know what these elements were. This means we can discover adaptation caused by unknown and unexamined factors.

We may be able to formulate a hypothesis regarding a mode of adaptation in the case of the TAAR13C genes. The olfactory receptor gene, TAAR13C, is directly involved in detecting putrescine, cadaverine, and other diamines (typically associated with decomposing flesh) (Hussain *et al*. 2013; Tessarolo *et al*. 2014; Liberles 2015; Gainetdinov *et al*. 2018). The TAAR13C gene has also previously been found to be associated with sea age at maturity in Atlantic salmon (*Salmo salar*) (Sinclair-Waters *et al*. 2022), and to possibly be under selection in another study of Atlantic salmon (not peer reviewed at the time of writing) (Miettinen *et al*. 2023). Since the TAAR13C gene appears to be involved in the timing of maturation, one hypothesis is that diamines could act as a signal for maturation for the different genetic groups of sockeye salmon in the Fraser River. One source of diamines to consider for this hypothesis comes from eggs. A study of Arctic charr (*Salvelinus alpinus*) eggs and alevin revealed that during these developmental stages different amounts of putrescine and cadaverine were produced during the spawning season (Srivastava *et al*. 1992). If diamines influenced maturation of MFR and UFR groups of sockeye salmon, this information would be valuable in conservation and management. Understanding the modes of adaptation of other potential adaptive loci could likewise be useful for these purposes.

## Conclusion

From analyzing hundreds of resequenced genomes of Chinook, sockeye, and coho salmon, mostly from the Fraser River, we identified genetic groups that had previously been identified with only a few microsatellites. With data from resequenced genomes, we were able to examine how these groups might have formed. They appear to have been influenced by many factors, including isolation-by-distance, migration barriers, separate glacial refugia in sockeye salmon, and by the diverse environmental factors of the Fraser River drainage. We identified twenty potentially adaptive loci among the different groups of salmon, which is indicative of unique patterns of selection in the various regions of the river. Two of these loci overlapped with homologs of the TAAR13C gene. This gene could be an important target for future studies, to investigate if there is a link between it and timing of maturation in Fraser River sockeye salmon. Finally, by examining three species, we were able to identify commonalities and differences. Patterns of historical effective population size were dramatically different among the three species and could explain the variable genetic diversity currently observed among them. In terms of information useful from a conservation and management perspective, we have generated resequenced genomes that can be reused in future studies, determined metrics for comparison, and identified loci that could impact the timing of maturation of salmon in the Fraser River.

## Data availability

Resequenced genomes from this study are available on the NCBI (coho salmon: PRJNA986075, sockeye salmon: PRJNA930425, and Chinook salmon: PRJNA694998 and PRJNA1090956). Truth SNP datasets used for SNP calling are available in File S2. SNP datasets are available in FigShare.

Scripts are available at github.com/KrisChristensen (repositories: PrivateAllelePerInd, PrivateAllele, VCF2Relate, VCFstats, and MapVCF2NewGenome).

## Acknowledgements

We would like to thank the many people involved with sample collection at Fisheries and Oceans Canada (dfo-mpo.gc.ca). For computer resources, we would like to acknowledge the Digital Research Alliance of Canada (alliancecan.ca) and its regional partner the BC DRI Group at the University of Victoria. For their sequencing services, we would like to thank the Michael Smith Genome Science Centre and the technicians there.

## Supporting Information

**Figure S1. Expanded maps and sampling locations.** Slide 1) Sockeye salmon maps, Slide 2) Chinook salmon map, and Slide 3) Coho salmon maps.

**Figure S2. Relationship between average SNP coverage and the total length of runs of homozygosity.** a) The average SNP coverage per sockeye salmon sample relative to the total length of runs of homozygosity per sample. The line was determined using the loess function (local regression). b) Same as a) for Chinook salmon. c) Same as a) for coho salmon.

**Figure S3. Admixture analyses.** Slide 1) Average admixture ancestry values of the different admixture groups were plotted for sampling locations in QGIS using the inverse distance weighted interpolation method. The darker the sampling location, the greater the ancestry assignment to the specified admixture group. Slide 2) Plot of admixture ancestry values for clusters above and below the best supported k=5 for sockeye salmon. Slide 3) Plot of admixture ancestry values for clusters above and below the best supported k=3 for Chinook salmon. The arrows indicate that the name of a location was omitted because of space limitations. Slide 4) Plot of admixture ancestry values for clusters above and below the best supported k=3 for coho salmon. The arrows indicate that the name of a location was omitted because of space limitations.

**Figure S4. PCA of each species with different filtering criteria.** a) PCA of sockeye salmon. The first column is a PCA of all samples, the second column is a PCA with only one individual from pairs with high relatedness scores (≥ 0.15), and the third column is a PCA of samples with high SNP coverage (>15x coverage). The different admixture groups are noted in the PCA. Admixture numbers were arbitrary, so names were given based on geography for the first column PCA. Individuals with admixture ancestry values < 0.7, were given a different colour from the the main groups. b) Same as a) for Chinook salmon. c) Same as a) for coho salmon.

**Figure S5. PCA of sampling locations based on environmental factors.** Slide 1) Biplot of sockeye salmon sampling sites based on environmental variables from WorldClim version 2.1 and elevation. Sampling sites with an average ancestry assignment of less than 0.7 were assigned to intermediate genetic groups with the highest fraction named first (e.g., if a site had average ancestry fractions of 0.6 MFR and 0.4 LFR, it was assigned Mid-Coast). The Okanagan was an exception as it had high ancestry values for multiple genetic groups. Each sampling location is represented by a small point (unless there was only one site, in which case it has a larger symbol), larger points show the middle of ellipses, and ellipses were drawn at the 0.4 level. The average annual precipitation and distance to the ocean are shown on the right. Groups were combined if they shared a genetic group as the largest contributor to their ancestry values. Slide 2) Same as Slide 1, but for coho salmon, Slide 3) Same as Slide 1 but for Chinook salmon.

**Figure S6. Estimates of historical effective population size for different salmon species and groups.** Slide 1) Effective population size for all species and sampling sites. Some Chinook sampling locations are off the graph. These sites may have recent admixture, which can throw off these estimates. These sites were removed in all other figures. Slides 2-3) Fraser River admixture group comparisons of each species. Slides 4-5) Effective population size estimations of different groups of Chinook salmon. Individuals were grouped by admixture and region (Coastal: Northern BC Coast, Vancouver Island, Lower Fraser River; Mid Fraser: Mid-Upper Fraser River, Mid-Coast Fraser River, Mid Fraser River; Upper Fraser: Upper Fraser River, Upper Mid Fraser River). Slides 6-7) Effective population size estimations of different groups of coho salmon (Coastal: Mid Alaska, Northern BC/Southern Alaska, Southern BC, Coastal Fraser River, California-Oregon; Mid Fraser: Mid Fraser-Coastal Fraser, Mid Fraser; Upper Fraser River). Slides 8-9) Effective population size estimations of different groups of sockeye salmon (Coastal: Northern Coast, Coast-Mid Fraser, Coast-Coast Fraser, Coast Fraser; Mid Fraser: Mid-Coast Fraser, Mid-Upper Fraser, Mid Fraser; Upper Fraser: Upper Fraser – Columbia, Upper Fraser River; Columbia: Upper Columbia River Kokanee, Upper Columbia – Upper Fraser; Okanagan Lake).

**Figure S7. Total runs of homozygosity for each sample location.** Slide 1) Boxplot of total runs of homozygosity (kb) for all individuals from each sample location of Chinook salmon. Individuals were also highlighted by Admixture group (individuals with ancestry values < 0.7 were represented by mixes of admixture groups with the largest contributor first). The x-axis represents the sampling site (see File S1 for full name) and the y-axis represents the total length of runs of homozygosity within each individual genome. Slide 2) same as slide 1 except for coho salmon. Slide 3) same as slide 1 except for sockeye salmon.

**Figure S8. Number of polymorphic loci for each sample location.** a) The number of polymorphic loci per location is shown on the top for all species. The number of samples per location is shown on the bottom. b) The relationship between the number of samples per location and the number of polymorphic loci identified for those locations. c) A boxplot of the number of polymorphic loci based on admixture group. Locations with average admixture group ancestry values ≥ 0.7 only have one label (i.e., lower Fraser River – LFR, mid Fraser River – MFR, and upper Fraser River – UFR). Those with < 0.7 have two labels with the first having the largest ancestry value. Only locations with at least four samples were used for this analysis. d) Relationship between the number of polymorphic loci and nucleotide diversity for locations with at least four individuals per site. Each species has its own linear regression line plotted.

**Figure S9. Extended haplotype homozygosity comparisons between LRF and MFR admixture groups.** a) A manhattan plot of -log10 p-values from an Rsb analysis between LFR and MFR sockeye salmon admixture groups. Sockeye salmon chromosomes were based on a draft genome assembly submitted to the NCBI (now GCA_034236695.1). Significant peaks were indicated on the graph with a vertical line and a number. b) Same as a) for Chinook salmon. c) Same as a) for coho salmon.

**File S1. Sample summary information in a spreadsheet format.**

**File S2. Truth SNPs used in GATK recalibration for each species (includes a readme file). File S3. Significant extended haplotype homozygosity from all comparisons among groups.**

