## Supplementary figures and images for "Revealing the evolutionary history and contemporary population structure of Pacific salmon in the Fraser River through genome resequencing"

### Figure S1

#### Sockeye salmon

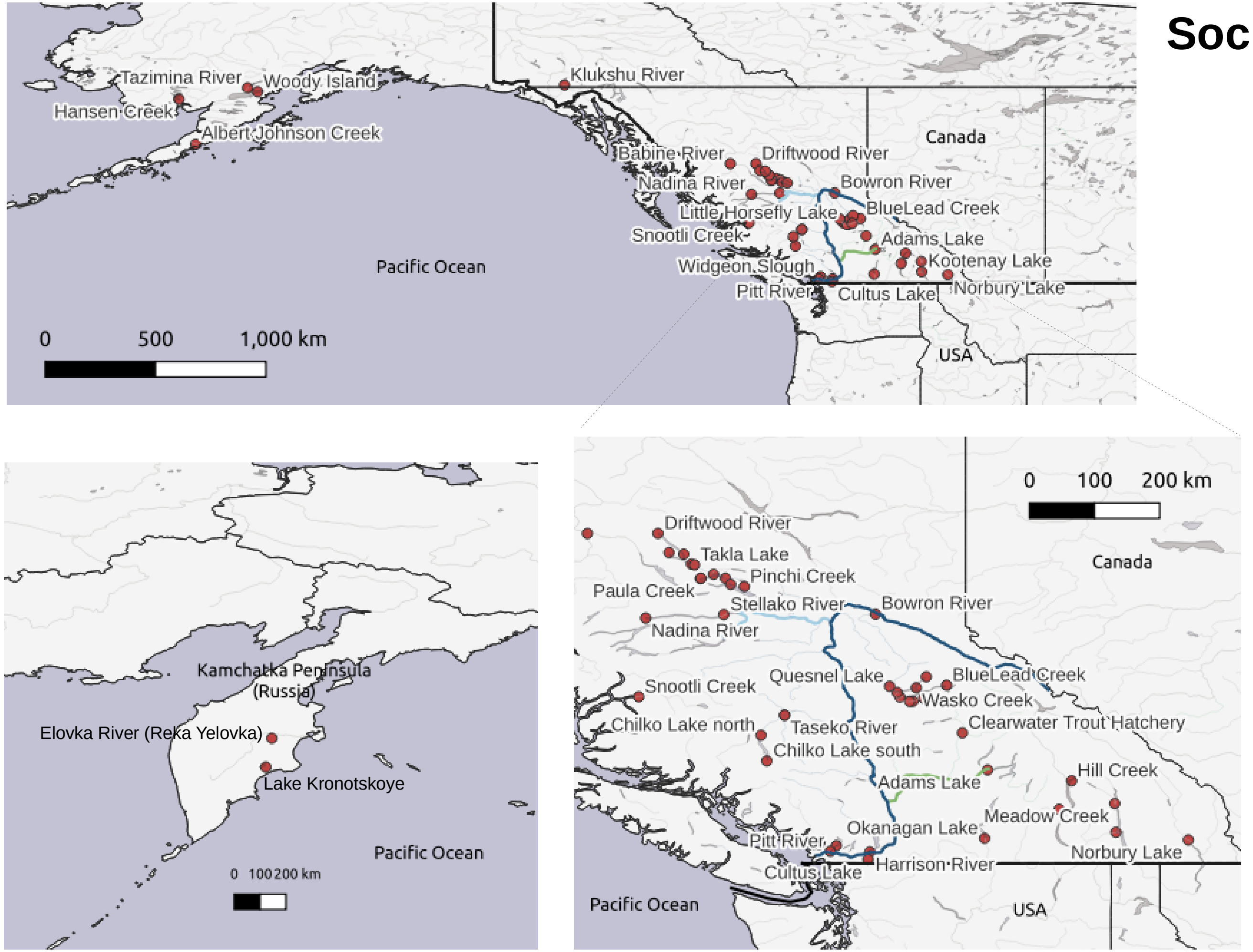

#### Chinook salmon

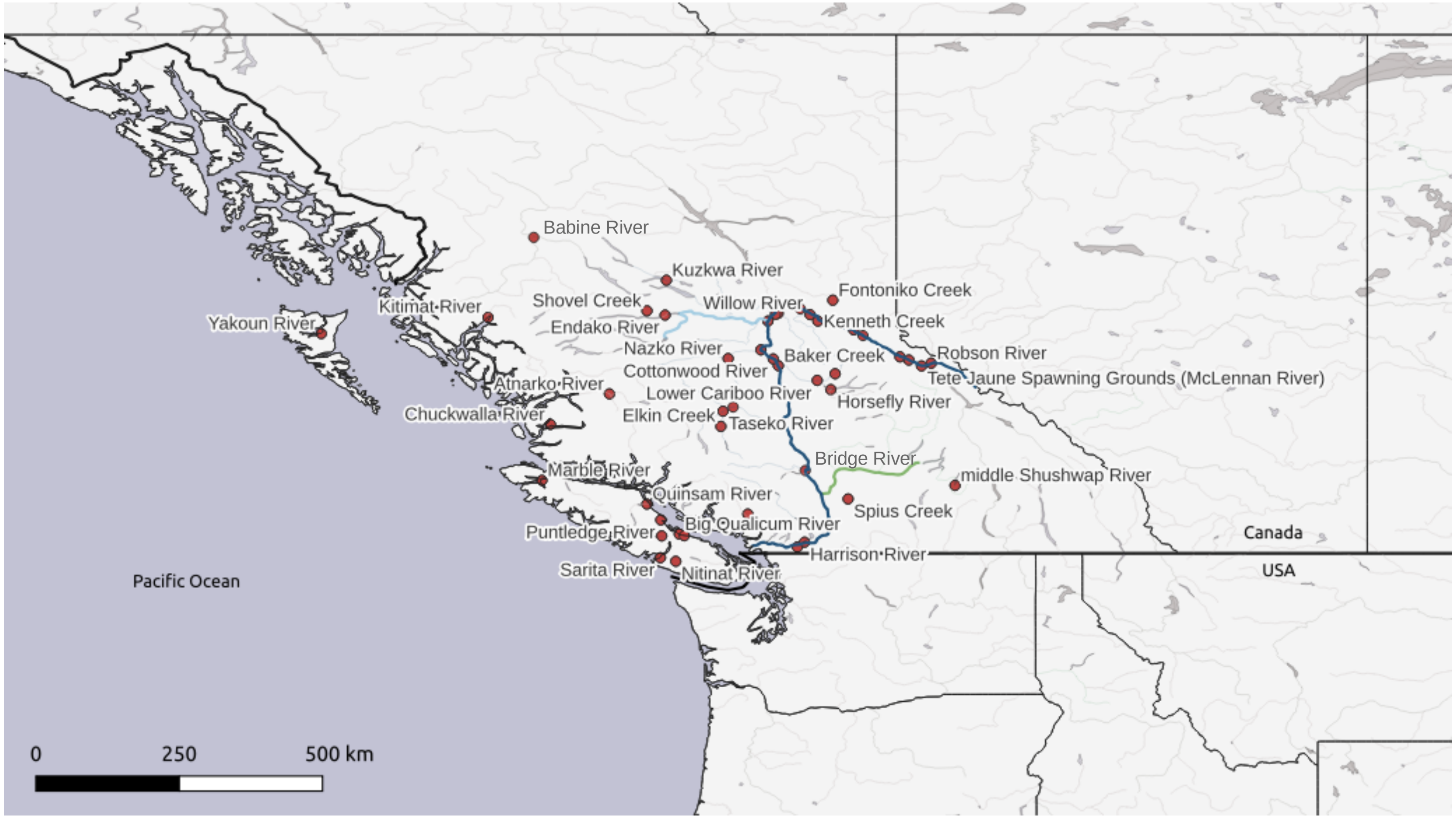

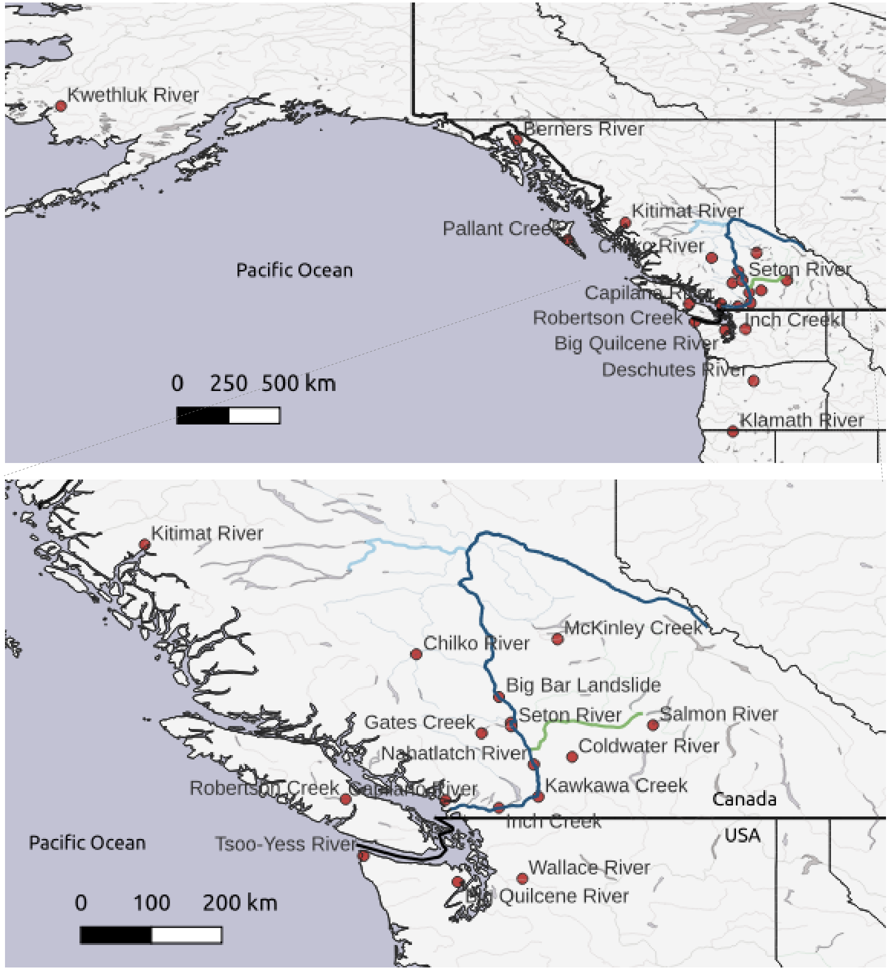

### Figure S2

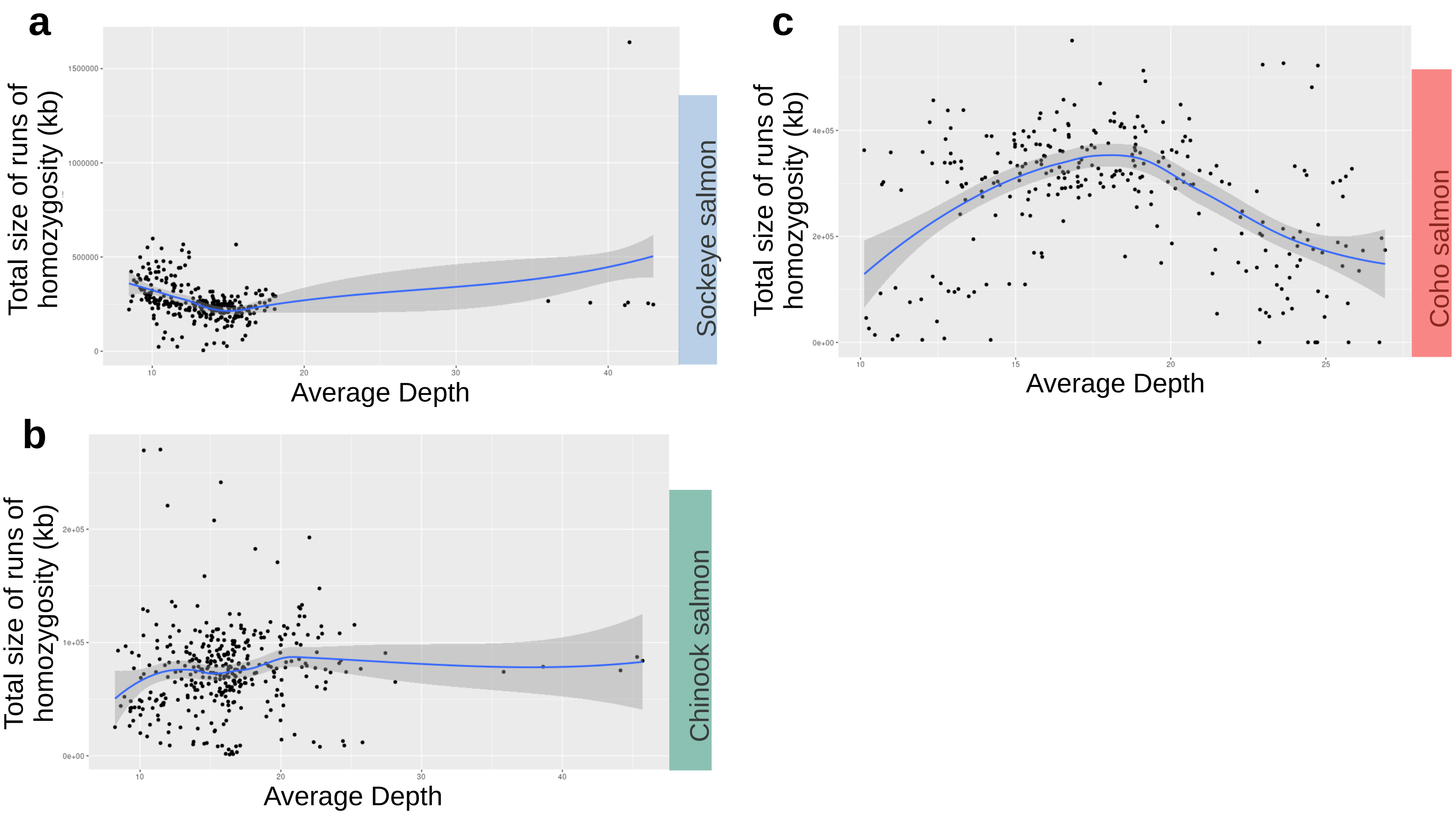

### Figure S3

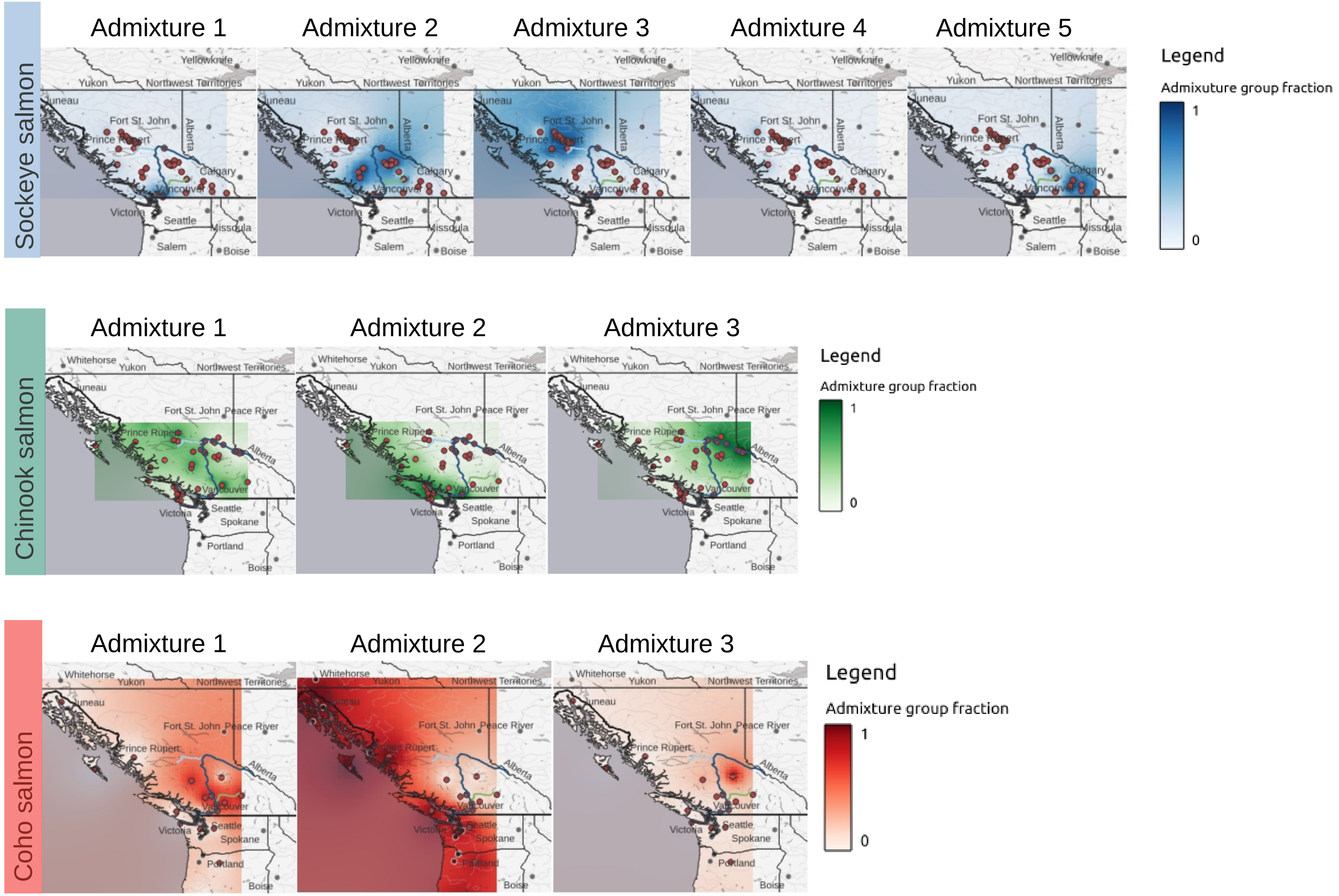

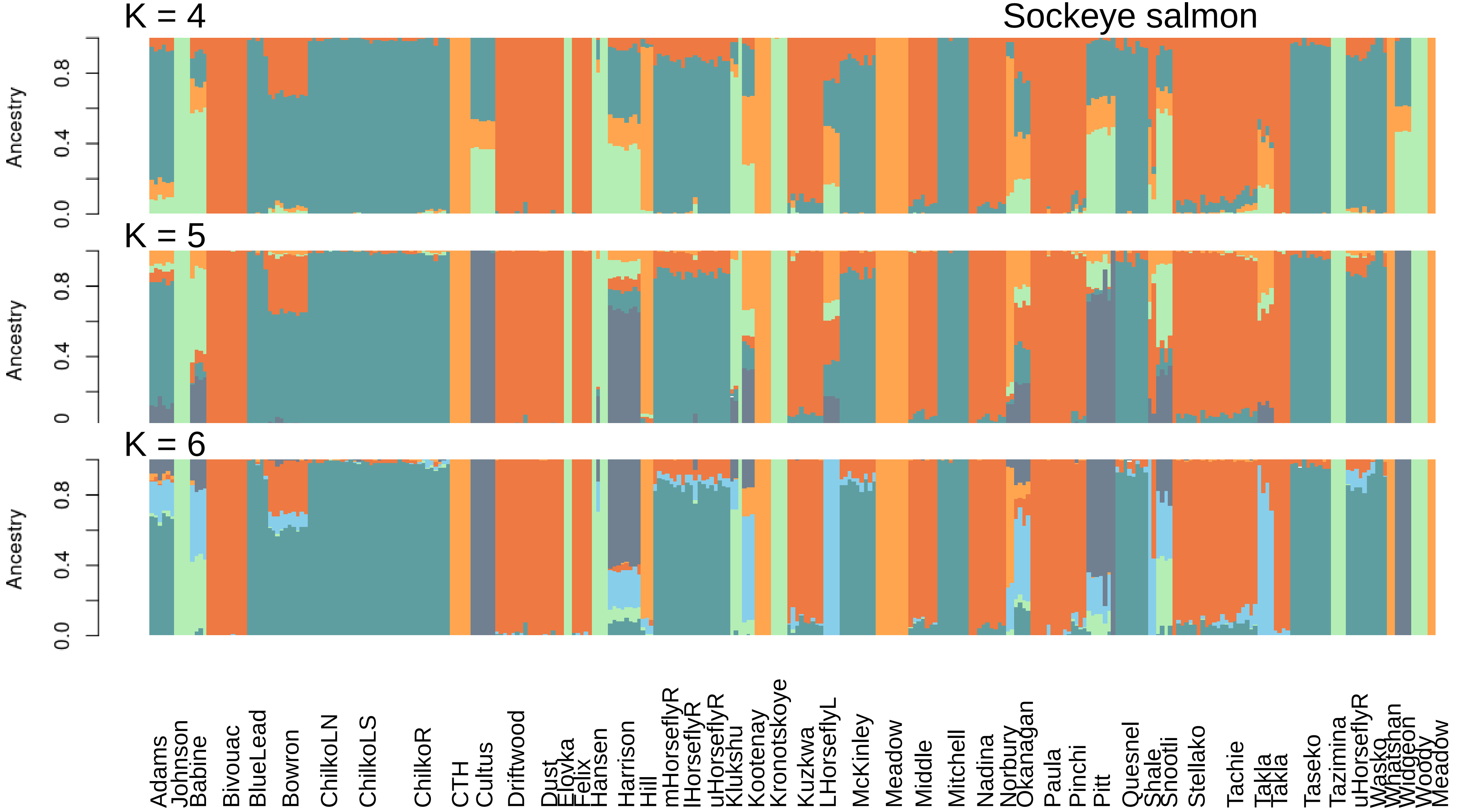

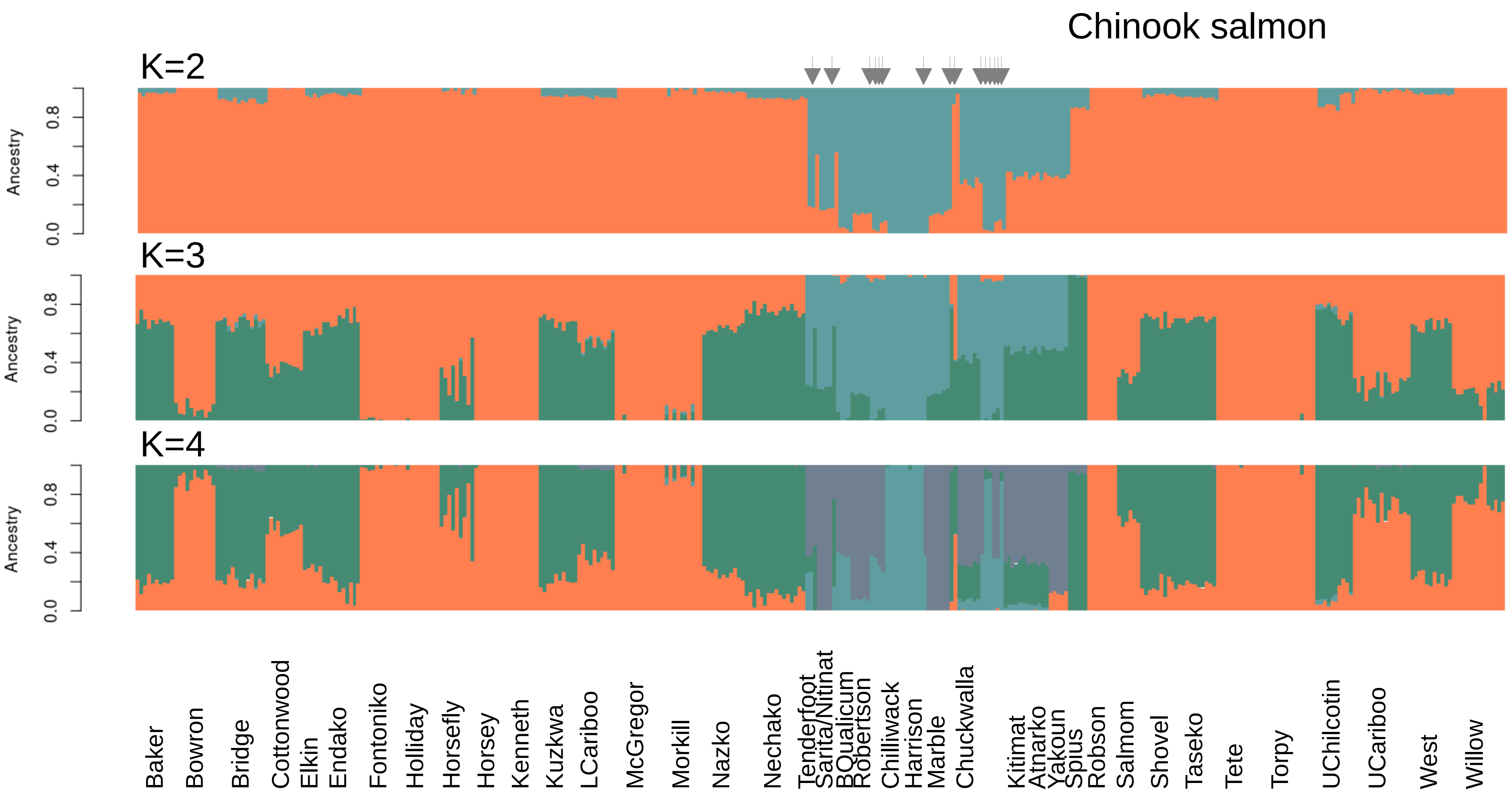

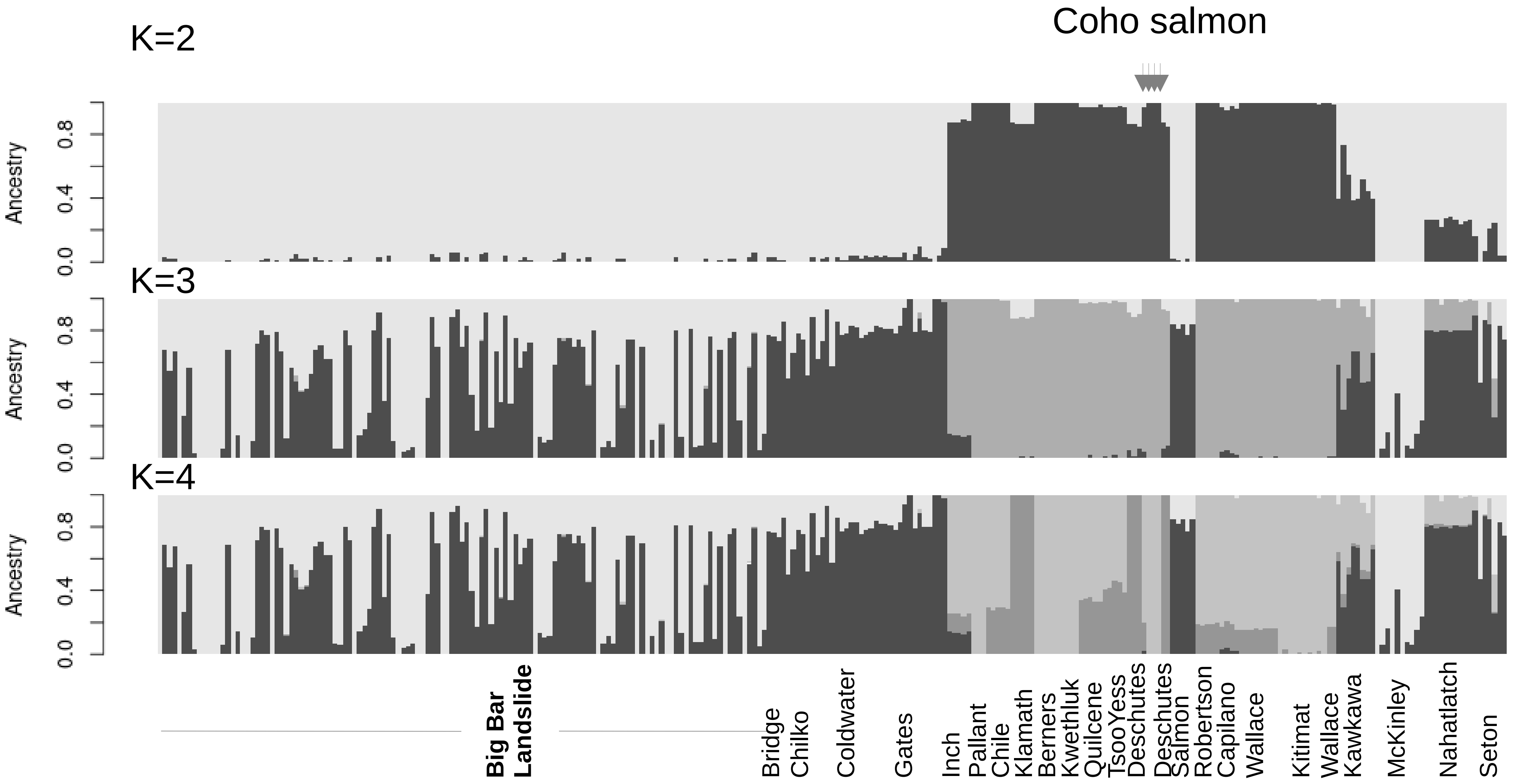

### Figure S4

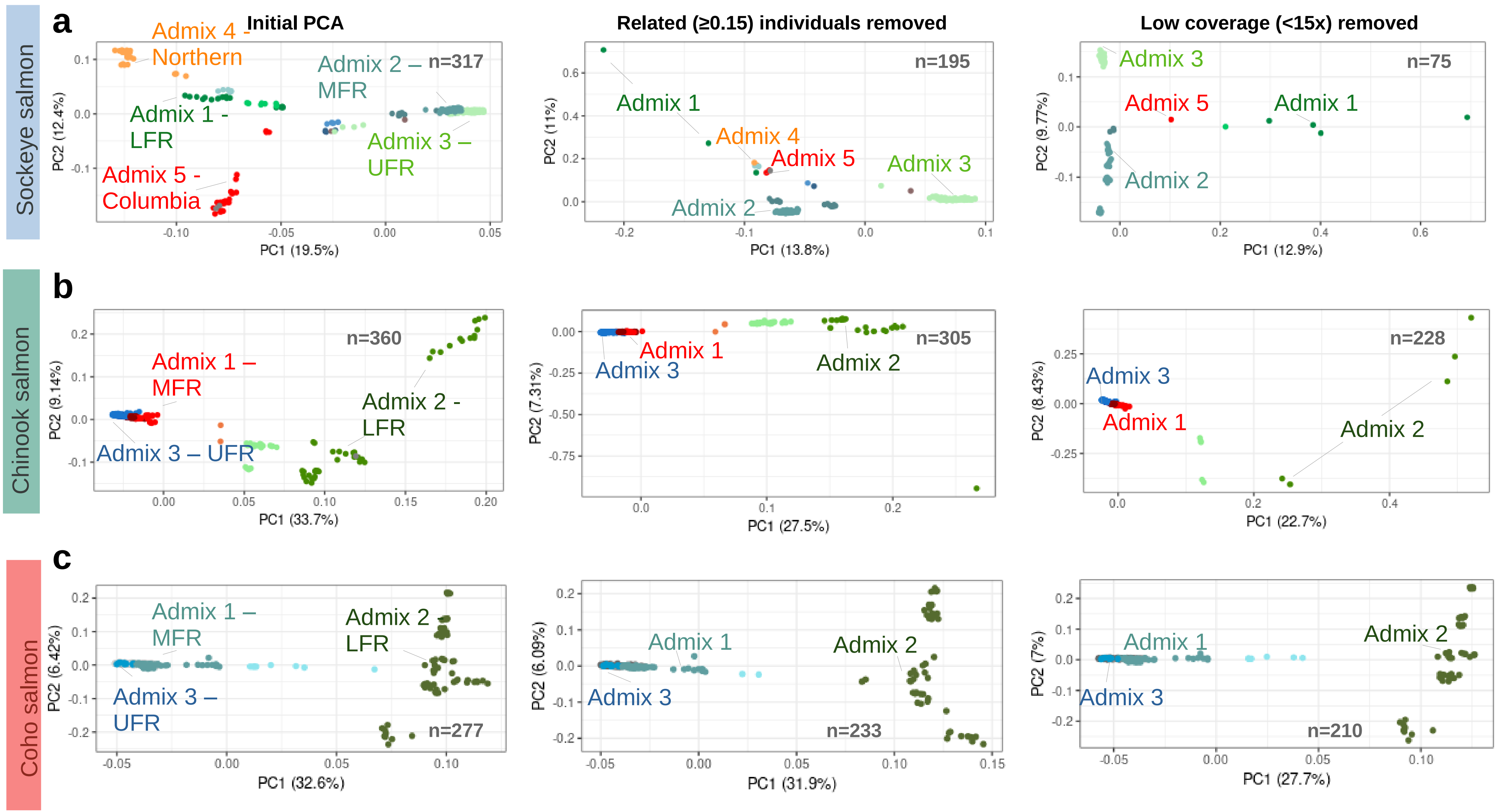

### Figure S6

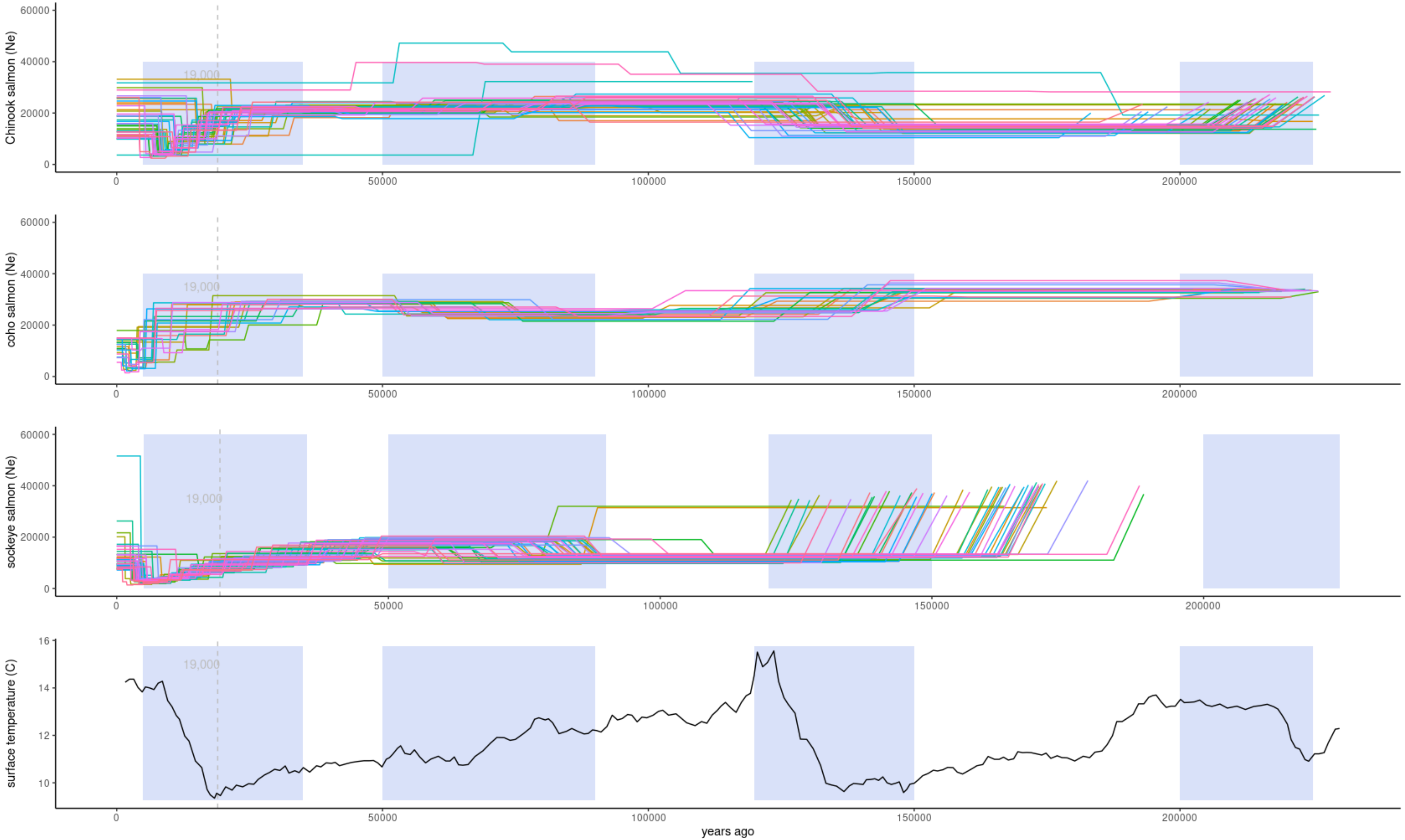

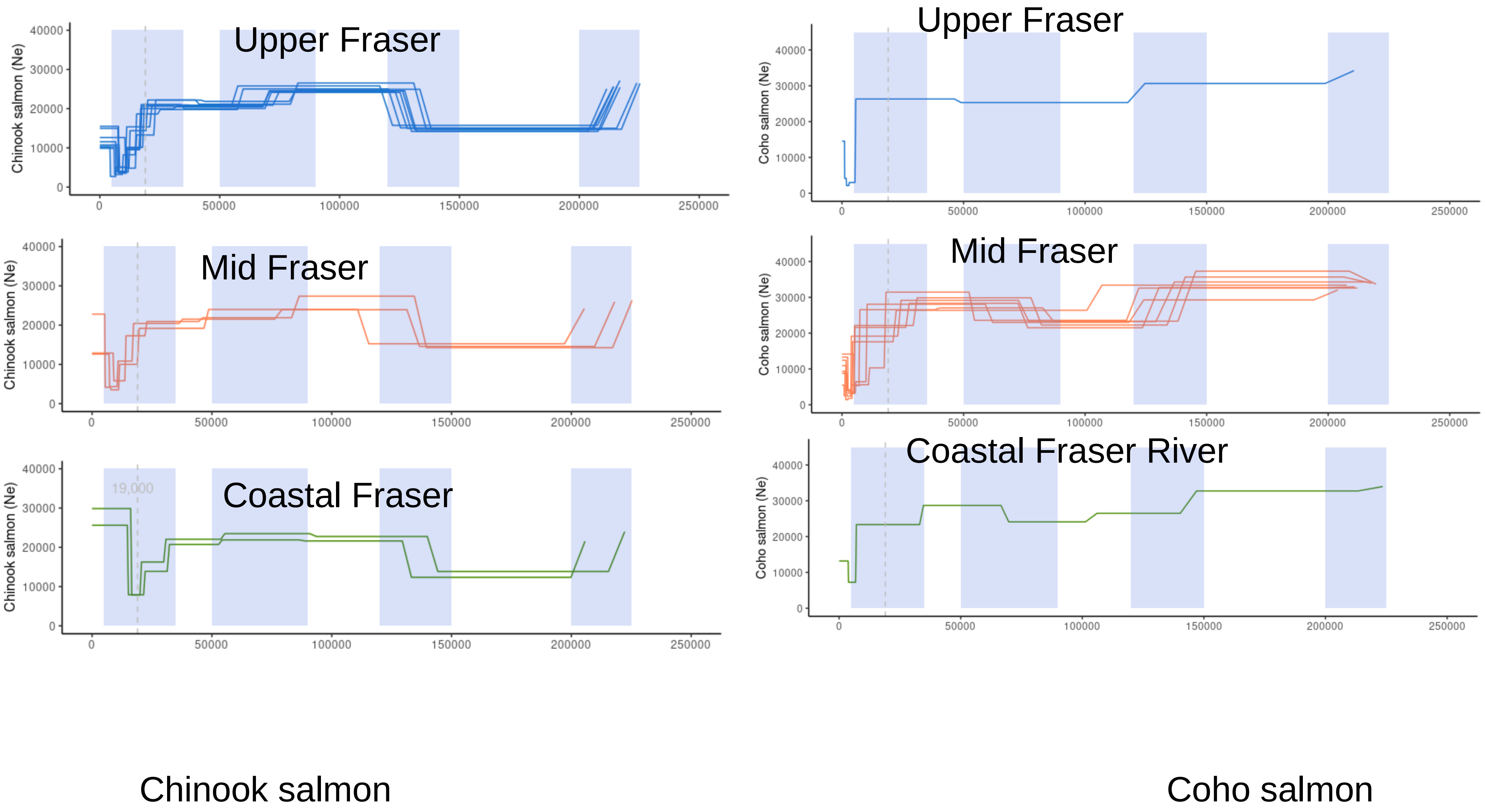

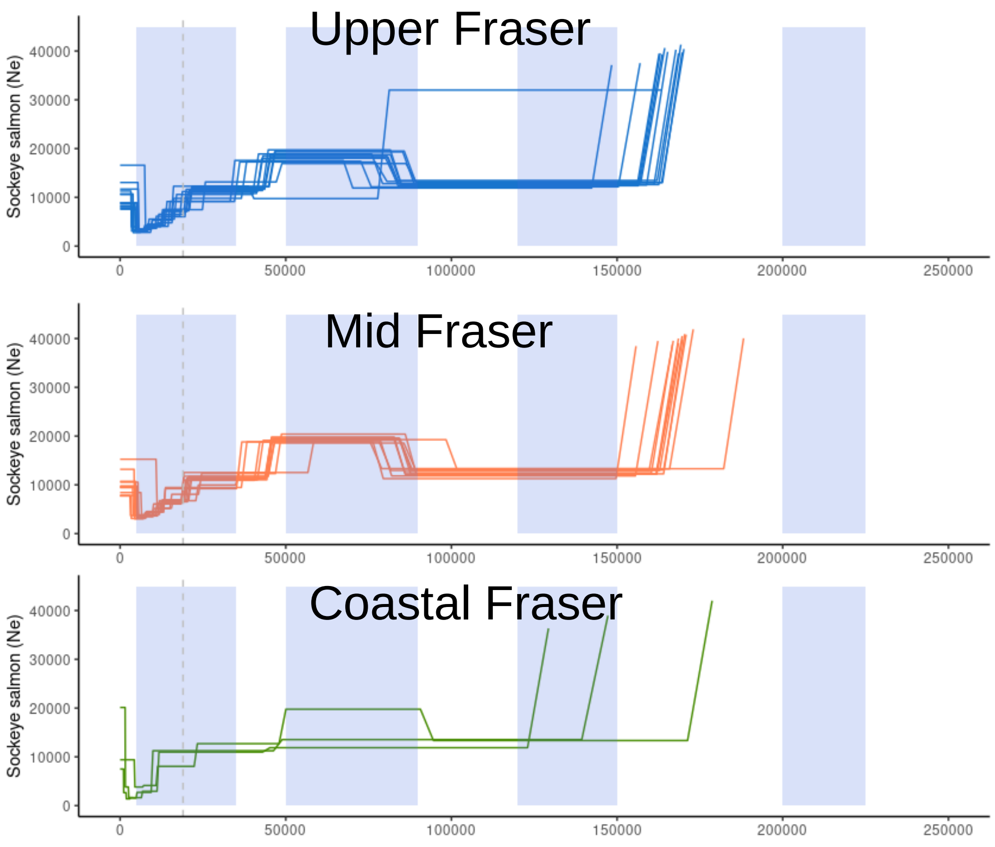


Sockeye salmon

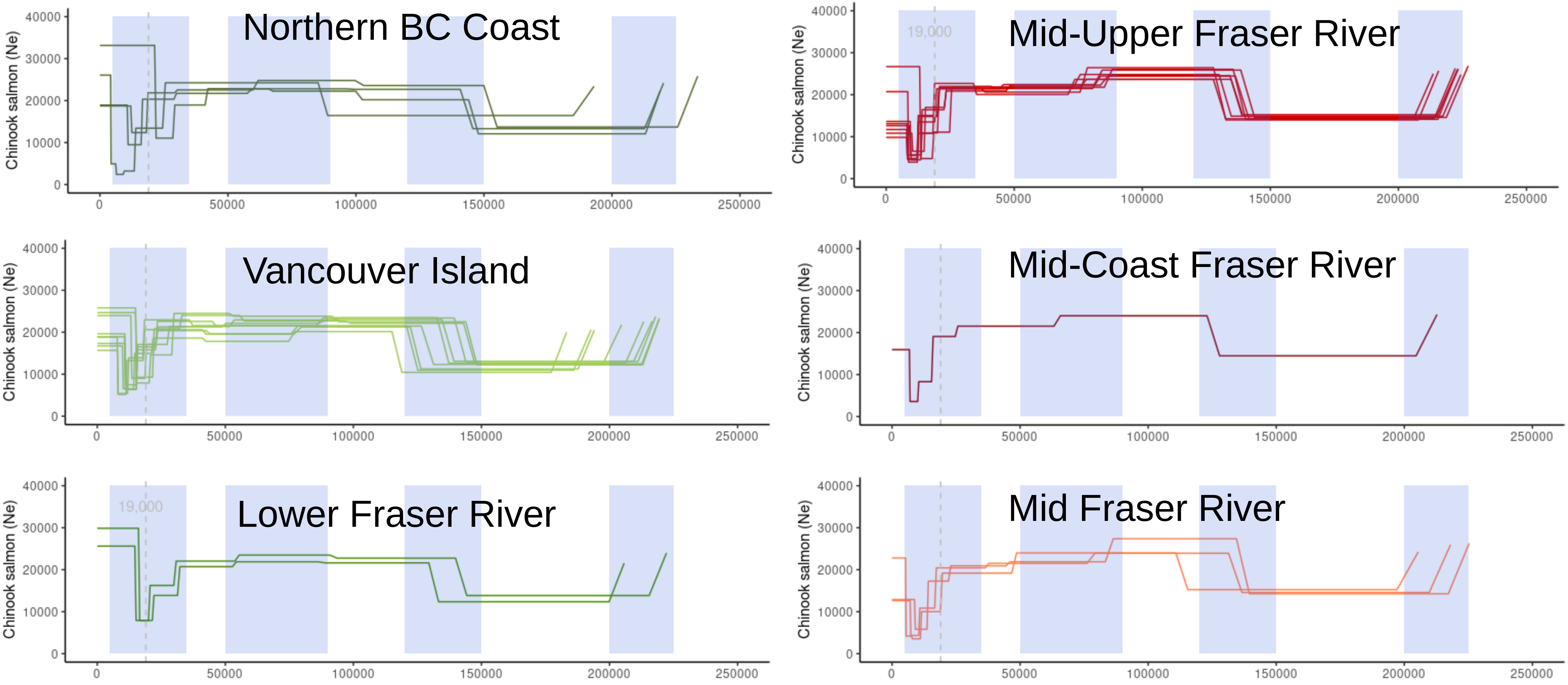


Chinook salmon

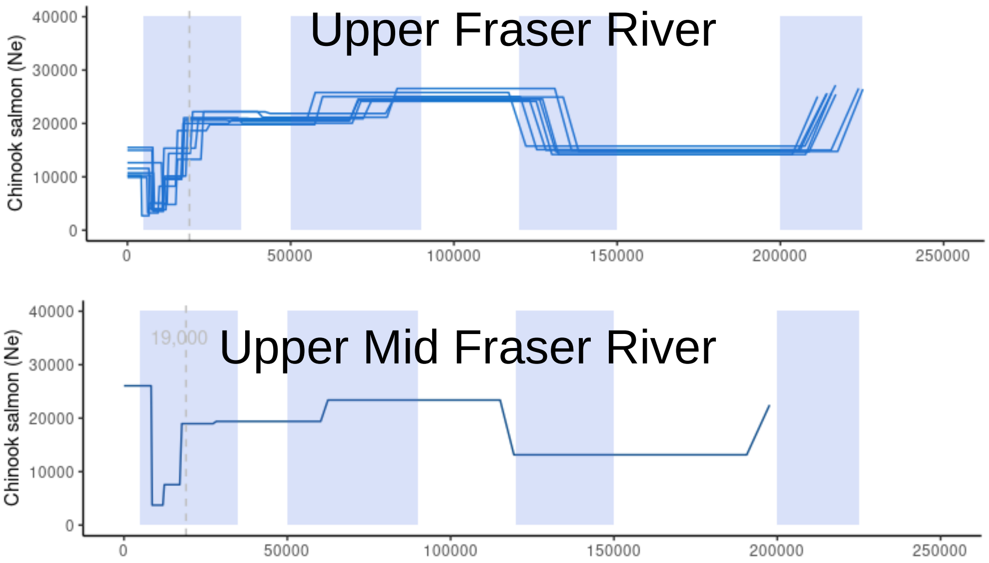


Chinook salmon

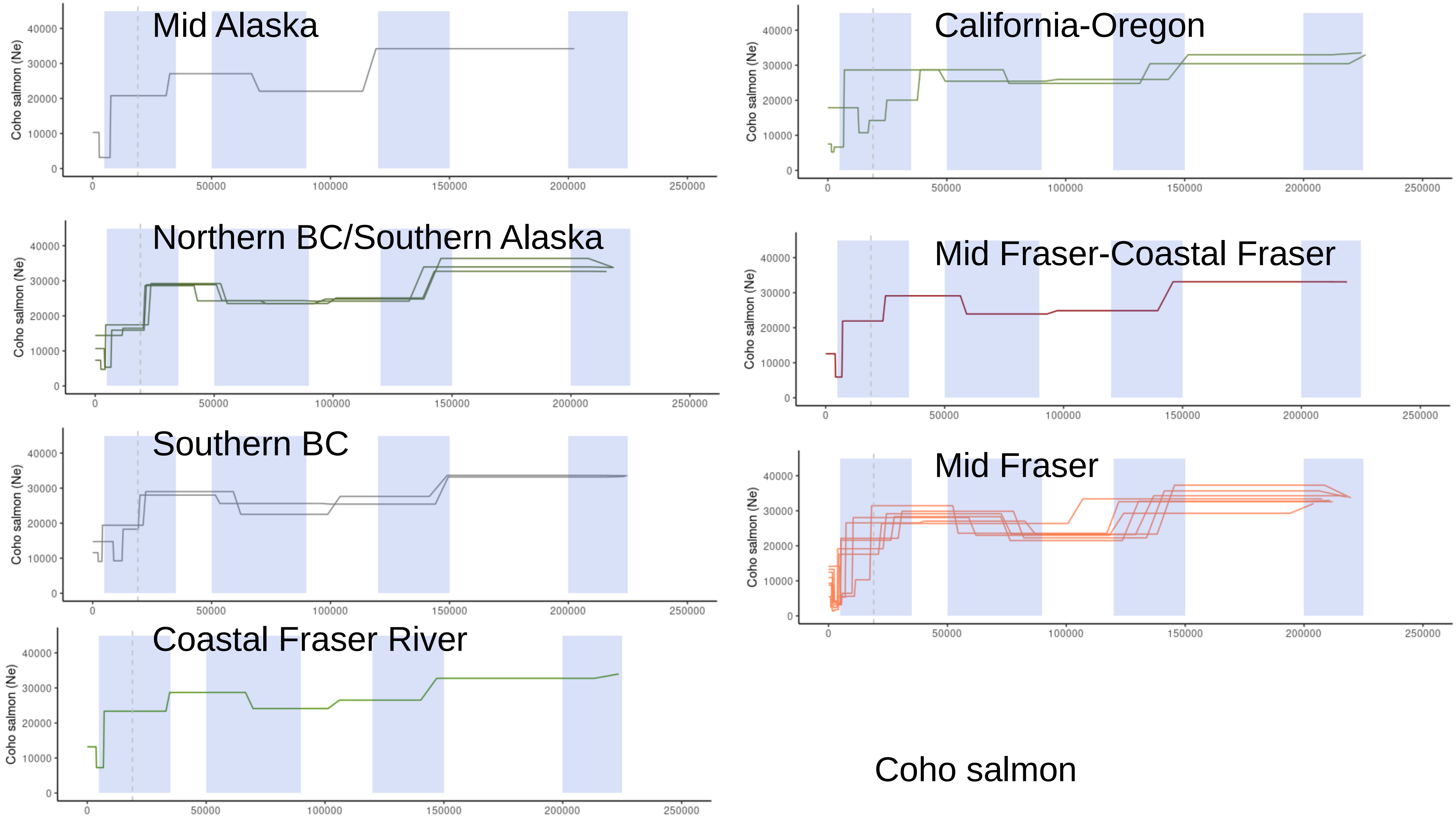

##### Upper Fraser

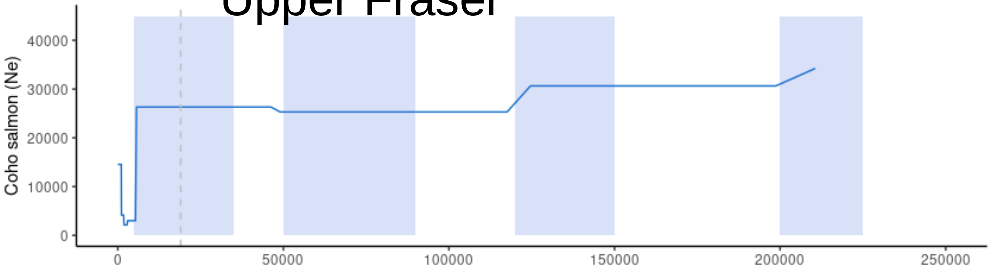


Coho salmon

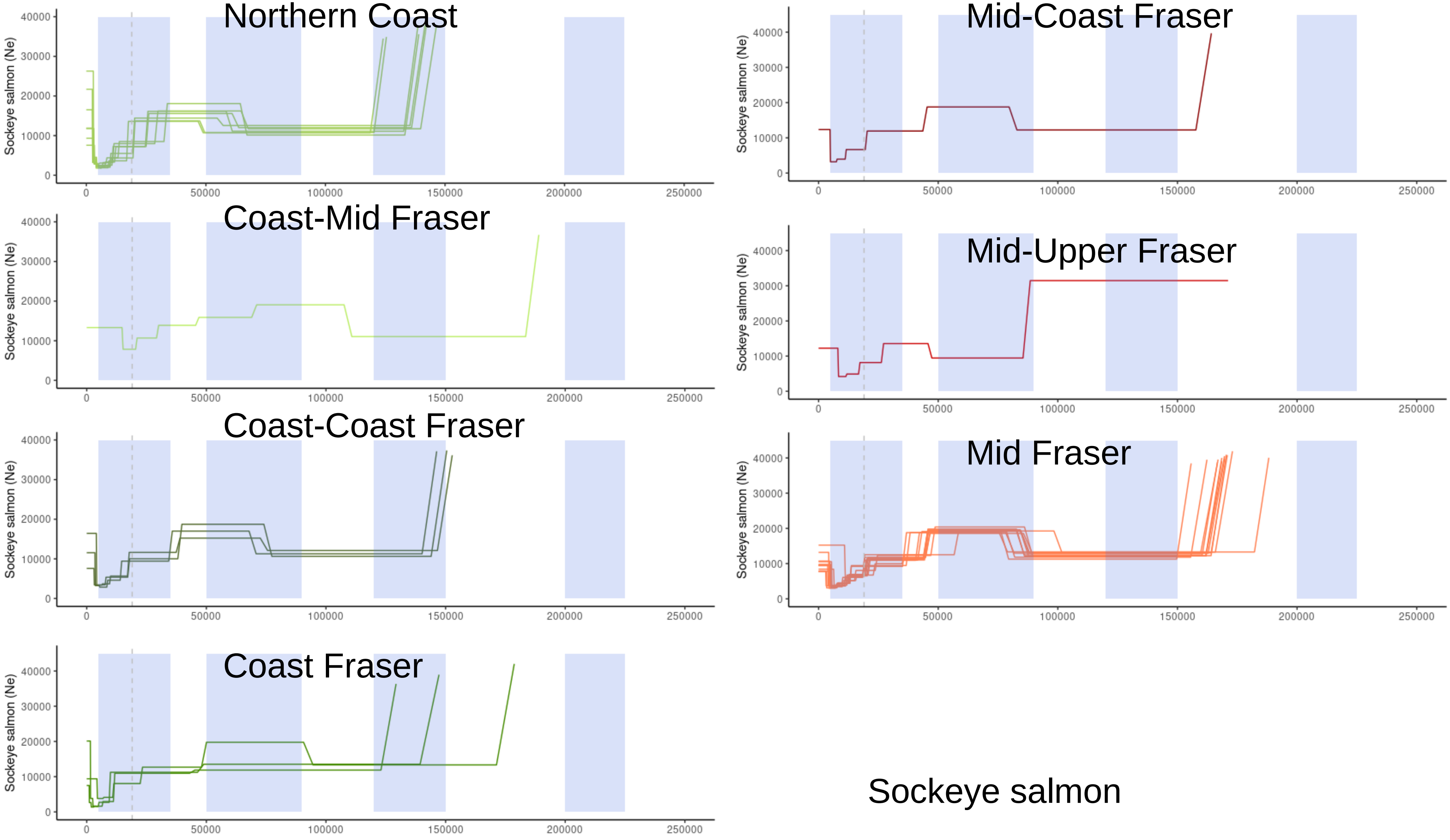

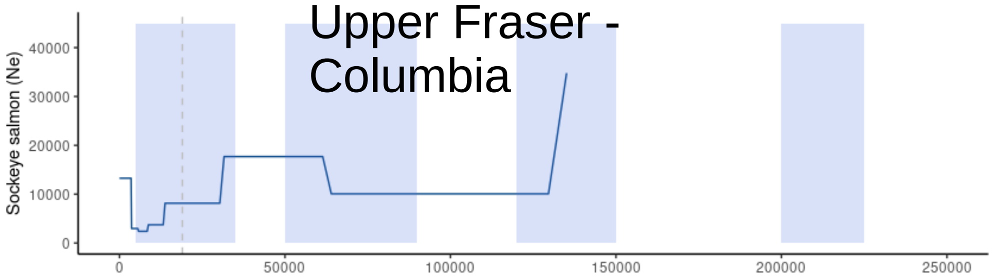


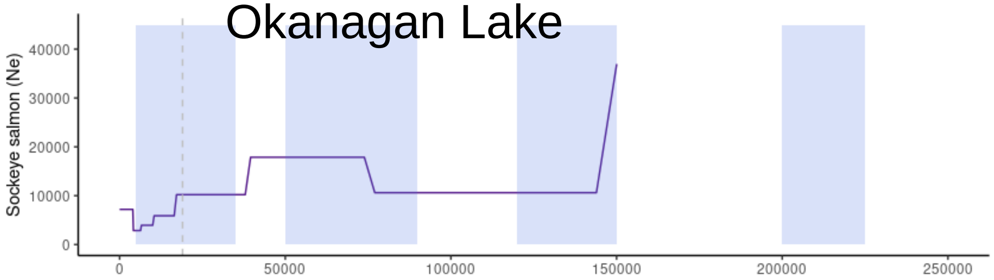


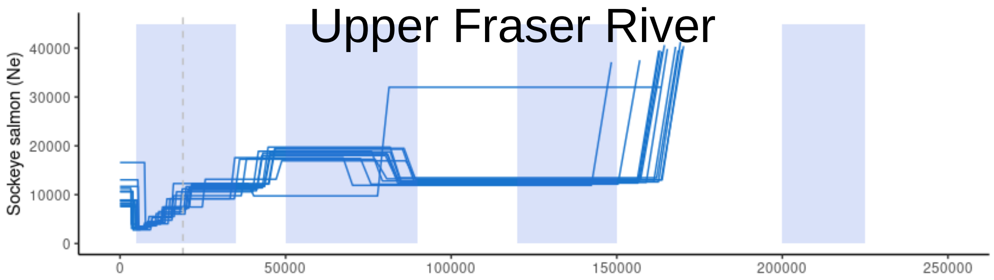


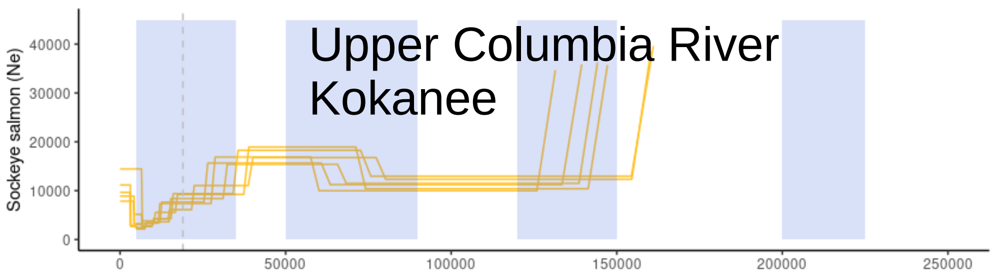


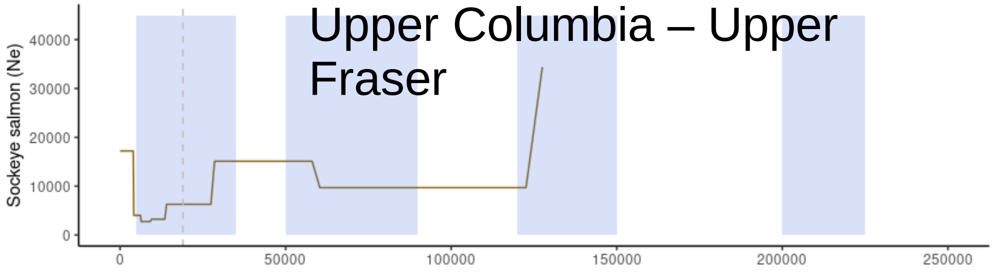


Sockeye salmon

### Figure S7

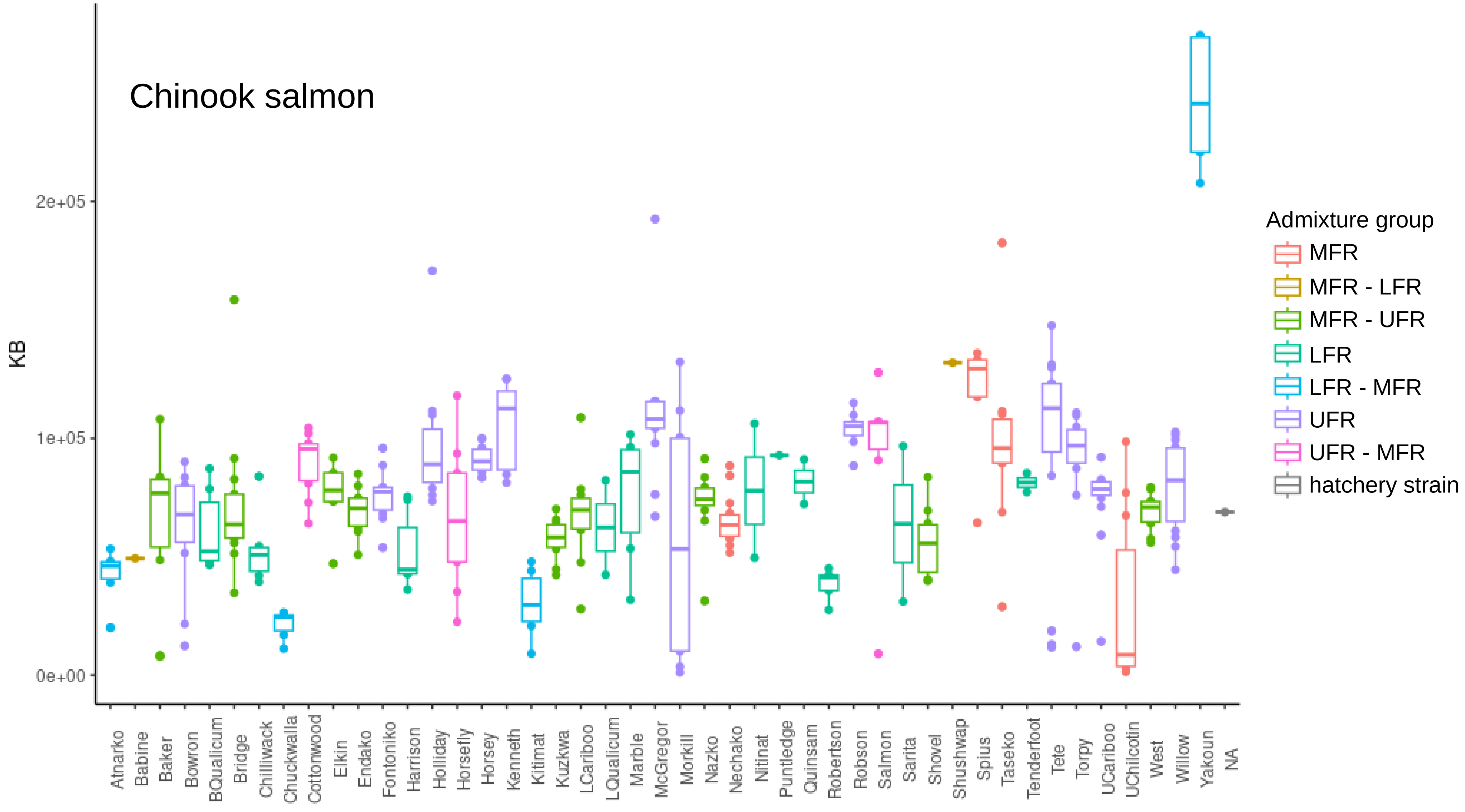

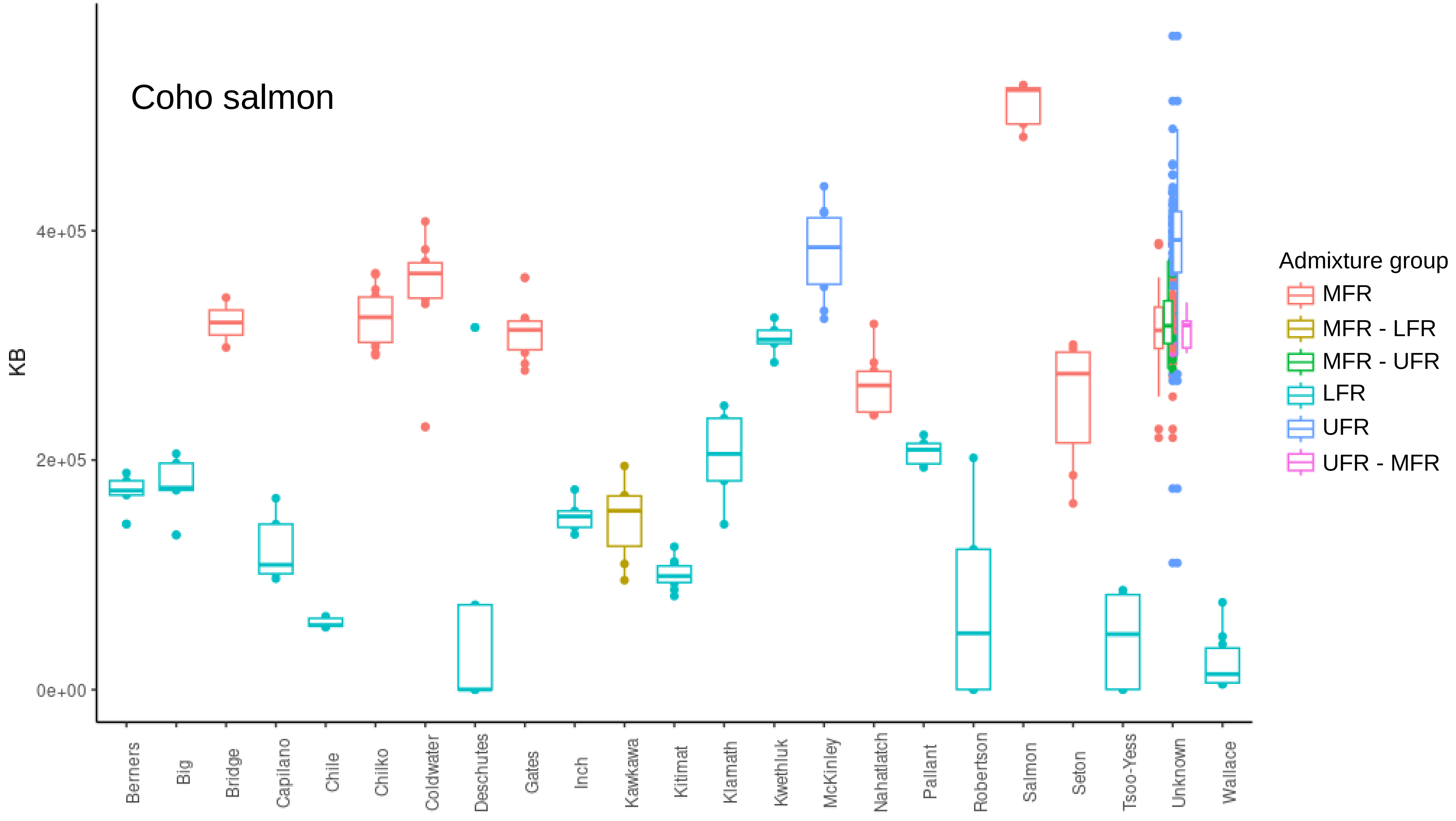

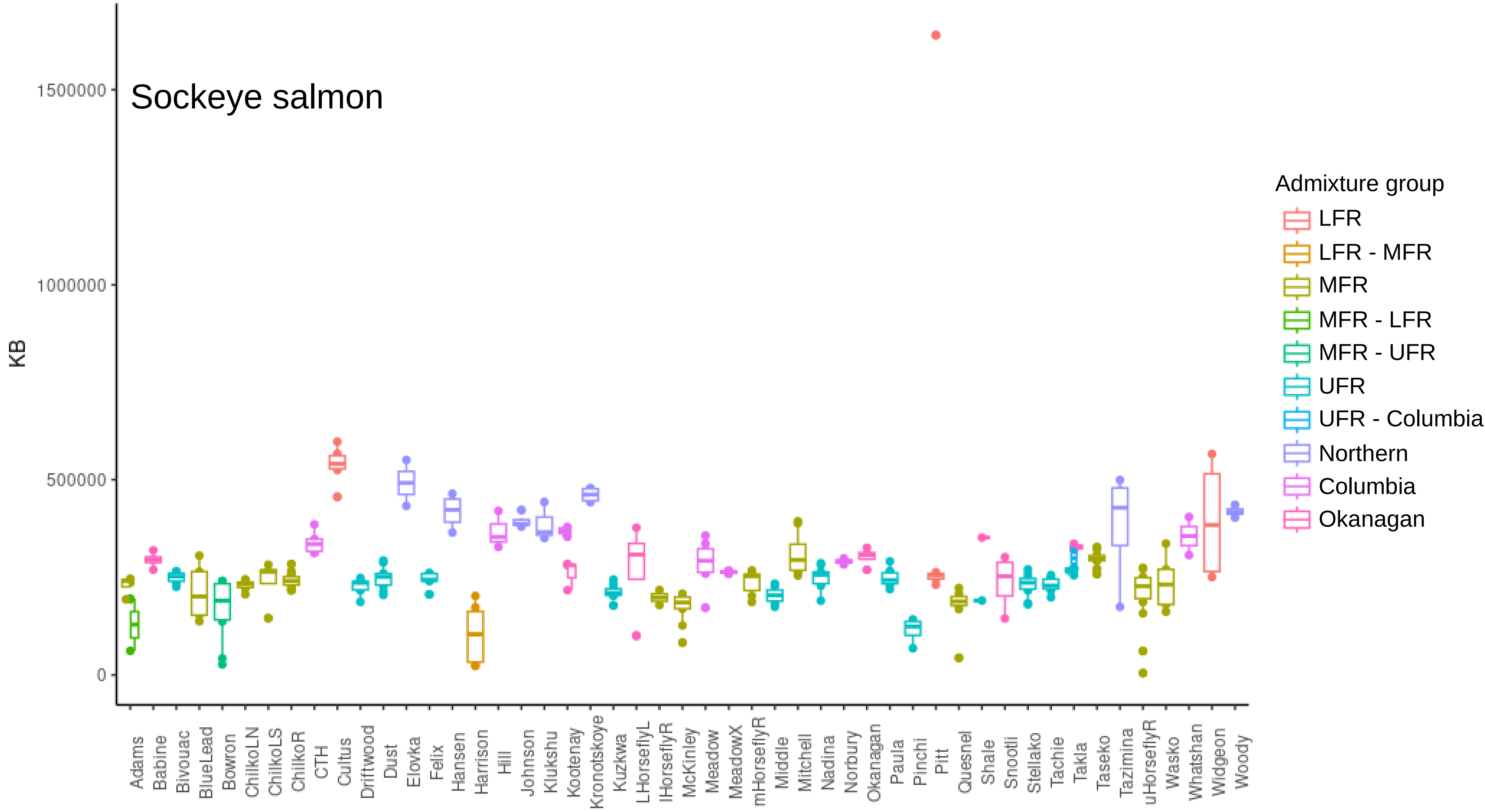

### Figure S8

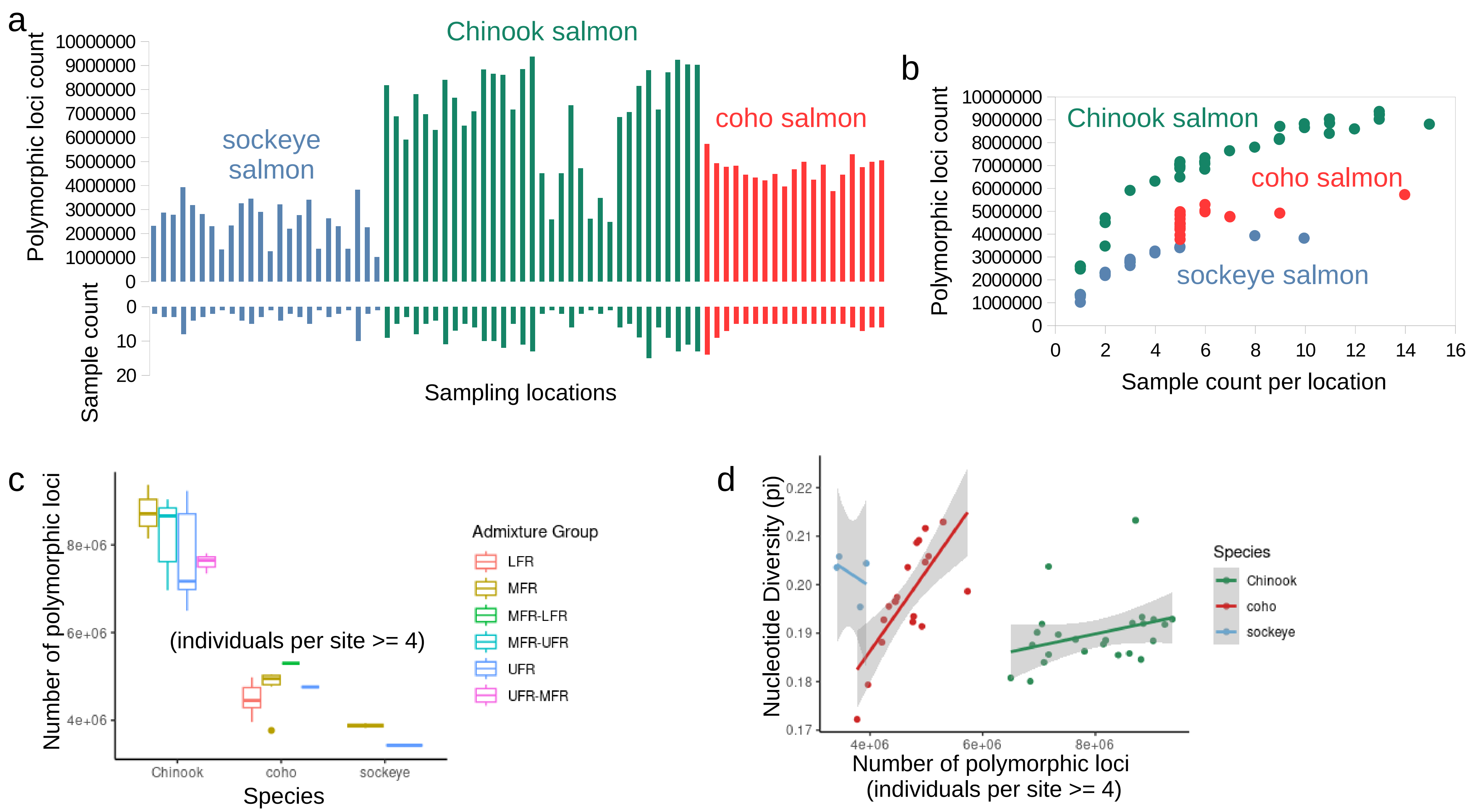

### Figure S9

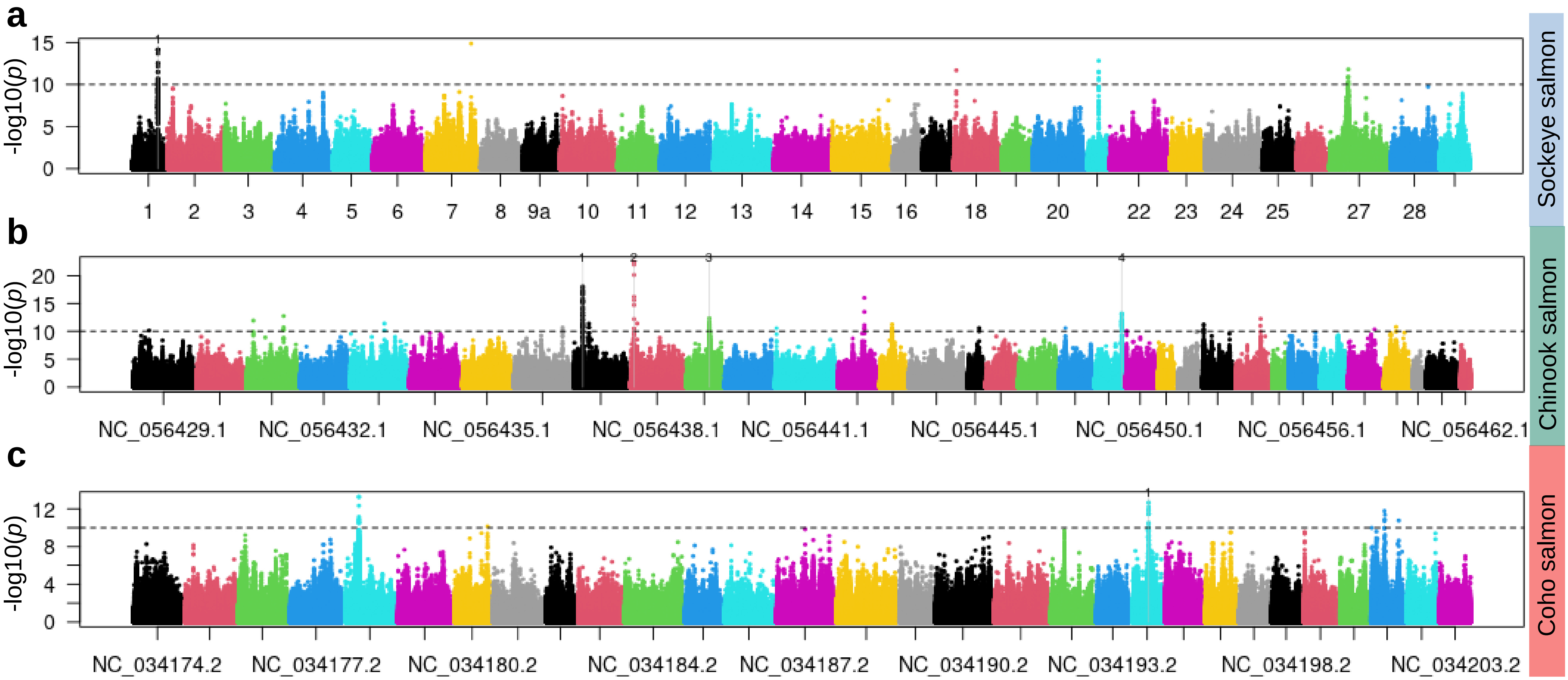
