## Supplementary material for "Revealing the evolutionary history and contemporary population structure of Pacific salmon in the Fraser River through genome resequencing": Figure S5

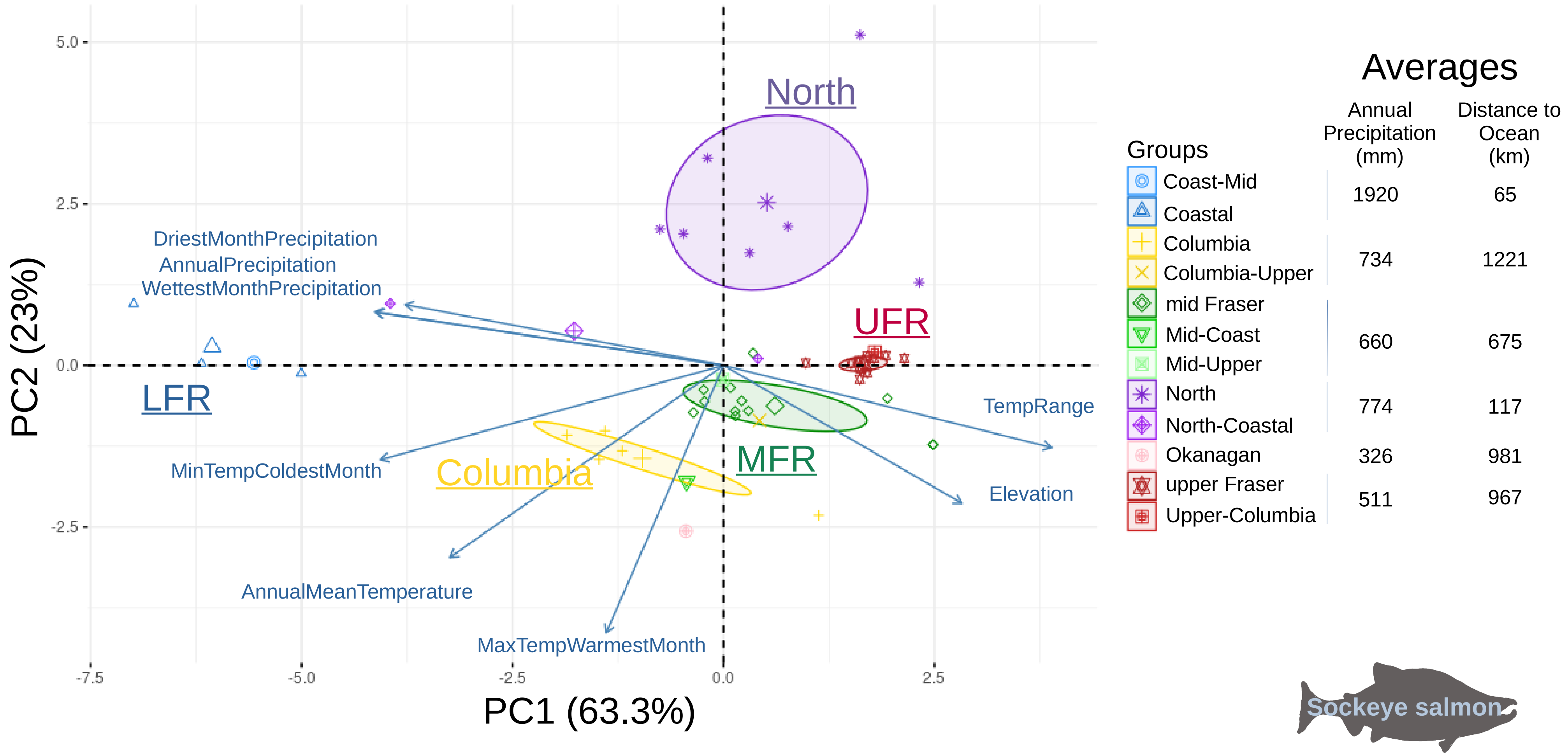
North
UFR
LFR
MFR
Columbia
DriestMonthPrecipitation
AnnualPrecipitation
WettestMonthPrecipitation
TempRange
Elevation
MinTempColdestMonth
AnnualMeanTemperature
MaxTempWarmestMonth
5.0
2.5
0.0
-2.5
-7.5
-5.0
-2.5
0.0
2.5
PC2 (23%)
PC1 (63.3%)
Averages
Groups
Annual Precipitation (mm)
Distance to Ocean (km)
Coast-Mid
Coastal
1920
65
Columbia
Columbia-Upper
734
1221
mid Fraser
Mid-Coast
Mid-Upper
660
675
North
North-Coastal
774
117
Okanagan
326
981
upper Fraser
Upper-Columbia
511
967
Sockeye salmon

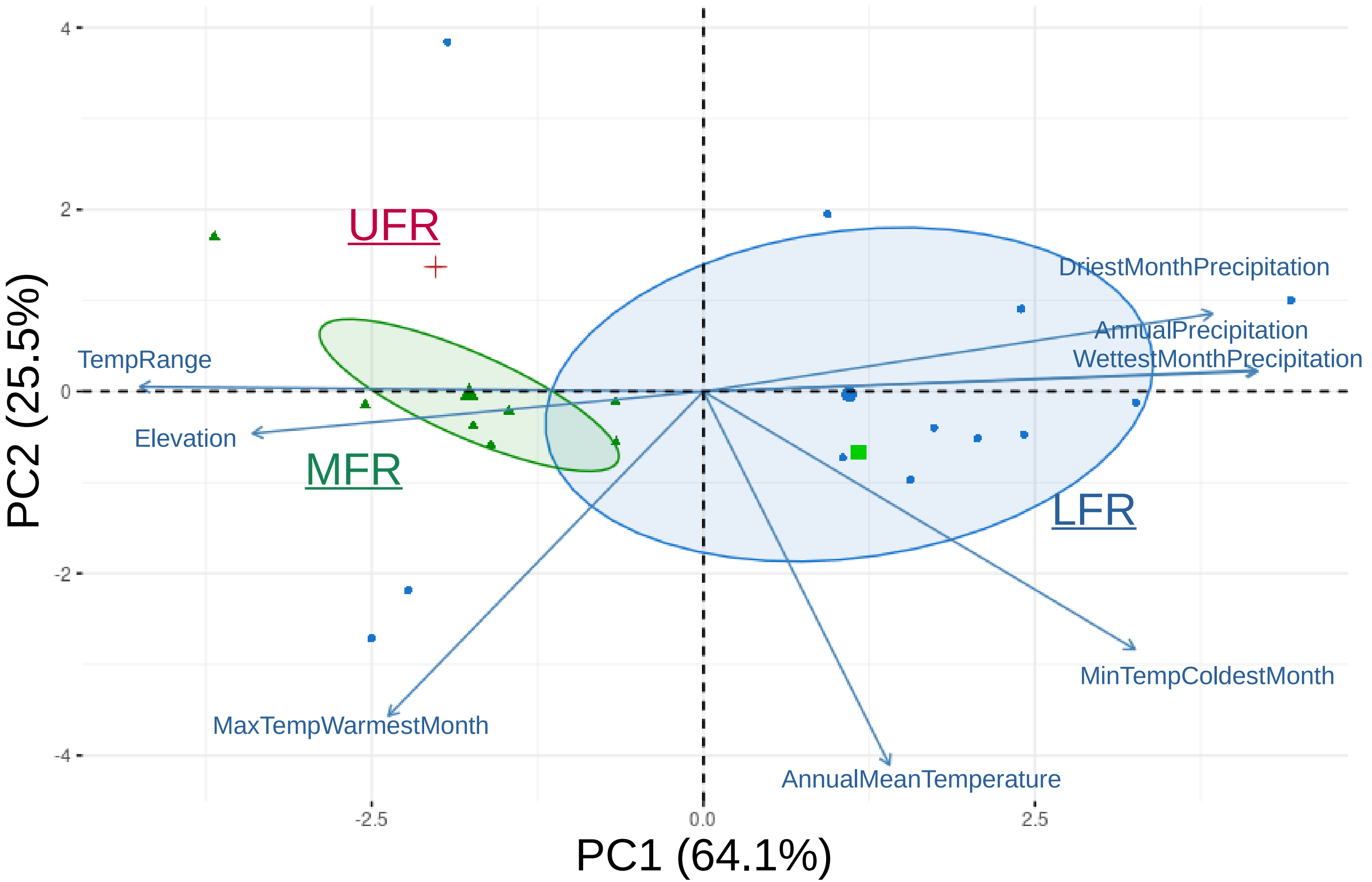


#### Averages

| Groups | Annual Precipitation (mm) | Distance to Ocean (km) |
|---|---|---|
| Coastal | 1598 | 84 |
| mid Fraser | 517 | 372 |
| Mid-Coast* | 1721 | 152 |
| upper Fraser* | 711 | 740 |

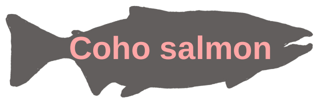

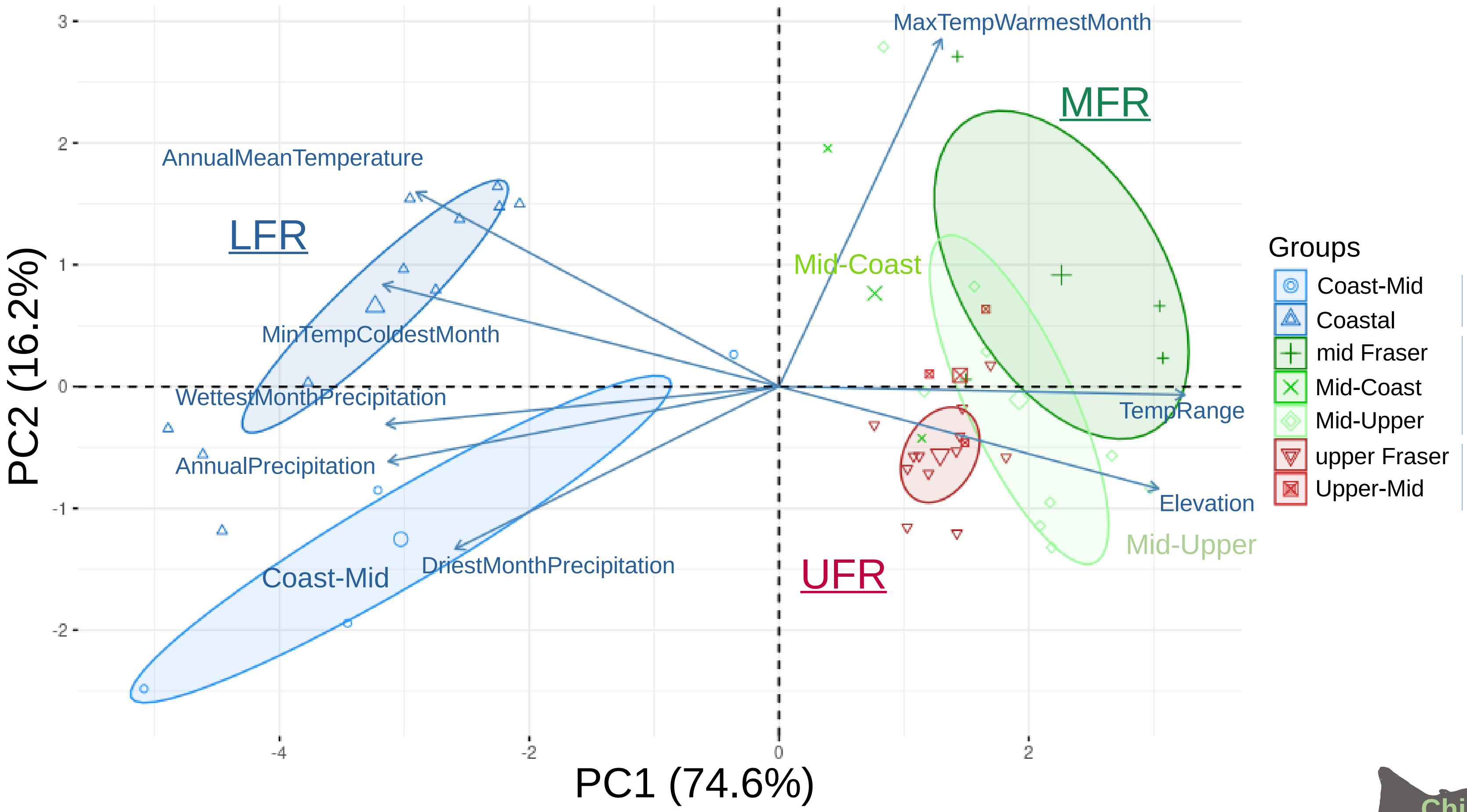


### Averages

| Groups | Annual Precipitation (mm) | Distance to Ocean (km) |
|---|---|---|
| Coast-Mid, Coastal | 1992 | 19 |
| mid Fraser, Mid-Coast, Mid-Upper | 499 | 607 |
| upper Fraser, Upper-Mid | 731 | 845 |

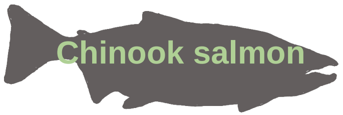
