## Supplementary material for "Revealing the evolutionary history and contemporary population structure of Pacific salmon in the Fraser River through genome resequencing": File S2: readme.txt

#############################################
### Truth Set  ##############################
#############################################

Brieuc et al. 3187 SNPs
Meeks et al.  821 SNPs
Total         4008 SNPs

	### Brieuc et al. 2014
	
		1) Identify truth data set
		
			Brieuc et al. 2014 (~7000 variants) 
			A Dense Linkage Map for Chinook salmon (Oncorhynchus tshawytscha) Reveals Variable Chromosomal Divergence After an Ancestral Whole Genome Duplication Event
			File S2 and S3
			F3 2.6 has the positions, but they look like they are 0-based instead of 1-based
			Manually created a vcf file so I could use a previous methodology
			
			$ samtools faidx Brieuc_Consensus_AllSequences.fasta ###(8127 sequences)

			####BWA mem doesn't handle sequences smaller than 70 bp, so I had to modify the script
			$ python3 ~/Desktop/Documents/General_Scripts/mapSNP2NewGenome/VCF_2_Fasta_v1.1_SpecificProject_OtshTruth.py \
			-vcf Brieuc_Consensus.vcf -fasta Brieuc_Consensus_AllSequences.fasta \
			-fai Brieuc_Consensus_AllSequences.fasta.fai -flanking 10 \
			> Brieuc_Consensus_AllSequences.FlankingSeqs.fasta
	
				Opened vcf file
					Finished reading vcf file: Found 8639 variants, used 8125
					
			$ bwa mem -M \
			/scratch2/shared/NCBI.Genomes/GCF_018296145.1_Otsh_v2.0_genomic.fna \
			Brieuc_Consensus_AllSequences.FlankingSeqs.fasta | \
			samtools sort \
			-o GCF_018296145.1.vs.Brieuc2014.bam
			
			(base) kchristensen@ccr:/scratch/kchristensen/Otsh/BCSRIF/truth$ samtools index GCF_018296145.1.vs.Brieuc2014.bam
			
			####BWA mem doesn't handle sequences smaller than 70 bp, so I had to modify the script
			$ python Bam2SNPPosition.v1.1_SpecificProject_OtshTruth.py \
			-file GCF_018296145.1.vs.Brieuc2014.bam \
			-qual 40 \
			-multiple 1 \
			-minLen 69 \
			-minPid 90 \
			-pos 11 > \
			GCF_018296145.1.vs.Brieuc2014.positions.txt
			
			$ python CheckMappingVCF.v1.1.py \
			-map GCF_018296145.1.vs.Brieuc2014.positions.txt \
			-fasta1 Brieuc_Consensus_AllSequences.FlankingSeqs.fasta \
			-fai1 Brieuc_Consensus_AllSequences.FlankingSeqs.fasta.fai \
			-fasta2 GCF_018296145.1_Otsh_v2.0_genomic.fna \
			-fai2 GCF_018296145.1_Otsh_v2.0_genomic.fna.fai \
			-flanking 5 \
			-output most > \
			GCF_018296145.1.vs.Brieuc2014.positions.QCfiltered.txt
			
			Opened map file
				Finished reading vcf file: Found 7238 variants, correct variants:2466, unknown variants:655, CompleteMismatch:3253, variantOnlyMismatch:792, partialMatch:72
			
			###Manually checked for markers that didn't match linkage group
			###Had to manually remove part of the name in the Brieuc2Otsh.NCBIv2.positions.custom.QCfiltered.txt file that was added by the VCF_2_Fasta script
			
			###Map reads to new vcf file
			$ python3 ~/Desktop/Documents/General_Scripts/mapSNP2NewGenome/Assign_New_SNP_Positions.V1.1.py \
			-map GCF_018296145.1.vs.Brieuc2014.positions.QCfiltered.txt \
			-vcf Brieuc_Consensus.vcf \
			-fai GCF_018296145.1_Otsh_v2.0_genomic.fna.fai > \
			Brieuc_GCF_018296145.vcf

			(3187 sites)
			
			###Sort vcf file
			$ bcftools sort Brieuc_GCF_018296145.vcf > Brieuc_GCF_018296145.sorted.vcf
			
			###Index
			$ bgzip Brieuc_NCBI.v2.sorted.vcf 
			$ tabix Brieuc_NCBI.v2.sorted.vcf.gz 
	
	### Meeks et al (Sequencing improves our ability to study threatened migratory species: Genetic population assignment in California’s Central Valley Chinook salmon)
	
		1) Get the data in a format I can use other standard scripts with (those used from Brieuc et al. 2014 at least):
		
			Meeks/doi_10.5061_dryad.j29f5__v1$ \
			python3 GetVariantPostions.Meeks.v1.0.py \
			-fasta Meek_et_al_EandE_Sequence_Data.txt > \
			Meeks_et_al.FlankingSeqs.fasta 2> \
			Meeks_et_al.vcf
			
			###Added information to the Meeks_et_al.vcf file to match a vcf file (Header information)
			
			###Filter for markers on the genetic map (2208 markers left)
			
			Meeks/doi_10.5061_dryad.j29f5__v1$ vcftools --vcf Meeks_et_al.vcf --recode --recode-INFO-all --out Meeks_et_all.GeneticMapOnly --chr R021914 --chr R011144 --chr R009188 --chr R021729 --chr R014732 --chr R014190 --chr R019338 --chr R002309 \
			--chr R000853 --chr R011597 --chr R021560 --chr R010154 --chr R013428 --chr R002477 --chr R018209 --chr R006225 --chr R016045 --chr R018776 \
			--chr R007812 --chr R000069 --chr R018367 --chr R002660 --chr R011342 --chr R001230 --chr R014958 --chr R001300 --chr R013183 --chr R011452 \
			--chr R015187 --chr R001836 --chr R020745 --chr R005088 --chr R002804 --chr R014013 --chr R009295 --chr R002943 --chr R015080 --chr R000001 \
			--chr R005557 --chr R005873 --chr R017079 --chr R002471 --chr R012681 --chr R001338 --chr R000330 --chr R008843 --chr R001874 --chr R016333 \
			--chr R007017 --chr R009230 --chr R011853 --chr R015473 --chr R019475 --chr R006553 --chr R023626 --chr R013347 --chr R001359 --chr R016682 \
			--chr R017501 --chr R004058 --chr R004396 --chr R011849 --chr R013492 --chr R016850 --chr R000607 --chr R006508 --chr R021582 --chr R000805 \
			--chr R022127 --chr R021278 --chr R021212 --chr R006335 --chr R021970 --chr R011466 --chr R013787 --chr R016455 --chr R008499 --chr R000859 \
			--chr R013290 --chr R009968 --chr R019090 --chr R012007 --chr R005000 --chr R002653 --chr R000574 --chr R004352 --chr R011392 --chr R002105 \
			--chr R004635 --chr R008595 --chr R004449 --chr R021912 --chr R023271 --chr R011550 --chr R009355 --chr R013034 --chr R007827 --chr R013794 \
			--chr R020771 --chr R023652 --chr R004939 --chr R017265 --chr R000322 --chr R007555 --chr R005172 --chr R004188 --chr R013232 --chr R003855 \
			--chr R005159 --chr R008795 --chr R008919 --chr R017235 --chr R017911 --chr R015938 --chr R000475 --chr R002511 --chr R000056 --chr R000783 \
			--chr R018575 --chr R009143 --chr R013061 --chr R008236 --chr R009698 --chr R021148 --chr R003994 --chr R014822 --chr R004751 --chr R017351 \
			--chr R003941 --chr R007981 --chr R001857 --chr R010971 --chr R017397 --chr R015521 --chr R020478 --chr R010755 --chr R010082 --chr R006338 \
			--chr R003420 --chr R000395 --chr R009145 --chr R001559 --chr R009755 --chr R007503 --chr R018626 --chr R012949 --chr R000176 --chr R003076 \
			--chr R008561 --chr R012074 --chr R014489 --chr R003458 --chr R007312 --chr R001096 --chr R004261 --chr R008927 --chr R019137 --chr R012511 \
			--chr R008294 --chr R002720 --chr R019746 --chr R019817 --chr R018268 --chr R019872 --chr R009024 --chr R001944 --chr R009861 --chr R003690 \
			--chr R001848 --chr R003195 --chr R010585 --chr R008894 --chr R001065 --chr R008761 --chr R018275 --chr R002970 --chr R020723 --chr R012566 \
			--chr R015804 --chr R007627 --chr R013661 --chr R008450 --chr R004785 --chr R014827 --chr R013447 --chr R023088 --chr R009519 --chr R012062 \
			--chr R012833 --chr R000952 --chr R002874 --chr R020055 --chr R008729 --chr R004320 --chr R001364 --chr R012287 --chr R013278 --chr R008300 \
			--chr R011336 --chr R001397 --chr R009737 --chr R018007 --chr R004313 --chr R020328 --chr R008244 --chr R017974 --chr R007464 --chr R008256 \
			--chr R005675 --chr R006115 --chr R001451 --chr R016001 --chr R020854 --chr R001363 --chr R014752 --chr R018121 --chr R000421 --chr R009446 \
			--chr R017767 --chr R011726 --chr R004471 --chr R013531 --chr R011716 --chr R013578 --chr R002055 --chr R011535 --chr R005928 --chr R002422 \
			--chr R005536 --chr R007621 --chr R000552 --chr R013491 --chr R001366 --chr R002472 --chr R016758 --chr R008442 --chr R003323 --chr R008216 \
			--chr R012022 --chr R008288 --chr R006938 --chr R017413 --chr R011240 --chr R002948 --chr R017512 --chr R001409 --chr R007650 --chr R013059 \
			--chr R002068 --chr R005540 --chr R007054 --chr R007798 --chr R010385 --chr R003768 --chr R001929 --chr R008429 --chr R008731 --chr R018927 \
			--chr R005309 --chr R014581 --chr R004149 --chr R005334 --chr R003461 --chr R005262 --chr R007658 --chr R009086 --chr R004516 --chr R009538 \
			--chr R016838 --chr R013533 --chr R010286 --chr R016699 --chr R003545 --chr R018233 --chr R022968 --chr R002596 --chr R007445 --chr R006360 \
			--chr R022848 --chr R000319 --chr R003409 --chr R016818 --chr R003738 --chr R013256 --chr R005082 --chr R005594 --chr R002972 --chr R004641 \
			--chr R012375 --chr R011366 --chr R021554 --chr R015393 --chr R015149 --chr R009170 --chr R016935 --chr R013398 --chr R007751 --chr R008191 \
			--chr R007961 --chr R010036 --chr R001708 --chr R001861 --chr R011736 --chr R003732 --chr R021972 --chr R005154 --chr R011027 --chr R000469 \
			--chr R001412 --chr R004275 --chr R018759 --chr R007625 --chr R001153 --chr R011637 --chr R011458 --chr R006956 --chr R016069 --chr R015878 \
			--chr R002821 --chr R019614 --chr R022699 --chr R008632 --chr R001991 --chr R007955 --chr R013451 --chr R012736 --chr R012416 --chr R000243 \
			--chr R002724 --chr R013792 --chr R008306 --chr R008415 --chr R000229 --chr R004489 --chr R018254 --chr R006229 --chr R010504 --chr R000501 \
			--chr R008190 --chr R002785 --chr R005342 --chr R016418 --chr R001236 --chr R000716 --chr R011172 --chr R004560 --chr R003496 --chr R003856 \
			--chr R007963 --chr R008715 --chr R011895 --chr R001075 --chr R009654 --chr R002295 --chr R006241 --chr R015882 --chr R010013 --chr R008419 \
			--chr R021383 --chr R000629 --chr R013744 --chr R010059 --chr R010748 --chr R001395 --chr R010895 --chr R021258 --chr R005587 --chr R006496 \
			--chr R002030 --chr R001967 --chr R002641 --chr R010568 --chr R020553 --chr R018058 --chr R010157 --chr R009583 --chr R018631 --chr R008733 \
			--chr R002171 --chr R013092 --chr R003907 --chr R011358 --chr R023389 --chr R000067 --chr R007655 --chr R002404 --chr R008852 --chr R012077 \
			--chr R018782 --chr R004802 --chr R011580 --chr R012567 --chr R018018 --chr R005908 --chr R012640 --chr R019149 --chr R009893 --chr R012079 \
			--chr R001903 --chr R005898 --chr R015246 --chr R004553 --chr R009849 --chr R010446 --chr R016461 --chr R003667 --chr R001226 --chr R006669 \
			--chr R008425 --chr R000657 --chr R008408 --chr R000289 --chr R012039 --chr R003200 --chr R006484 --chr R020460 --chr R019899 --chr R019046 \
			--chr R008276 --chr R005330 --chr R013931 --chr R019091 --chr R008745 --chr R011631 --chr R006180 --chr R016637 --chr R002041 --chr R001362 \
			--chr R009967 --chr R009727 --chr R007918 --chr R020235 --chr R006186 --chr R012504 --chr R016272 --chr R019514 --chr R020444 --chr R017219 \
			--chr R014296 --chr R021310 --chr R000430 --chr R014318 --chr R007116 --chr R013056 --chr R011016 --chr R006525 --chr R003058 --chr R009476 \
			--chr R004657 --chr R008975 --chr R009373 --chr R017226 --chr R003736 --chr R012002 --chr R009816 --chr R004247 --chr R020798 --chr R010152 \
			--chr R004950 --chr R017498 --chr R016369 --chr R010148 --chr R005169 --chr R005352 --chr R008861 --chr R001003 --chr R018028 --chr R010918 \
			--chr R016335 --chr R000985 --chr R011368 --chr R000241 --chr R000255 --chr R004820 --chr R006265 --chr R019365 --chr R012453 --chr R010566 \
			--chr R008279 --chr R002406 --chr R018640 --chr R001437 --chr R014510 --chr R009810 --chr R006545 --chr R013761 --chr R003282 --chr R013168 \
			--chr R010764 --chr R013283 --chr R006278 --chr R005392 --chr R000082 --chr R013628 --chr R006765 --chr R014185 --chr R011837 --chr R020964 \
			--chr R023373 --chr R016453 --chr R007240 --chr R013798 --chr R015837 --chr R013658 --chr R019646 --chr R017466 --chr R007712 --chr R008743 \
			--chr R010611 --chr R001930 --chr R011523 --chr R005028 --chr R006260 --chr R006626 --chr R005445 --chr R000992 --chr R009025 --chr R012244 \
			--chr R016346 --chr R000079 --chr R015658 --chr R015556 --chr R014301 --chr R008383 --chr R010461 --chr R002272 --chr R009178 --chr R000924 \
			--chr R005062 --chr R017961 --chr R008718 --chr R004707 --chr R005160 --chr R019139 --chr R020265 --chr R001527 --chr R009394 --chr R006745 \
			--chr R009253 --chr R012709 --chr R017986 --chr R003357 --chr R006931 --chr R010421 --chr R023825 --chr R000967 --chr R023041 --chr R006192 \
			--chr R003601 --chr R013448 --chr R008116 --chr R000916 --chr R000609 --chr R010274 --chr R010326 --chr R000821 --chr R012089 --chr R004365 --chr R001272 --chr R018153 --chr R011352 --chr R020302 --chr R009604 --chr R012515 --chr R000614 --chr R018270 --chr R014238 --chr R002313 --chr R011441 --chr R008208 --chr R003100 --chr R004367 --chr R015033 --chr R005620 --chr R000756 --chr R001762 --chr R008886 --chr R002925 --chr R017703 --chr R018086 --chr R016540 --chr R017213 --chr R011813 --chr R006531 --chr R013873 --chr R006023 --chr R017833 --chr R018850 --chr R011086 --chr R023573 --chr R003318 --chr R008813 --chr R002545 --chr R007411 --chr R004216 --chr R008627 --chr R021835 --chr R007486 --chr R000147 --chr R000663 --chr R004158 --chr R004521 --chr R023532 --chr R003588 --chr R003138 --chr R008645 --chr R012274 --chr R003656 --chr R002440 --chr R012912 --chr R020882 --chr R000604 --chr R003024 --chr R008142 --chr R011746 --chr R003089 --chr R006376 --chr R005823 --chr R000939 --chr R022947 --chr R023072 --chr R007704 --chr R006130 --chr R001044 --chr R007940 --chr R009073 --chr R003731 --chr R005676 --chr R021177 --chr R014895 --chr R009628 --chr R003806 --chr R006743 --chr R009126 --chr R010756 --chr R002465 --chr R011087 --chr R000276 --chr R015357 --chr R022516 --chr R003666 --chr R014968 --chr R008464 --chr R006227 --chr R003781 --chr R014930 --chr R003644 --chr R004523 --chr R016158 --chr R015377 --chr R004182 --chr R003491 --chr R009947 --chr R001220 --chr R000815 --chr R003054 --chr R000934 --chr R002279 --chr R011739 --chr R012673 --chr R000281 --chr R001776 --chr R013584 --chr R000908 --chr R011791 --chr R007539 --chr R016697 --chr R008751 --chr R010066 --chr R007548 --chr R011668 --chr R007646 --chr R002527 --chr R001176 --chr R015070 --chr R002905 --chr R016202 --chr R007198 --chr R020327 --chr R003385 --chr R005771 --chr R019507 --chr R001965 --chr R007594 --chr R017042 --chr R008520 --chr R011606 --chr R016174 --chr R010744 --chr R000577 --chr R018384 --chr R009286 --chr R004025 --chr R020498 --chr R008741 --chr R002353 --chr R000883 --chr R002393 --chr R019719 --chr R006925 --chr R016903 --chr R013920 --chr R020652 --chr R001863 --chr R000978 --chr R012180 --chr R006391 --chr R019436 --chr R018467 --chr R008460 --chr R006775 --chr R011848 --chr R000818 --chr R013465 --chr R007832 --chr R002525 --chr R001571 --chr R010411 --chr R023377 --chr R002446 --chr R022561 --chr R008760 --chr R010293 --chr R005721 --chr R007176 --chr R012892 --chr R011932 --chr R011133 --chr R013634 --chr R013545 --chr R006294 --chr R015338 --chr R002613 --chr R008052 --chr R006550 --chr R009021 --chr R002815 --chr R007713 --chr R002707 --chr R002108 --chr R002759 --chr R005389 --chr R017730 --chr R004416 --chr R012983 --chr R017718 --chr R004738 --chr R019579 --chr R016129 --chr R022752 --chr R000071 --chr R001295 --chr R006432 --chr R012434 --chr R003275 --chr R004430 --chr R006543 --chr R002558 --chr R001560 --chr R001303 --chr R013975 --chr R020920 --chr R005450 --chr R005975 --chr R006083 --chr R016595 --chr R003117 --chr R008695 --chr R002415 --chr R019067 --chr R016041 --chr R020274 --chr R000297 --chr R019808 --chr R000087 --chr R023775 --chr R004956 --chr R012857 --chr R010175 --chr R007534 --chr R021868 --chr R004944 --chr R008563 --chr R000051 --chr R020343 --chr R008221 --chr R009294 --chr R000721 --chr R004710 --chr R004001 --chr R004787 --chr R008128 --chr R013259 --chr R021165 --chr R013405 --chr R015979 --chr R009554 --chr R012934 --chr R000057 --chr R002862 --chr R014660 --chr R016125 --chr R019280 --chr R001436 --chr R005754 --chr R012884 --chr R015083 --chr R007121 --chr R010074 --chr R002015 --chr R010580 --chr R003799 --chr R018574 --chr R001621 --chr R006540 --chr R010853 --chr R002129 --chr R004932 --chr R001919 --chr R017598 --chr R008593 --chr R019642 --chr R015952 --chr R003778 --chr R018415 --chr R002877 --chr R023104 --chr R018483 --chr R019270 --chr R011114 --chr R000350 --chr R018945 --chr R022247 --chr R022856 --chr R004309 --chr R003453 --chr R002283 --chr R003264 --chr R006689 --chr R004991 --chr R012558 --chr R007195 --chr R010137 --chr R014567 --chr R018363 --chr R015565 --chr R011429 --chr R017214 --chr R016560 --chr R021682 --chr R004616 --chr R013233 --chr R003153 --chr R001627 --chr R020573 --chr R015013 --chr R007931 --chr R020192 --chr R004417 --chr R021053 --chr R004678 --chr R006445 --chr R016647 --chr R010614 --chr R010127 --chr R012539 --chr R007933 --chr R020970 --chr R021499 --chr R015839 --chr R010665 --chr R005194 --chr R012487 --chr R003717 --chr R002548 --chr R010493 --chr R011394 --chr R017433 --chr R019381 --chr R002448 --chr R008711 --chr R004298 --chr R002106 --chr R011908 --chr R006271 --chr R016588 --chr R016854 --chr R004192 --chr R018283 --chr R007748 --chr R013587 --chr R007879 --chr R011670 --chr R016187 --chr R007935 --chr R015986 --chr R002870 --chr R013345 --chr R007096 --chr R011860 --chr R011142 --chr R022218 --chr R013906 --chr R018696 --chr R016815 --chr R002246 --chr R002072 --chr R012547 --chr R000035 --chr R017588 --chr R008932 --chr R009130 --chr R001163 --chr R015160 --chr R020631 --chr R019501 --chr R005167 --chr R015691 --chr R008009 --chr R022024 --chr R011270 --chr R002928 --chr R001346 --chr R007499 --chr R000381 --chr R010405 --chr R002530 --chr R006383 --chr R009398 --chr R005615 --chr R004151 --chr R007568 --chr R002321 --chr R009541 --chr R021777 --chr R000259 --chr R008312 --chr R024086 --chr R000749 --chr R018939 --chr R007153 --chr R006868 --chr R014330 --chr R007082 --chr R011071 --chr R015792 --chr R000820 --chr R010912 --chr R002155 --chr R016017 --chr R007657 --chr R015778 --chr R003493 --chr R001645 --chr R021156 --chr R013987 --chr R015488 --chr R003422 --chr R001933 --chr R001033 --chr R015178 --chr R005788 --chr R016176 --chr R010210 --chr R013579 --chr R001988 --chr R002643 --chr R010238 --chr R004501 --chr R007509 --chr R002070 --chr R003104 --chr R009515 --chr R002508 --chr R013470 --chr R005603 --chr R007996 --chr R003533 --chr R012579 --chr R000129 --chr R004704 --chr R016905 --chr R000161 --chr R008241 --chr R018976 --chr R009198 --chr R022505 --chr R008273 --chr R014142 --chr R008581 --chr R015294 --chr R009978 --chr R002782 --chr R012000 --chr R008713 --chr R005293 --chr R011277 --chr R006087 --chr R013645 --chr R016769 --chr R007046 --chr R023518 --chr R010094 --chr R002547 --chr R021051 --chr R008517 --chr R003142 --chr R015379 --chr R004246 --chr R004718 --chr R008810 --chr R003281 --chr R020637 --chr R001251 --chr R020960 --chr R011683 --chr R005644 --chr R005811 --chr R002166 --chr R019694 --chr R014544 --chr R005832 --chr R000393 --chr R018291 --chr R014309 --chr R020132 --chr R012956 --chr R006611 --chr R022627 --chr R004846 --chr R022271 --chr R009868 --chr R007037 --chr R016140 --chr R016052 --chr R018745 --chr R007293 --chr R013540 --chr R004605 --chr R004094 --chr R013660 --chr R000099 --chr R011750 --chr R009467 --chr R001357 --chr R015870 --chr R016250 --chr R009132 --chr R007550 --chr R021572 --chr R010790 --chr R015942 --chr R003140 --chr R009880 --chr R012344 --chr R005382 --chr R005001 --chr R007407 --chr R013636 --chr R001428 --chr R013563 --chr R002746 --chr R006112 --chr R023094 --chr R001212 --chr R012931 --chr R014768 --chr R016192 --chr R002302 --chr R011436 --chr R008762 --chr R013755 --chr R001274 --chr R003703 --chr R003157 --chr R019322 --chr R010069 --chr R011851 --chr R010582 --chr R002326 --chr R000429 --chr R011407 --chr R010065 --chr R000315 --chr R001143 --chr R005793 --chr R019389 --chr R008511 --chr R020386 --chr R016499 --chr R006281 --chr R018643 --chr R001396 --chr R004632 --chr R014683 --chr R006043 --chr R019551 --chr R010875 --chr R007139 --chr R017285 --chr R012382 --chr R007974 --chr R023676 --chr R019631 --chr R008405 --chr R004967 --chr R000450 --chr R002275 --chr R009399 --chr R001670 --chr R017511 --chr R023978 --chr R021974 --chr R002962 --chr R015015 --chr R002028 --chr R001612 --chr R008976 --chr R001433 --chr R005436 --chr R022967 --chr R006577 --chr R013131 --chr R018003 --chr R023033 --chr R015604 --chr R006616 --chr R020837 --chr R020364 --chr R002260 --chr R014919 --chr R020287 --chr R006653 --chr R008699 --chr R006501 --chr R002463 --chr R009726 --chr R006524 --chr R012991 --chr R014722 --chr R005101 --chr R005374 --chr R018016 --chr R006237 --chr R011846 --chr R019386 --chr R001322 --chr R010640 --chr R008709 --chr R016262 --chr R013801 --chr R013417 --chr R002107 --chr R005551 --chr R021186 --chr R004233 --chr R007677 --chr R011980 --chr R008640 --chr R021309 --chr R013675 --chr R003594 --chr R001629 --chr R011930 --chr R008136 --chr R000671 --chr R009261 --chr R015446 --chr R013442 --chr R012852 --chr R004386 --chr R008430 --chr R001036 --chr R004484 --chr R015823 --chr R001686 --chr R008636 --chr R008831 --chr R004613 --chr R017879 --chr R018256 --chr R012814 --chr R006621 --chr R007680 --chr R008282 --chr R005515 --chr R000896 --chr R010115 --chr R012103 --chr R006878 --chr R012738 --chr R000347 --chr R003976 --chr R017553 --chr R009030 --chr R002119 --chr R022681 --chr R023442 --chr R020578 --chr R009417 --chr R002080 --chr R005339 --chr R018355 --chr R001821 --chr R013097 --chr R004543 --chr R011744 --chr R000196 --chr R004150 --chr R009518 --chr R016165 --chr R007229 --chr R010423 --chr R001666 --chr R004809 --chr R022117 --chr R014343 --chr R015082 --chr R000091 --chr R010948 --chr R002752 --chr R014912 --chr R008980 --chr R004481 --chr R001820 --chr R014689 --chr R010543 --chr R012815 --chr R000567 --chr R013830 --chr R003830 --chr R006742 --chr R000231 --chr R002945 --chr R011091 --chr R003112 --chr R003720 --chr R015163 --chr R019727 --chr R006156 --chr R003878 --chr R022577 --chr R001126 --chr R009695 --chr R012630 --chr R000493 --chr R020827 --chr R022498 --chr R009833 --chr R010788 --chr R003606 --chr R007286 --chr R006057 --chr R014922 --chr R023752 --chr R006569 --chr R007645 --chr R000767 --chr R000733 --chr R013355 --chr R012679 --chr R010234 --chr R000428 --chr R005504 --chr R015417 --chr R000171 --chr R022298 --chr R006677 --chr R002730 --chr R000317 --chr R021823 --chr R013112 --chr R004732 --chr R019010 --chr R007249 --chr R011214 --chr R007520 --chr R012087 --chr R001425 --chr R000342 --chr R017716 --chr R009535 --chr R012242 --chr R010548 --chr R021576 --chr R013307 --chr R005422 --chr R022142 --chr R013843 --chr R001714 --chr R020258 --chr R022966 --chr R000107 --chr R004286 --chr R013959 --chr R023741 --chr R007522 --chr R014182 --chr R011329 --chr R005555 --chr R005589 --chr R005677 --chr R007731 --chr R014990 --chr R019177 --chr R007146 --chr R013566 --chr R022260 --chr R003700 --chr R006413 --chr R010905 --chr R009479 --chr R012447 --chr R004425 --chr R017337 --chr R000073 --chr R008164 --chr R002754 --chr R001875 --chr R014716 --chr R022125 --chr R008781 --chr R011981 --chr R018025 --chr R010708 --chr R007792 --chr R017960 --chr R021049 --chr R000842 --chr R011567 --chr R012711 --chr R019510 --chr R004242 --chr R009905 --chr R003740 --chr R001219 --chr R005253 --chr R017513 --chr R018356 --chr R017483 --chr R003676 --chr R016767 --chr R005355 --chr R019185 --chr R012295 --chr R010003 --chr R008239 --chr R013649 --chr R014617 --chr R013730 --chr R009599 --chr R005090 --chr R015665 --chr R016616 --chr R018322 --chr R013997 --chr R000415 --chr R019924 --chr R015655 --chr R012011 --chr R020562 --chr R010725 --chr R003631 --chr R014730 --chr R018924 --chr R014000 --chr R000250 --chr R016970 --chr R002088 --chr R010852 --chr R010257 --chr R024067 --chr R017377 --chr R016795 --chr R002350 --chr R008473 --chr R006950 --chr R017172 --chr R003899 --chr R003664 --chr R017021 --chr R003967 --chr R003462 --chr R018984 --chr R013518 --chr R024123 --chr R020203 --chr R008665 --chr R003329 --chr R020112 --chr R004931 --chr R006896 --chr R010243 --chr R016859 --chr R009811 --chr R011576 --chr R010864 --chr R019097 --chr R012840 --chr R000135 --chr R017991 --chr R023973 --chr R017109 --chr R014446 --chr R009809 --chr R014027 --chr R018848 --chr R020127 --chr R000476 --chr R016016 --chr R004679 --chr R020006 --chr R014548 --chr R020373 --chr R006916 --chr R013298 --chr R001510 --chr R020526 --chr R015028 --chr R002230 --chr R014561 --chr R018619 --chr R014023 --chr R004953 --chr R009718 --chr R009614 --chr R023311 --chr R009468 --chr R022131 --chr R008249 --chr R022684 --chr R007438 --chr R013594 --chr R013577 --chr R005181 --chr R005034 --chr R003347 --chr R007492 --chr R017288 --chr R022809 --chr R001179 --chr R006561 --chr R003541 --chr R021494 --chr R006671 --chr R019240 --chr R020684 --chr R007667 --chr R013582 --chr R018014 --chr R014342 --chr R016812 --chr R003872 --chr R003305 --chr R000630 --chr R005713 --chr R007837 --chr R020473 --chr R021674 --chr R017572 --chr R009659 --chr R018410 --chr R000025 --chr R006355 --chr R002812 --chr R017452 --chr R013941 --chr R009332 --chr R006157 --chr R000076 --chr R019885 --chr R002810 --chr R005897 --chr R010379 --chr R019573 --chr R020196 --chr R013021 --chr R006537 --chr R001652 --chr R004949 --chr R010242 --chr R018935 --chr R006350 --chr R011695 --chr R005919 --chr R010206 --chr R006374 --chr R008286 --chr R004397 --chr R015654 --chr R019088 --chr R003481 --chr R011463 --chr R009184 --chr R001382 --chr R003084 --chr R003750 --chr R010168 --chr R000819 --chr R008680 --chr R012312 --chr R020242 --chr R006771 --chr R015981 --chr R010967 --chr R004798 --chr R001334 --chr R014058 --chr R002867 --chr R000958 --chr R009251 --chr R001556 --chr R002543 --chr R007571 --chr R000490 --chr R005050 --chr R013567 --chr R023598 --chr R018234 --chr R018285 --chr R019328 --chr R010914 --chr R016529 --chr R023734 --chr R016357 --chr R006394 --chr R008238 --chr R001136 --chr R005182 --chr R003972 --chr R012492 --chr R015663 --chr R007049 --chr R014218 --chr R014709 --chr R007952 --chr R002686 --chr R006349 --chr R009738 --chr R001040 --chr R014361 --chr R016037 --chr R007182 --chr R000579 --chr R020935 --chr R000670 --chr R006865 --chr R007089 --chr R021233 --chr R010495 --chr R001689 --chr R016153 --chr R004848 --chr R003819 --chr R000581 --chr R013497 --chr R010944 --chr R001223 --chr R019754 --chr R003760 --chr R003777 --chr R004948 --chr R007214 --chr R021829 --chr R012817 --chr R000291 --chr R015541 --chr R011826 --chr R006623 --chr R017384 --chr R002100 --chr R010641 --chr R003032 --chr R006321 --chr R004363 --chr R001408 --chr R005520 --chr R019825 --chr R003210 --chr R004832 --chr R022441 --chr R006897 --chr R018297 --chr R020710 --chr R018087 --chr R007523 --chr R006084 --chr R000074 --chr R014198 --chr R011308 --chr R000907 --chr R000127 --chr R021690 --chr R018519 --chr R017074 --chr R022911 --chr R010023 --chr R000857 --chr R009793 --chr R009741 --chr R009128 --chr R020390 --chr R009867 --chr R018513 --chr R017838 --chr R010441 --chr R002531 --chr R008361 --chr R001865 --chr R021312 --chr R006650 --chr R009759 --chr R003683 --chr R001529 --chr R007648 --chr R002163 --chr R001452 --chr R008291 --chr R014995 --chr R017300 --chr R018838 --chr R022754 --chr R004945 --chr R000104 --chr R005561 --chr R014520 --chr R005579 --chr R005921 --chr R013775 --chr R013338 --chr R012109 --chr R005127 --chr R003301 --chr R007717 --chr R009648 --chr R017439 --chr R010516 --chr R008365 --chr R021707 --chr R019440 --chr R013556 --chr R014398 --chr R021962 --chr R005549 --chr R010196 --chr R019878 --chr R004500 --chr R011507 --chr R008315 --chr R017635 --chr R011909 --chr R004294 --chr R011858 --chr R020310 --chr R019326 --chr R024065 --chr R011383 --chr R010797 --chr R001444 --chr R009495 --chr R015966 --chr R023942 --chr R009549 --chr R003589 --chr R007804 --chr R001279 --chr R002609 --chr R015212 --chr R000795 --chr R004159 --chr R006664 --chr R006270 --chr R013514 --chr R008222 --chr R010794 --chr R003646 --chr R023422 --chr R004683 --chr R017786 --chr R010969 --chr R014168 --chr R006565 --chr R023385 --chr R002520 --chr R009756 --chr R014113 --chr R016814 --chr R007300 --chr R002510 --chr R002946 --chr R000707 --chr R008093 --chr R008019 --chr R017407 --chr R012722 --chr R009083 --chr R001478 --chr R004465 --chr R019277 --chr R016933 --chr R001491 --chr R012356 --chr R006080 --chr R003277 --chr R004960 --chr R018716 --chr R002212 --chr R008586 --chr R003177 --chr R004525 --chr R008660 --chr R008501 --chr R002218 --chr R010957 --chr R004767 --chr R000903 --chr R010829 --chr R006167 --chr R000302 --chr R004268 --chr R006707 --chr R024104 --chr R021816 --chr R001268 --chr R002618 --chr R000460 --chr R009739 --chr R002949 --chr R008662 --chr R020368 --chr R020691 --chr R001462 --chr R005371 --chr R017885 --chr R019484 --chr R001214 --chr R002312 --chr R016171 --chr R001161 --chr R005782 --chr R006965 --chr R015621 --chr R020966 --chr R001570 --chr R001677 --chr R002299 --chr R007528 --chr R010017 --chr R015868 --chr R000642 --chr R003964 --chr R007800 --chr R007066 --chr R018182 --chr R001549 --chr R013654 --chr R014108 --chr R005002 --chr R002243 --chr R000167 --chr R002509 --chr R007998 --chr R014669 --chr R010916 --chr R010236 --chr R000111 --chr R010435 --chr R001805 --chr R005636 --chr R005106 --chr R004614 --chr R002888 --chr R002424 --chr R011103 --chr R002556 --chr R006679 --chr R010009 --chr R021866 --chr R016940 --chr R013406 --chr R005743 --chr R002460 --chr R006967 --chr R008168 --chr R002959 --chr R007813 --chr R001221 --chr R013226 --chr R002931 --chr R005386 --chr R006819 --chr R001313 --chr R015166 --chr R014239 --chr R012847 --chr R007628 --chr R021687 --chr R020763 --chr R017555 --chr R009636 --chr R000639 --chr R017234 --chr R023870 --chr R013243 --chr R004389 --chr R016179 --chr R018062 --chr R014851 --chr R008454 --chr R007271 --chr R001285 --chr R020141 --chr R011843 --chr R011699 --chr R013334 --chr R021196 --chr R019630 --chr R004016 --chr R022224 --chr R004930 --chr R005033 --chr R006947 --chr R000218 --chr R009270 --chr R003889 --chr R010572 --chr R011487 --chr R009091 --chr R000272 --chr R016113 --chr R002365 --chr R007340 --chr R000752 --chr R006411 --chr R004064 --chr R023465 --chr R009715 --chr R011431 --chr R001613 --chr R008252 --chr R000437 --chr R004862 --chr R001494 --chr R010791 --chr R004270 --chr R019083 --chr R012069 --chr R006221 --chr R010471 --chr R001447 --chr R004385 --chr R013586 --chr R007136 --chr R016554 --chr R017409 --chr R016713 --chr R018993 --chr R007985 --chr R018736 --chr R002227 --chr R005419 --chr R014686 --chr R006732 --chr R010761 --chr R003233 --chr R011117 --chr R012661 --chr R005005 --chr R002506 --chr R013690 --chr R004920 --chr R006243 --chr R002010 --chr R002110 --chr R000371 --chr R017033 --chr R021163 --chr R001623 --chr R001092 --chr R011698 --chr R006244 --chr R008969 --chr R000676 --chr R001808 --chr R009032 --chr R008179 --chr R006208 --chr R012592 --chr R003751 --chr R004061 --chr R004053 --chr R007886 --chr R008903 --chr R006515 --chr R003807 --chr R001971 --chr R018142 --chr R007090 --chr R010453 --chr R006322 --chr R004896 --chr R021116 --chr R004468 --chr R011502 --chr R012298 --chr R020709 --chr R011007 --chr R011459 --chr R000300 --chr R021022 --chr R001691 --chr R015496 --chr R021857 --chr R017165 --chr R012893 --chr R014595 --chr R022407 --chr R009842 --chr R000400 --chr R011620 --chr R013149 --chr R011108 --chr R015997 --chr R014867 --chr R004402 --chr R001793 --chr R008962 --chr R016572 --chr R007291 --chr R007758 --chr R012755 --chr R007858 --chr R008876 --chr R018060 --chr R005055 --chr R010375 --chr R018560 --chr R011179 --chr R006674 --chr R001614 --chr R010226 --chr R017662 --chr R010237 --chr R008375 --chr R009267 --chr R012881 --chr R016315 --chr R004962 --chr R006858 --chr R001424 --chr R009956 --chr R003622 --chr R005806 --chr R013606 --chr R017638 --chr R002153 --chr R005446 --chr R007243 --chr R008346 --chr R002616 --chr R014834 --chr R002304 --chr R010770 --chr R006516 --chr R013985 --chr R008071 --chr R023051 --chr R008995 --chr R000027 --chr R002458 --chr R001993 --chr R012578 --chr R001688 --chr R014634 --chr R019932 --chr R018257 --chr R015299 --chr R009937 --chr R006293 --chr R010125 --chr R018097 --chr R015787 --chr R023840 --chr R021503 --chr R003876 --chr R020159 --chr R012004 --chr R011118 --chr R016413 --chr R001530 --chr R019262 --chr R006081 --chr R000989 --chr R006955 --chr R014714 --chr R003743 --chr R024098 --chr R005490 --chr R014906 --chr R003136 --chr R009050 --chr R010688 --chr R012590 --chr R002675 --chr R005155 --chr R001438 --chr R013295 --chr R012652 --chr R006003 --chr R015884 --chr R009663 --chr R003983 --chr R001427 --chr R007349 --chr R017244 --chr R001832 --chr R005718 --chr R001021 --chr R013808 --chr R016898 --chr R006891 --chr R008081 --chr R012411 --chr R015916 --chr R006704 --chr R007193 --chr R001679 --chr R012449 --chr R018463 --chr R014786 --chr R015957 --chr R008399 --chr R018666 --chr R008566 --chr R014071 --chr R011925 --chr R018996 --chr R015362 --chr R021676 --chr R012404 --chr R013984 --chr R017385 --chr R024009 --chr R000549 --chr R011450 --chr R003634 --chr R018238 --chr R005687 --chr R000109 --chr R000760 --chr R007001 --chr R010410 --chr R012389 --chr R017332 --chr R002073 --chr R022791 --chr R006332 --chr R003452 --chr R016706 --chr R002828 --chr R002958 --chr R000927 --chr R004665 --chr R003797 --chr R012901 --chr R008735 --chr R003944 --chr R003566 --chr R017597 --chr R016582 --chr R005272 --chr R015481 --chr R018544 --chr R006867 --chr R019928 --chr R012293 --chr R006603 --chr R016692 --chr R005135 --chr R004431 --chr R006979 --chr R009774 --chr R007526 --chr R000515 --chr R015602 --chr R006739 --chr R006000 --chr R021146 --chr R001565 --chr R001543 --chr R011489 --chr R006155 --chr R014593 --chr R001296 --chr R010845 --chr R001479 --chr R010479 --chr R013431 --chr R005834 --chr R005178 --chr R010773 --chr R004621 --chr R006935 --chr R019265 --chr R006538 --chr R021354 --chr R018106 --chr R017663 --chr R011112 --chr R009316 --chr R016035 --chr R005773 --chr R022740 --chr R008811 --chr R006930 --chr R008977 --chr R006184 --chr R008125 --chr R004591 --chr R008247 --chr R024079 --chr R017168 --chr R009432 --chr R017148 --chr R020129 --chr R022382 --chr R005370 --chr R013943 --chr R004806 --chr R006870 --chr R012812 --chr R010200 --chr R024127 --chr R008630 --chr R002904 --chr R010956 --chr R008170 --chr R003364 --chr R010723 --chr R013138 --chr R003206 --chr R007282 --chr R016088 --chr R013042 --chr R016057 --chr R014792 --chr R017030 --chr R019781 --chr R012439 --chr R019532 --chr R008202 --chr R013327 --chr R003887 --chr R001752 --chr R012254 --chr R016106 --chr R004006 --chr R005459 --chr R009036 --chr R005694 --chr R017451 --chr R010347 --chr R006161 --chr R009837 --chr R018569 --chr R006368 --chr R022315 --chr R003490 --chr R017432 --chr R009264 --chr R004410 --chr R016154 --chr R013289 --chr R015020 --chr R016439 --chr R008915 --chr R004072 --chr R010445 --chr R003612 --chr R006760 --chr R007715 --chr R006933 --chr R003051 --chr R016571 --chr R006544 --chr R004401 --chr R009392 --chr R007545 --chr R010308 --chr R012807 --chr R020341 --chr R016009 --chr R005758 --chr R000120 --chr R004295 --chr R009033 --chr R019535 --chr R007518 --chr R003146 --chr R020220 --chr R013135 --chr R005716 --chr R014577 --chr R010373 --chr R013323 --chr R015428 --chr R002358 --chr R015892
			
		###Align the sequences:
		
		$ bwa mem -M GCF_018296145.1_Otsh_v2.0_genomic.fna Meeks_et_al.FlankingSeqs.fasta | samtools sort -o GCF_018296145.1.vs.Meeks2016.bam
			
		$ samtools index GCF_018296145.1.vs.Meeks2016.bam
		
		####BWA mem doesn't handle sequences smaller than 70 bp, so I had to modify the script
		
		$ python Bam2SNPPosition.v1.1_SpecificProject_OtshTruth.py -file GCF_018296145.1.vs.Meeks2016.bam -qual 40 -multiple 1 -minLen 79 -minPid 90 -pos 11 > GCF_018296145.1.vs.Meeks2016.positions.txt
		
		#### QC
		
		$ python CheckMappingVCF.v1.1.py \
		-map GCF_018296145.1.vs.Meeks2016.positions.txt \
		-fasta1 Meeks_et_al.FlankingSeqs.fasta \
		-fai1 Meeks_et_al.FlankingSeqs.fasta.fai \
		-fasta2 /scratch2/shared/NCBI.Genomes/GCF_018296145.1_Otsh_v2.0_genomic.fna \
		-fai2 /scratch2/shared/NCBI.Genomes/GCF_018296145.1_Otsh_v2.0_genomic.fna.fai \
		-flanking 5 \
		-output most > \
		GCF_018296145.1.vs.Meeks2016.positions.QCfiltered.txt
		
		Finished reading vcf file: Found 19356 variants, correct variants:5939, unknown variants:3634, CompleteMismatch:7812, variantOnlyMismatch:1879, partialMatch:92
		
		###Map reads to new vcf file
		###Had to manually remove part of the name in the Brieuc2Otsh.NCBIv2.positions.custom.QCfiltered.txt file that was added by the VCF_2_Fasta script
		Meeks/doi_10.5061_dryad.j29f5__v1$ python3 \
		~/Desktop/Documents/General_Scripts/mapSNP2NewGenome/Assign_New_SNP_Positions.V1.1.py \
		-map GCF_018296145.1.vs.Meeks2016.positions.QCfiltered.txt \
		-vcf Meeks_et_all.GeneticMapOnly.recode.vcf \
		-fai ../../Brieuc/GCF_018296145.1_Otsh_v2.0_genomic.fna.fai > \
		Meeks_et_all.GeneticMapOnly_GCF_018296145.vcf

		
		Meeks/doi_10.5061_dryad.j29f5__v1$ \
		bcftools sort Meeks_et_all.GeneticMapOnly_GCF_018296145.vcf > Meeks_et_all.GeneticMapOnly_GCF_018296145.sorted.vcf
	
		Meeks/doi_10.5061_dryad.j29f5__v1$ \
		bgzip Meeks_et_all.GeneticMapOnly_GCF_018296145.sorted.vcf
		Meeks/doi_10.5061_dryad.j29f5__v1$ \
		tabix Meeks_et_all.GeneticMapOnly_GCF_018296145.sorted.vcf.gz
		
		###821 SNPs left
		
		
	###Manually combined both vcf files (and removed duplications - 3760 variants left)
		
	CombinedTruth_22Nov2022_GCF_018296145.sorted.vcf 
	
	
	###Adding variants that have been filtered from called SNPs:
	
	$ vcftools \
	--gzvcf ../vcf/variants.GATK.iteration.1.Full.sorted.vcf.gz \
	--out TrainingSetHardFilterMAF0.3_Depth10-60_HWE0.3_Missing0.1_BiSNPs \
	--remove-indels \
	--remove-filtered-all \
	--recode \
	--recode-INFO-all \
	--maf 0.3 \
	--max-maf 0.7 \
	--min-alleles 2 \
	--max-alleles 2 \
	--min-meanDP 10 \
	--max-meanDP 60 \
	--hwe 0.3 \
	--max-missing 0.9

		VCFtools - 0.1.15
		(C) Adam Auton and Anthony Marcketta 2009
		Parameters as interpreted:
			--gzvcf ../vcf/variants.GATK.iteration.1.Full.sorted.vcf.gz
			--recode-INFO-all
			--maf 0.3
			--max-alleles 2
			--max-maf 0.7
			--max-meanDP 60
			--min-alleles 2
			--hwe 0.3
			--min-meanDP 10
			--max-missing 0.9
			--out TrainingSetHardFilterMAF0.3_Depth10-60_HWE0.3_Missing0.1_BiSNPs
			--recode
			--remove-filtered-all
			--remove-indels

	After filtering, kept 574586 out of a possible 69628954 Sites

	$ vcftools --vcf TrainingSetHardFilterMAF0.3_Depth10-60_HWE0.3_Missing0.1_BiSNPs.recode.vcf --out TrainingSetHardFilterMAF0.3_Depth10-60_HWE0.3_Missing0.1_BiSNPs_Thin1Mbp_Chr --recode --recode-INFO-all --thin 500000 --chr NC_056429.1 --chr NC_056430.1 --chr NC_056431.1 --chr NC_056432.1 --chr NC_056433.1 --chr NC_056434.1 --chr NC_056435.1 --chr NC_056436.1 --chr NC_056437.1 --chr NC_056438.1 --chr NC_056439.1 --chr NC_056440.1 --chr NC_056441.1 --chr NC_056442.1 --chr NC_056443.1 --chr NC_056444.1 --chr NC_056445.1 --chr NC_056446.1 --chr NC_056447.1 --chr NC_056448.1 --chr NC_056449.1 --chr NC_056450.1 --chr NC_056451.1 --chr NC_056452.1 --chr NC_056453.1 --chr NC_056454.1 --chr NC_056455.1 --chr NC_056456.1 --chr NC_056457.1 --chr NC_056458.1 --chr NC_056459.1 --chr NC_056460.1 --chr NC_056461.1 --chr NC_056462.1
		
	After filtering, kept 3840 out of a possible 574586 Sites
		
#############################################
### Recalibration      ######################
#############################################

	###Find the intersection ###
	
	$ vcf-isec \
	-n =2 \
	-f vcf/variants.GATK.iteration.1.Full.sorted.vcf.gz \
	truth/CombinedTruth_22Nov2022_GCF_018296145.sorted.vcf.gz \
	| bgzip -c > CombinedTruth_22Nov2022_GCF_018296145.sorted.intersection.vcf.gz
	
	After filtering, kept 3478 out of a possible 3478 Sites
	
	
	###Actual variant recalibration ### (errored out without enough truth markers -- trying to add a few thousand more -- see truth section for filtering criteria -- that worked need ~7000-8000 markers)
	
	$ java -Xms5g -Xmx176g \
	-jar GenomeAnalysisTK.jar \
	-T VariantRecalibrator \
	-R genome/GCF_018296145.1_Otsh_v2.0_genomic.fna \
	-input vcf/variants.GATK.iteration.1.Full.sorted.vcf.gz \
	-recalFile recal/variants.GATK.iteration.1.Full.recal \
	-tranchesFile tranches/variants.GATK.iteration.1.Full.tranches \
	-rscriptFile rFiles/variants.GATK.iteration.1.Full.R \
	-resource:GeneticMaps,known=false,training=true,truth=true,prior=15.0 truth/CombinedTruth_22Nov2022_GCF_018296145.sorted.intersection.vcf.gz \
	-resource:HardFilt,known=false,training=true,truth=false,prior=10.0 truth2/TrainingSetHardFilterMAF0.3_Depth10-60_HWE0.3_Missing0.1_BiSNPs_Thin1Mbp_Chr.recode.vcf.gz \
	-an QD -an MQ -an MQRankSum -an ReadPosRankSum -an FS -an SOR -an DP -an InbreedingCoeff -mode SNP
	
	###########################################################################
	### Apply recalibration ###################################################
	###########################################################################
	
	$ java -Xms5g -Xmx176g \
	-jar GenomeAnalysisTK.jar \
	-T ApplyRecalibration \
	-R genome/GCF_018296145.1_Otsh_v2.0_genomic.fna \
	-input vcf/variants.GATK.iteration.1.Full.sorted.vcf.gz \
	-recalFile recal/variants.GATK.iteration.1.Full.recal \
	-tranchesFile tranches/variants.GATK.iteration.1.Full.tranches \
	-mode SNP \
	-o vcf/variants.GATK.iteration.1.Full.sorted.recalibrated.vcf \
	--ts_filter_level 99.5
	
	
