## Supplementary material for "Revealing the evolutionary history and contemporary population structure of Pacific salmon in the Fraser River through genome resequencing": File S2: readme.txt

###Generate Truth dataset from a genetic map generated using a 200K SNP chip data (unpublished)

###First make a vcf file from the mareyMap
$ grep "NC_" Order.Okis.Michelle.Filt.LOD.15.size.90.mask.1.LOD.10.LODD.4.SexAveraged.1.mMap | awk '{print $5,$6,$7}' > Order.Okis.Michelle.Filt.LOD.15.size.90.mask.1.LOD.10.LODD.4.SexAveraged.1.vcf

################################
###Map reads to new genome #####
################################

###get software
git clone https://github.com/KrisChristensen/MapVCF2NewGenome.git

###Get fasta file of sequences (modified script so I didn't need to get the reference allele which was available from the mareyMap):

$ python3 MapVCF2NewGenome/VCF_2_Fasta_v1.3.py -vcf Order.Okis.Michelle.Filt.LOD.15.size.90.mask.1.LOD.10.LODD.4.SexAveraged.1.vcf -fasta ../genome/GCF_002021735.1_Okis_V1_genomic.fna -fai ../genome/GCF_002021735.1_Okis_V1_genomic.fna.fai > GeneticMapMarkers.all.22Dec2022.fasta

###Align the sequences:

$ bwa mem -M ../genome/GCF_002021735.2_Okis_V2_genomic.wMito.wsdY.fna GeneticMapMarkers.all.22Dec2022.fasta | samtools sort -o GCF_002021735.2.vs.GeneticMapMarkers.all.22Dec2022.bam

###Index files:

$ samtools index GCF_002021735.2.vs.GeneticMapMarkers.all.22Dec2022.bam

###Return new positions:
$ python3 MapVCF2NewGenome/Bam2SNPPosition.v1.1.py -file GCF_002021735.2.vs.GeneticMapMarkers.all.22Dec2022.bam -qual 40 -multiple 1 -minLen 100 -minPid 95 -pos 51 > Old2New.positions.txt

###Filter new positions:
$ python3 MapVCF2NewGenome/CheckMappingVCF.v1.0.py -map Old2New.positions.txt -fasta1 ../genome/GCF_002021735.1_Okis_V1_genomic.fna -fai1 ../genome/GCF_002021735.1_Okis_V1_genomic.fna.fai -fasta2 ../genome/GCF_002021735.2_Okis_V2_genomic.wMito.wsdY.fna -fai2 ../genome/GCF_002021735.2_Okis_V2_genomic.wMito.wsdY.fna.fai -flanking 10 -output most > Old2New.positions.filtered.txt

###List positions for vcftools
kchristensen@woodstock:/scratch/kchristensen/okis/BCSRIF/GATK/truth$ awk '{print $3, $4}' Old2New.positions.filtered.txt > Old2New.positions.filtered.vcftools.txt

###Subset the vcf file for those in the position file:
$ vcftools --gzvcf ../vcf/variants.GATK.iteration.1.Full.sorted.vcf.gz --recode --recode-INFO-all --out variants.GATK.iteration.1.Full.sorted.truth --positions Old2New.positions.filtered.vcftools.txt

	After filtering, kept 90703 out of a possible 49558651 Sites
	Run Time = 1966.00 seconds

$ bgzip variants.GATK.iteration.1.Full.sorted.truth.recode.vcf
$ tabix variants.GATK.iteration.1.Full.sorted.truth.recode.vcf.gz

###Actual variant recalibration ###

$ java -Xms5g -Xmx176g -jar GenomeAnalysisTK.jar -T VariantRecalibrator -R genome/GCF_002021735.2_Okis_V2_genomic.wMito.wsdY.fna -input vcf/variants.GATK.iteration.1.Full.sorted.vcf.gz -recalFile recal/variants.GATK.iteration.1.Full.recal -tranchesFile tranches/variants.GATK.iteration.1.Full.tranches -rscriptFile rFiles/variants.GATK.iteration.1.Full.R -resource:GeneticMaps,known=false,training=true,truth=true,prior=15.0 truth/variants.GATK.iteration.1.Full.sorted.truth.recode.vcf.gz -an QD -an MQ -an MQRankSum -an ReadPosRankSum -an FS -an SOR -an DP -an InbreedingCoeff -mode SNP

### Apply recalibration ###################################################

$ java -Xms5g -Xmx176g -jar GenomeAnalysisTK.jar -T ApplyRecalibration -R genome/GCF_002021735.2_Okis_V2_genomic.wMito.wsdY.fna -input vcf/variants.GATK.iteration.1.Full.sorted.vcf.gz -recalFile recal/variants.GATK.iteration.1.Full.recal -tranchesFile tranches/variants.GATK.iteration.1.Full.tranches -mode SNP -o vcf/variants.GATK.iteration.1.Full.sorted.recalibrated.vcf --ts_filter_level 99.5
