## Supplementary material for "Revealing the evolutionary history and contemporary population structure of Pacific salmon in the Fraser River through genome resequencing": File S2: readme.txt

#########################################################################################
#### Truth variants (Starting 10040 unique variants among studies)   ####################
#########################################################################################

####Combined truth datasets from "The sockeye salmon genome, transcriptome and analyses identifying population defining regions of the genome"
####Originally from: 
####Genomic signatures among Oncorhynchus nerka ecotypes to inform conservation and management of endangered Sockeye Salmon.
####Genomic islands of divergence linked to ecotypic variation in sockeye salmon.
####Genomic Changes Associated with Reproductive and Migratory Ecotypes in Sockeye Salmon (Oncorhynchus nerka).:
####This was done by pasting all variants from the previous vcf files:

	Larson2016.primary.softfiltered.manualFiltered.truth.vcf
	Nichols2016.primary.softfiltered.manualFiltered.truth.vcf
	Veale2017.primary.softfiltered.manualFiltered.truth.vcf

###Combine them

	truthDatasetCombined08Aug2022.vcf
	
###Then using the scripts from github

	### get program
	git clone https://github.com/KrisChristensen/MapVCF2NewGenome.git
	
	### Get Fasta
	$ python MapVCF2NewGenome/VCF_2_Fasta_v1.1.py \
	-vcf truthDatasetCombined08Aug2022.vcf \
	-fasta /scratch2/shared/NCBI.Genomes/GCF_006149115.1_Oner_1.0_genomic.fna \
	-fai /scratch2/shared/NCBI.Genomes/GCF_006149115.1_Oner_1.0_genomic.fna.fai \
	-flanking 100 > \
	truthDatasetCombined08Aug2022.flanking100.fasta

	Finished reading vcf file: Found 10040 variants, used 10019
	
	### Align to new genome
	$ bwa mem -M \
	Oner2021.flye.racon.pilon2.v1.mitoRemoved.min200.mitoAdded.fasta \
	truthDatasetCombined08Aug2022.flanking100.fasta \
	| samtools sort \
	-o truthDatasetCombined08Aug2022.flanking100.vs.V2Genome.bam
	
	$ samtools index truthDatasetCombined08Aug2022.flanking100.vs.V2Genome.bam
	
	### Find new positions
	$ python MapVCF2NewGenome/Bam2SNPPosition.v1.1.py \
	-file truthDatasetCombined08Aug2022.flanking100.vs.V2Genome.bam -qual 60 -multiple 1 -minLen 200 -minPid 99 -pos 101 > old2new.positions.txt
	
	$ python CheckMappingVCF.v1.0.py \
	-map old2new.positions.txt \
	-fasta1 GCF_006149115.1_Oner_1.0_genomic.fna \
	-fai1 GCF_006149115.1_Oner_1.0_genomic.fna.fai \
	-fasta2 Oner2021.flye.racon.pilon2.v1.mitoRemoved.min200.mitoAdded.fasta \
	-fai2 Oner2021.flye.racon.pilon2.v1.mitoRemoved.min200.mitoAdded.fasta.fai \
	-flanking 20 \
	-verbose yes \
	-output most > old2new.positions.filtered.flank20.most.txt
	
		Finished reading vcf file: Found 8922 variants, correct variants:6582, unknown variants:0, CompleteMismatch:91, variantOnlyMismatch:1953, partialMatch:296
	
	### Had to modify the script because the original didn't have a header #######
		
	python AssignNewSNPPositions.v1.3.truth.py \
	-map old2new.positions.filtered.flank20.most.txt \
	-vcf truthDatasetCombined08Aug2022.vcf \
	-gen Oner2021.flye.racon.pilon2.v1.mitoRemoved.min200.mitoAdded.fasta \
	-fai Oner2021.flye.racon.pilon2.v1.mitoRemoved.min200.mitoAdded.fasta.fai \
	-header yes > truthDatasetCombined09Aug2022.map2V2.vcf

	
	### Find the intersection ##############################
	
	$ vcf-isec \
	-n =2 \
	-f variants.GATK.iteration.1.Full.sorted.vcf.gz \
	truthDatasetCombined09Aug2022.map2V2.sorted.vcf.gz | \
	bgzip -c > truthDatasetCombined09Aug2022.vs.full.intersection.vcf.gz
	
	
	###########################################################################
	### Variant recalibration #################################################
	###########################################################################
	
	$ java -Xms5g -Xmx176g -jar GenomeAnalysisTK.jar \
	-T VariantRecalibrator \
	-R Oner2021.flye.racon.pilon2.v1.mitoRemoved.min200.mitoAdded.fasta \
	-input variants.GATK.iteration.1.Full.sorted.vcf.gz \
	-recalFile recal/variants.GATK.iteration.1.Full.recal \
	-tranchesFile tranches/variants.GATK.iteration.1.Full.tranches \
	-rscriptFile rFiles/variants.GATK.iteration.1.Full.R \
	-resource:GeneticMaps,known=false,training=true,truth=true,prior=15.0 truthDatasetCombined09Aug2022.vs.full.intersection.vcf.gz \
	-an QD -an MQ -an MQRankSum -an ReadPosRankSum -an FS -an SOR -an DP -an InbreedingCoeff -mode SNP
	
	###########################################################################
	### Apply recalibration ###################################################
	###########################################################################
	
	$ java -Xms5g -Xmx176g -jar GenomeAnalysisTK.jar \
	-T ApplyRecalibration \
	-R Oner2021.flye.racon.pilon2.v1.mitoRemoved.min200.mitoAdded.fasta \
	-input variants.GATK.iteration.1.Full.sorted.vcf.gz \
	-recalFile recal/variants.GATK.iteration.1.Full.recal \
	-tranchesFile tranches/variants.GATK.iteration.1.Full.tranches \
	-mode SNP \
	-o variants.GATK.iteration.1.Full.sorted.recalibrated.vcf \
	--ts_filter_level 99.5
	
